# Apoplastic CBM1-interacting proteins bind conserved carbohydrate binding module 1 motifs in fungal hydrolases to counter pathogen invasion

**DOI:** 10.1101/2021.12.31.474618

**Authors:** Takumi Takeda, Machiko Takahashi, Motoki Shimizu, Yu Sugihara, Hiromasa Saitoh, Koki Fujisaki, Kazuya Ishikawa, Hiroe Utsushi, Eiko Kanzaki, Yuichi Sakamoto, Akira Abe, Ryohei Terauchi

## Abstract

When infecting plants, fungal pathogens secrete cell wall degrading enzymes (CWDEs) that break down cellulose and hemicellulose, the primary components of plant cell walls. Some fungal CWDEs contain a unique domain, named the carbohydrate binding module (CBM), that facilitates their access to polysaccharides. However, little is known about how plants counteract pathogen degradation of their cell walls. Here, we show that the rice cysteine-rich repeat secretion protein OsCBMIP binds to and inhibits xylanase MoCel10A of the blast fungus pathogen *Magnaporthe oryzae*, interfering with its access to the rice cell wall and degradation of rice xylan. We found binding of OsCBMIP to various CBM1-containing enzymes, suggesting it has a general role in inhibiting the catalytic activities of fungal enzymes. OsCBMIP is localized to the apoplast, and its expression is strongly induced in leaves infected with *M. oryzae*. Remarkably, knockdown of *OsCBMIP* reduced rice defense against *M. oryzae*, demonstrating that inhibition of CBM1-containing fungal enzymes by OsCBMIP is crucial for rice defense. We also identified additional CBMIP-related proteins from *Arabidopsis thaliana* and *Setaria italica*, indicating that a wide range of plants counteract pathogens through this mechanism.

**Summary:** Plants have evolved various activity-inhibiting proteins as a defense against fungal cell wall degrading enzymes (CWDEs), but how plants counteract the function of fungal enzymes containing carbohydrate binding modules (CBMs) remains unknown. Here, we demonstrate that OsCBMIP, a member of the cysteine-rich repeat secretion protein family, interacts with fungal CBM1. OsCBMIP binding to CBM1 of a blast fungal xylanase blocks access to cellulose, resulting in the inhibition of xylanase enzymatic activity. Our study provides significant insights into plant countermeasure against CWDEs in the apoplastic space during plant–fungal pathogen interactions. It also reveals a molecular function of the DUF26 domain widely distributed in plant proteins.

## Introduction

Plant pathogens have developed various strategies to infect and exploit host plants. In response, plants have evolved a complex and multi-layered immune system to overcome invading pathogens (Jones and Dangl 2006; Spoel and Dong 2012). Many plant pathogens invade via the apoplast, or extracellular space, of host plants. To this end, fungal pathogens secrete various effector molecules into the host apoplastic space, including hydrolytic enzymes that degrade plant cell wall polysaccharides (Walton 1994; Stergiopoulos and de Wit 2009; Jashni et al. 2015). Plants counter the pathogens by secreting proteases, hydrolytic enzymes and activity-inhibiting proteins, and also by inducing immune responses after recognition of pathogen-derived molecules (Juge 2006; van der Hoorn 2008; Kim et al. 2009; Valueva and Mosolov 2014;). Therefore, study of apoplastic molecules that play key roles in plant pathogen invasion and host immunity is essential for understanding host– pathogen interactions.

Plant apoplastic space is filled with primary cell wall, mainly composed of polysaccharides including cellulose, hemicellulose and pectin. Hemicellulosic polysaccharides play an important role in controlling the physical properties of the cell wall. Xyloglucan in dicotyledonous and xylan in monocotyledonous plants are the major hemicellulosic polysaccharides by quantity and strengthen the cell wall by forming cross-bridges between cellulose microfibrils (Takeda et. al. 2002; Takahashi et al. 2014). A cell wall composed of heteropolysaccharides also provides a physical barrier against plant pathogen invasion (Vorwerk et al. 2004).

Plant pathogenic fungi secrete a battery of cell wall degrading enzymes (CWDEs) that catalyze hydrolytic and oxidative degradation of plant cell wall polysaccharides, assisting fungal penetration and colonization. Sugars released from degraded cell walls serve as a carbon source for the pathogens. Notably, some CWDEs possess a carbohydrate binding module (CBM) connected to the catalytic core domain by a linker peptide. CBMs are classified into 88 subgroups (CAZy, http://www.cazy.org), of which CBM family 1 (CBM1) specifically binds to cellulose and is found only in fungal enzymes (Kraulis et al. 1989; Mattinen et al. 1997; Lehtiö et al. 2003). Critical roles of CBMs in the CWDE-mediated hydrolysis of water-insoluble substrates have been demonstrated. CBM-truncated versions of CWDEs show reduced hydrolytic activities, whereas addition of an extra CBM enhances hydrolytic activities compared with those of wild-type proteins (Hägglund et al. 2003; Ito et al. 2004; Takahashi et al. 2010). Direct application of a CBM1-containing xylanase (MoCel10A) from *Magnaporthe oryzae* to wheat coleoptile segments reduced cell wall strength to a greater degree than xylanase lacking CBM1 (Takahashi et al. 2014). These observations highlight the role of CBM-containing CWDEs in plant cell wall degradation. Plants counter this enzymatic hydrolysis by pathogens by producing proteins that inhibit enzyme activities of xyloglucan-specific endoglucanase, xylanase and polygalacturonase (York et al., 2004; Di et al. 2006; Jones and Dangl 2006; Dornez et al. 2010; Maulik et al. 2012). However, plant proteins that inhibit CBM function have not yet been reported.

Cysteine-rich repeat secretion proteins (CRRSPs) are widespread in land plants and consist of a secretion signal sequence and one or more repeats of the domain of unknown function 26 (DUF26, PF01657) containing a cysteine-rich repeat motif (CRR motif, C-X8-C-X2-C). Ginkbilobin-2 (Gnk2), found in the endosperm of *Ginkgo biloba* seed, is an extracellular CRRSP containing a single DUF26 domain with antifungal activity (Sawano et al. 2007). Biochemical and structural analyses of Gnk2 revealed a mannose-binding protein composed of two α-helices and a five-stranded β-sheet (Miyakawa et al. 2009 and 2014), while mutant studies showed a correlation between Gnk2 mannose-binding and antifungal activity (Miyakawa et al. 2014). A jasmonic acid (JA)-induced rice protein, OsRMC, is reported to contain two DUF26 domains and be involved in root meander curling and salt stress responses; however, its biochemical function has not been determined (Jiang et al. 2007; Zhang et al. 2009). Ma et al. (2018) reported that the apoplastic effector Rsp3 of the smut fungus *Ustilago maydis* binds maize (*Zea mays*) extracellular DUF26-containing proteins AFP1 and AFP2. AFP1 binds mannose and exhibits antifungal activity against *U. maydis*, which is blocked by the Rsp3 effector. Virus-mediated gene silencing of *AFP1* and *AFP2* in maize enhances the virulence of *U. maydis*, suggesting that AFP1 and AFP2 may function in host defense. However, the molecular mechanism of AFP1 and AFP2 antifungal activity has not been elucidated. Cysteine-rich repeat kinases (CRKs), members of the receptor-like kinase (RLK) family, are composed of an extracellular domain with two DUF26 repeats, a transmembrane domain, and a C-terminal cytoplasmic Ser/Thr kinase domain. Forty-four CRK genes are known in the *Arabidopsis thaliana* genome, representing one of the largest groups of RLKs (Wrzaczek et al. 2010). CRKs are induced by reactive oxygen species (ROS) (Czernic et al. 1999) and salicylic acid (Czernic et al. 1999; Ohtake et al. 2000) and are involved in defense responses against pathogens (Chen et al. 2003; Acharya et al. 2007; Yeh et al. 2015; Chern et al. 2016; Yadeta et al. 2017; Du et al. 2019). CRKs are hypothesized to be involved in ROS/redox signaling and sensing, mediated by the conserved cysteine residues in the DUF26 domains (Bourdais et al. 2015; Yadeta 2017). Plasmodesmata-localized proteins (PDLPs), composed of two DUF26 domains in their extracellular region and a transmembrane domain, are involved in cytoplasmic signaling (Lim et al. 2016; Brunkard and Zambryski 2017), pathogen response and control of callose deposition (Caillaud et al. 2014; Cui and Lee 2016). Despite an increasing number of studies of CRKs and PDLPs, the molecular function of their DUF26 domains remains elusive. A recent amino acid sequence comparison of DUF26-containing proteins revealed the possible evolutionary history of this protein family (Vaattovaara et al. 2019). CRKs consisting of an extracellular DUF26 domain fused with a transmembrane domain and cytosolic kinase domain might have evolved from an ancestral protein with a single DUF26 domain. After the emergence of the ancestral CRK, the DUF26 domain duplicated, resulting in CRKs with two DUF26 domains. CRRSPs and PDLPs most likely originated from CRKs by deletion of both the transmembrane domain and kinase domain (CRRSPs) or just the kinase domain (PDLPs). The first and second DUF26 domains (DUF26-A and DUF26-B) in CRRSPs are phylogenetically distinct (Vaattovaara et al. 2019).

Here, we report on a CBM1-interacting protein (OsCBMIP) of rice (*Oryza sativa*). OsCBMIP is a member of the CRRSPs containing two DUF26 domains that binds *M. oryzae* CBM1-containing CWDEs. OsCBMIP inhibits cell wall degrading activity of fungal enzymes and plays an important role in rice defense against *M. oryzae* infection. We also show that CBM1-binding CRRSPs proteins are widespread among plant species. Our findings provide insight into the apoplastic molecular interactions between pathogen cell wall degrading enzymes and plant proteins containing DUF26 domains.

## Results

### Rice OsCBMIP, a CRRSP, binds CBM1 and inhibits blast pathogen xylanase activity

To identify rice proteins that interact with CBM1, we incubated protein extract prepared from rice leaves 4 days after *M. oryzae* inoculation with a His-tagged xylanase protein from *M. oryzae* (MoCel10A-His), which contains CBM1 and belongs to glycoside hydrolase family 10 (Takahashi et al. 2014). The fraction bound to His-tag affinity resin (His-resin) was analyzed by SDS-PAGE and protein staining. We detected a 25 kDa protein band from the mixture of rice protein extract and MoCel10A-His but not from the rice protein extract alone (Fig. 1*A*). Liquid chromatograph–tandem mass spectrometry (LC-MS/MS) analysis of the 25 kDa band revealed that the protein was a product of *OsRMC* (Jiang et al. 2007; Zhang et al. 2009), encoding a protein with an N-terminal secretion signal peptide and a pair of DUF26 domains (Fig. 1*A* and *SI Appendix*, Fig. S1). We generated rice suspension-cultured cells expressing the full-length *OsRMC* gene fused with a His-tag at its C-terminus driven by the maize ubiquitin promoter. Secreted OsRMC-His protein was mixed with Flag-tagged MoCel10A (MoCel10A-Flag) or CBM1-truncated MoCel10A (MoCel10AΔCBM-Flag), and the protein mixtures were separated into bound and unbound fractions using His-resin. We detected MoCel10A-Flag in the bound fraction and MoCel10AΔCBM-Flag in the unbound fraction (Fig. 1*B*). From this result, we concluded that OsRMC binds to the CBM1 domain of MoCel10A. We hereafter rename OsRMC as *O. sativa* CBM1-interacting protein (OsCBMIP) on the basis of its binding specificity. Gel-permeation chromatography of the mixture of OsCBMIP-His and MoCel10A-His confirmed that these proteins form a complex, in which the two proteins were detected at higher molecular weight positions compared with the individual molecules (*SI Appendix*, Fig. S2).

**Fig. 1.**
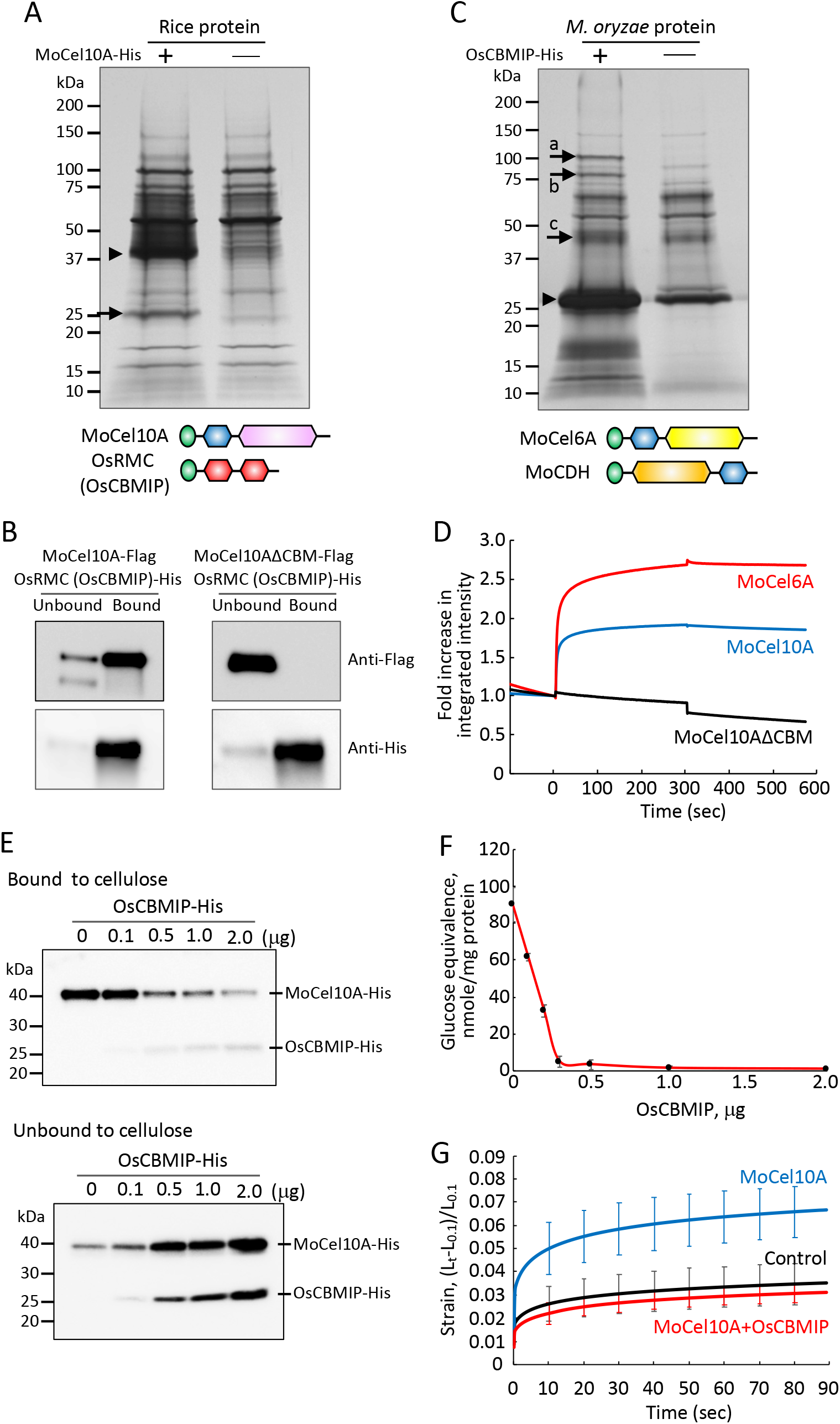
OsCBMIP binds to CBM1 of fungal enzymes. (A) Protein extract from rice leaves 4 days after inoculation with *M. oryzae* (Ken53-33) was incubated with (+) or without (-) MoCel10A-His in the presence of His-resin. Fractions bound to His-resin were subjected to SDS-PAGE followed by silver staining. Arrowhead indicates MoCel10A-His. The protein band indicated by an arrow was identified by LC-MS/MS. Schematic structures of predicted full-length proteins of MoCel10A and OsRMC (OsCBMIP) are shown: green, secretion signal peptide; blue, CBM1 (PF00734); pink, GH10 catalytic core domain (PF00331); red, DUF26 domain (PF01657). (B) OsCBMIP-His mixed with MoCel10A-Flag or MoCel10AΔCBM-Flag was applied to His-resin. Fractions unbound and bound to His-resin were subjected to SDS-PAGE followed by immunoblot analysis using anti-Flag and anti-His antibodies. (C) Culture filtrate from *M. oryzae* hyphae liquid culture was incubated with (+) or without (-) OsCBMIP-His in the presence of His-resin. Fractions bound to His-resin were subjected to SDS-PAGE followed by silver staining. Arrowhead and arrows indicate OsCBMIP-His and candidate proteins bound to OsCBMIP-His, respectively. Schematic structures of MoCel6A and OsCDH are shown: green, secretion signal peptide; blue, CBM1 (PF00734); yellow, GH6 catalytic core domain (PF01341); orange, CDH-cyt domain (PF16010). (D) Integrated intensity representing the binding kinetics of OsCBMIP to MoCel10A, MoCel6A and MoCel10AΔCBM. Association of MoCel10A, MoCel6A and MoCel10AΔCBM (10 µL, 2.0 mM), respectively, with OsCBMIP was measured from time 0 to 300 s. Binding buffer was used to measure protein dissociation from time 300 to 570 s. Fold increase in integrated intensity was calculated by dividing each trajectory by the value at time zero. Data are means ± SD of three independent determinations. (E) MoCel10A-His was incubated with cellulose in the presence of OsCBMIP-His (0–2.0 µg) for 1 h at 4 °C, and proteins bound and unbound to cellulose were detected by immunoblot analysis using an anti-His antibody. (F) MoCel10A-His (2.0 µg) was preincubated with OsCBMIP-His (0–2.0 µg) for 30 min at 4 °C, and the hydrolytic activity towards a wheat cell wall preparation was determined. Data are means ± SD of three independent determinations. (G) Heat-inactivated wheat coleoptile segments were treated with sodium phosphate buffer (100 mM, pH 6.0) containing MoCel10A-His (2.0 µg) with BSA (2.0 µg) or a mixture of MoCel10A-His and OsCBMIP-His (2.0 µg each). Samples treated with buffer were used as a control. Extension measurement was performed by loading a constant 200 mN to approximately 5 mm segments for 3 min in an extensometer. Strain was calculated as (*L*_*t*_ – *L*_0.1_)/*L*_0.1_ (*L*_*t*_, length of coleoptiles at each time point; *L*_0.1_, length of segments at time 0.1). Data are means ± SD of five independent determinations.

To isolate additional secreted proteins of *M. oryzae* that interact with OsCBMIP, we carried out a pull-down experiment by incubating OsCBMIP-His with *M. oryzae* culture filtrate followed by recovery of interacting proteins using His-resin (Fig. 1*C*). Protein bands detected from the mixture of OsCBMIP-His and *M. oryzae* culture filtrate, but not from the *M. oryzae* culture filtrate alone, were identified as (a) cellobiose dehydrogenase (MoCDH), (b) cellobiohydrolase (MoCel6A) and (c) xylanase (MoCel10A). In all cases, the predicted full-length proteins consisted of a secretion signal peptide, CBM1 and a catalytic core domain. We confirmed the interaction of OsCBMIP-His with Flag-tagged MoCDH and MoCel6A by pull-down assay (*SI Appendix*, Fig. S3). Furthermore, when we carried out pull-down assays using OsCBMIP-His and the culture filtrate of *Trichoderma reesei*, an ascomycete fungus, we identified six proteins possessing CBM1 in the OsCBMIP-binding fraction (*SI Appendix*, Fig. S4). These results indicate that OsCBMIP binds to CBM1 of various fungal enzymes.

We then measured the binding kinetics of OsCBMIP to MoCel10A and MoCel6A, both harboring N-terminal CBM1 (Fig. 1*D*). Binding of OsCBMIP to MoCel10A and MoCel6A occurred rapidly, and the resulting complexes were stable. Binding kinetics values of OsCBMIP with MoCel10A and MoCel6A, respectively, were as follows: *K*_a_, 2.98±0.87 and 3.89±0.51 (10^4^ M^-1^ s^-1^); *K*_d_, 1.36±0.42 and 8.45±1.39 (10^−4^ s^-1^); *K*_D_, 4.60±0.90 and 2.19±0.31 (10^−8^ M). Binding of OsCBMIP to MoCel10AΔCBM was not detected. These results suggest that OsCBMIP binds CBM1 quickly and the resulting complexes are stable.

CBM1 is known to tightly bind cellulose (Kraulis et al. 1989; Mattinen et al. 1997; Lehtiö et al. 2003). To investigate the effect of OsCBMIP on binding of CBM1 to cellulose, we mixed MoCel10A-His with variable amounts of OsCBMIP-His prior to incubation with cellulose, and then detected the resulting cellulose-bound and -unbound MoCel10A-His (40 kDa) as well as OsCBMIP-His (25 kDa) proteins using an anti-His antibody (Fig. 1*E*). The more OsCBMIP-His added, the less MoCel10A-His bound to cellulose, indicating that OsCBMIP interfered with MoCel10A binding to cellulose. We also tested the catalytic activity of MoCel10A-His for xylan hydrolysis in the presence of OsCBMIP-His using a water-insoluble wheat coleoptile cell wall preparation. The hydrolytic activity of MoCel10A decreased with the increase in OsCBMIP-His content (Fig. 1*F*). However, MoCel10A hydrolytic activity towards water-soluble xylan was not affected by OsCBMIP (*SI Appendix*, Fig. S5). These results agree with our previous finding that the presence of CBM1 facilitates enzymatic hydrolysis of water-insoluble polysaccharides (Takahashi et al. 2014).

Glucurono-arabinoxylan is a major hemicellulosic polysaccharide of primary cell walls in monocotyledonous plants and is thought to provide cell wall rigidity by forming cross-bridges between cellulose microfibrils. We therefore examined the effects of MoCel10A-His and OsCBMIP-His on the extensibility of heat-inactivated wheat coleoptile segments (Fig. 1*G*). Treatment with MoCel10A-His enhanced the strain on wheat coleoptile segments, implying a reduction in cell wall strength through the action of MoCel10A. However, addition of OsCBMIP-His completely blocked the effect of MoCel10A-His. This result indicates that binding of OsCBMIP to CBM1 inhibits xylan hydrolysis, resulting in suppressed extensibility of coleoptile segments.

### Apoplast-localized OsCBMIP contributes to rice defense against *M. oryzae* infection by CBM1 binding

To observe the function of OsCBMIP *in planta*, we transiently expressed OsCBMIP fused to GFP (OsCBMIP-GFP) in *Nicotiana benthamiana* leaves by agroinfiltration and observed its subcellular localization. We detected GFP signal at the outer periphery of cells when OsCBMIP-GFP was expressed, whereas we observed GFP control signal throughout the cells (Fig. 2*A*). Protein fractionation revealed OsCBMIP-GFP in the buffer-soluble fraction but not the membrane fraction of *N. benthamiana* leaves (*SI Appendix*, Fig. S6). In addition, OsCBMIP with C-terminal hemagglutinin epitope tag (HA-tag) overexpressed in rice suspension cells was secreted into the culture medium (*SI Appendix*, Fig. S6). These results indicate that OsCBMIP is present in the apoplastic space and is not anchored on cellular membranes.

**Fig. 2.**
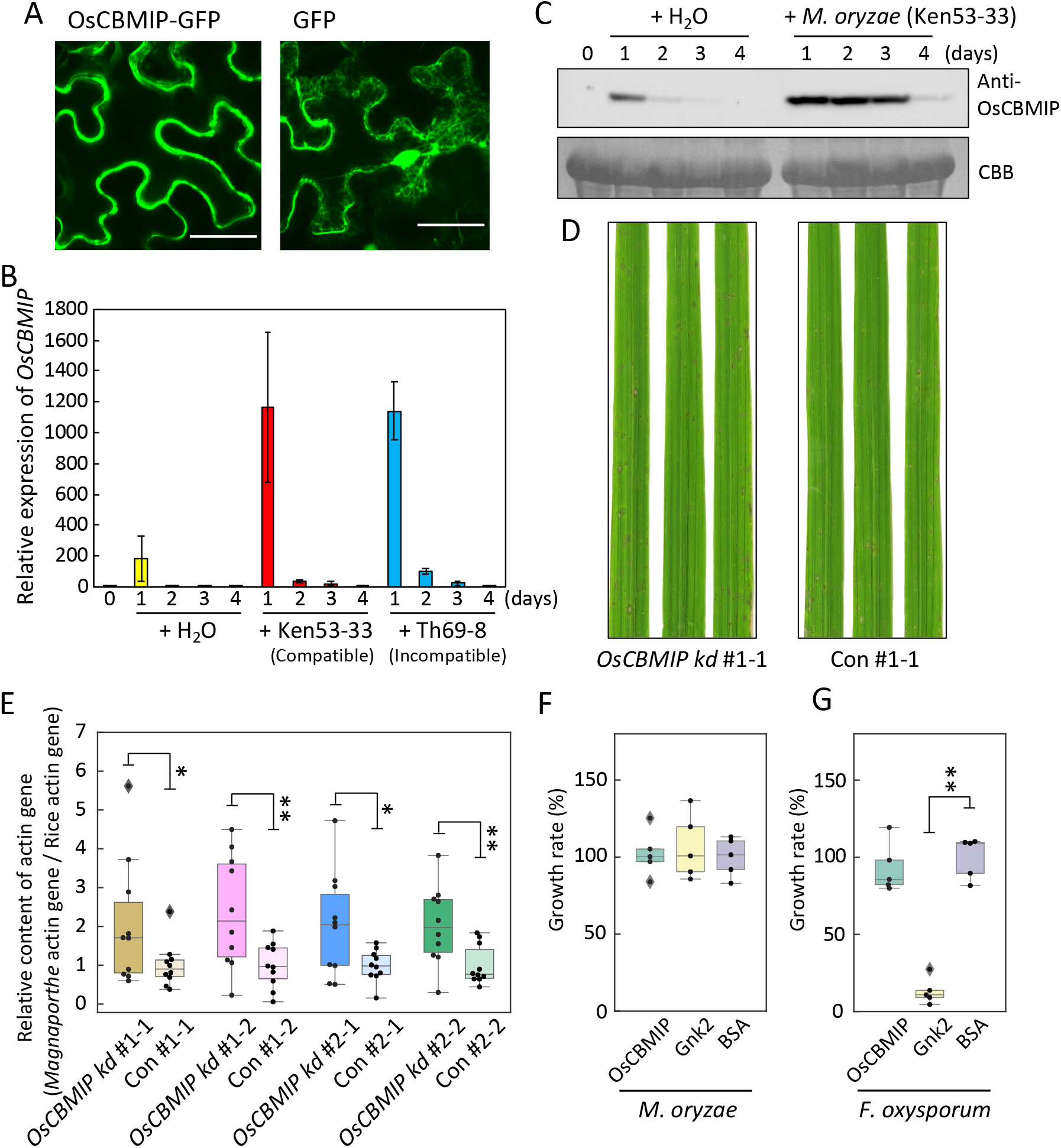
OsCBMIP is involved in rice defense against *M. oryzae* infection. (A) OsCBMIP localized to apoplast. Subcellular localization of OsCBMIP was determined using *N. benthamiana* leaves overexpressing OsCBMIP-GFP or GFP as control by UV-fluorescence microscopy. Bars, 10 nm. (B) *OsCBMIP* expression induced by *M. oryzae* infection in both compatible and incompatible interactions. Expression levels of *OsCBMIP* transcripts in rice (‘Hitomebore’) leaves after inoculation with *M. oryzae* (Ken53-33, compatible strain; TH69-8, incompatible strain) were determined by qRT-PCR. *OsCBMIP* expression level was normalized using that of the rice ubiquitin gene (*LOC_Os03g03920*.*1*). Data are means ± SD of three independent determinations. (C) OsCBMIP protein expression is induced after *M. oryzae* infection. Protein extracts (20 mg) from rice leaves 0–4 days after *M. oryzae* (Ken53-33) inoculation were subjected to immunoblot analysis using an anti-OsCBMIP antibody. Equal protein loading on SDS-PAGE was verified by Coomassie Brilliant Blue (CBB) staining. (D) Rice leaves of *OsCBMIP*-knockdown and wild-type control lines 4 days after *M. oryzae* (Ken53-33) inoculation. The *M. oryzae* infection test was carried out using T_1_ *OsCBMIP*-knockdown (*kd*) and wild-type control (Con) progenies (*n*=10) derived from selfing of T_0_ heterozygous *OsCBMIP*-knockdown lines. (E) *M. oryzae* infection is enhanced in *OsCBMIP* knockdown rice. The amount of *M. oryzae* fungal mass in rice leaf was monitored by quantifying the ratio of *M. oryzae* genomic DNA to rice genomic DNA, determined by qPCR of the respective *actin* genes. The average ΔΔCt value in wild-type control lines was defined as the unit for the ratio. Data are means ± SD of 10 independent determinations. Single and double asterisks indicate a significant difference at *P* < 0.05 and *P* < 0.01, respectively, according to Student *t*-test. (F) Antifungal activity of OsCBMIP and Gnk2 against *M. oryzae* and *F. oxysporum*. Cells of *M. oryzae* overexpressing luciferase or of *F. oxysporum* were cultured in the presence of OsCBMIP (10 µM, 79 µg/300 mL culture medium) supplemented with 21 µg of BSA, Gnk2 (10 µM, 38 µg/300 µL culture medium) supplemented with 62 µg of BSA, or BSA (100 µg) alone for 36 h at 25 °C. Growth of *M. oryzae* and *F. oxysporum* was evaluated by measuring luciferase activity and optical density at 600 nm, respectively. Data are means ± SD of five independent determinations.

We investigated the expression of *OsCBMIP* in rice leaves inoculated with *M. oryzae* using quantitative reverse-transcription PCR (qRT-PCR) (Fig. 2*B*). The basal level of *OsCBMIP* expression (at time zero) in leaves was very low. Gene expression was slightly induced 1 day after water treatment as control; by contrast, expression levels were more than fivefold higher than in the control 1 day after inoculation with *M. oryzae* isolates Ken53-33 (compatible) or Th69-8 (incompatible). Expression was then rapidly downregulated 2 days after *M. oryzae* inoculation. We also evaluated OsCBMIP protein accumulation in rice leaves using immunoblot analysis with anti-OsCBMIP antibody (Fig. 2*C*). OsCBMIP protein was undetectable at time zero and slightly induced at 1 day after water treatment. By contrast, OsCBMIP accumulated highly at 1–3 days after inoculation with compatible *M. oryzae* isolate Ken53-33 and then decreased at 4 days after inoculation. These results suggest that gene expression of *OsCBMIP* is markedly induced by *M. oryzae* inoculation and a high level of OsCBMIP protein accumulation is maintained for several days.

To investigate the role of OsCBMIP in rice immunity to *M. oryzae*, we performed knockdown of the *OsCBMIP* gene in rice cultivar ‘Moukoto’ by RNA interference (RNAi)-mediated gene silencing. We generated transgenic lines (#1 and #2) by introducing a gene-silencing vector containing DNA fragments from Open Reading Frame (311 bp) and 3′-UTR (433 bp) regions of the *OsCBMIP* transcript, respectively (*SI Appendix*, Fig. S7). Self-propagation of transgenic rice plants of the T_0_ generation produced the T_1_ generation, which segregated for progeny with and without the RNAi construct. We distinguished *OsCBMIP*-knockdown and *OsCBMIP*-expressing plants by PCR of the hygromycin gene in the RNAi construct and by immunoblotting of OsCBMIP (*SI Appendix*, Fig. S8). OsCBMIP protein accumulation was almost undetectable in *OsCBMIP*-knockdown lines, but was clearly detected in control lines. These T_1_ progeny were spray-inoculated with *M. oryzae* (Ken53-33), which is compatible with the rice cultivar ‘Moukoto’. Four days after *M. oryzae* inoculation, disease lesions caused by *M. oryzae* infection were visible on the inoculated rice leaves (Fig. 2*D*, *SI Appendix*, Fig. S8). More severe disease symptoms were observed in *OsCBMIP*-knockdown lines #1-1, #1-2, #2-1 and #2-2 than in the corresponding controls. We evaluated *M. oryzae* fungal mass in rice leaves by measuring the amount of *M. oryzae* genomic DNA normalized to the amount of rice genomic DNA by qPCR of the *ACTIN* gene (Fig. 2*E*, *SI Appendix*, Fig. S9). The amount of *M. oryzae* fungal mass was significantly higher in leaves of *OsCBMIP*-knockdown lines #1-1, #1-2, #2-1 and #2-2 than in those of the corresponding controls. These results indicate that OsCBMIP plays a role in rice defense against *M. oryzae* infection.

Two CRRSPs, Gnk2 of *G. biloba* (Sawano et al. 2007) and AFP1 of maize (Lay-Sun et al. 2018), bind mannose and show antifungal activities. To address whether the rice blast resistance mediated by OsCBMIP is caused by its antifungal activity, we cultured *M. oryzae* and *Fusarium oxysporum* in the presence of OsCBMIP, Gnk2 or BSA. We observed no difference in the growth of *M. oryzae* among the three treatments (Fig. 2*F*, *Appendix*, Fig. S10). In the case of *F. oxysporum*, there was no difference between the OsCBMIP and BSA treatments at 30 h; however, a significant reduction in growth was observed after the Gnk2 treatment (Fig. 2*G*), as reported previously (Miyakawa et al. 2014), suggesting that whereas Gnk2 exhibits antifungal activity against *F. oxysporum*, OsCBMIP does not possess antifungal activity against *M. oryzae* or *F. oxysporum*. This result indicates that the rice defense conferred by *OsCBMIP* is mediated by OsCBMIP binding to pathogen CBM1 and inhibition of fungal hydrolases, not by direct antifungal activity of the protein.

### A wide range of plant species have CBMIPs

To investigate whether plant species other than rice have CBMIP, we carried out a pull-down assay using MoCel6A-His and protein extract of *Setaria italica* (foxtail millet) leaves inoculated with *M. oryzae* (isolate GFSI 1-7-2) (*SI Appendix*, Fig. S11). We detected a protein band in the mixture of *S, italica* protein and MoCel6A-His but not in the sample with *S. italica* protein alone. This protein was identified as a member of the CRRSPs, named SiCBMIP (XM_004960453.3), which also bound to MoCel10A (Fig. 3*A*). We also searched the Arabidopsis gene database for candidate CBMIPs with amino acid sequence similarity to OsCBMIP and assayed three proteins for their binding to MoCel10A-His (*SI Appendix*, Fig. S12). One of them, encoded by *AT3G22060*.*1*, hereafter named AtCBMIP, showed binding to MoCel10A-His (Fig. 3*A*, *SI Appendix*, Fig. S12). These results suggest that a wide range of plant species possess CBM1-binding CRRSPs. CRKs are composed of an extracellular domain with two or more DUF26 domains, a transmembrane domain and a kinase-like-domain. To test whether CRKs bind CBM1, we produced recombinant proteins comprising the extracellular domains from seven rice CRKs in *N. benthamiana* leaves and assayed their binding to MoCel10A-His. The extracellular domain encoded by *LOC_Os07g35580*.*1*, hereafter named OsCBMIP-K, bound to MoCel10A (Fig. 3A, *SI Appendix*, Fig. S13).

**Fig. 3.**
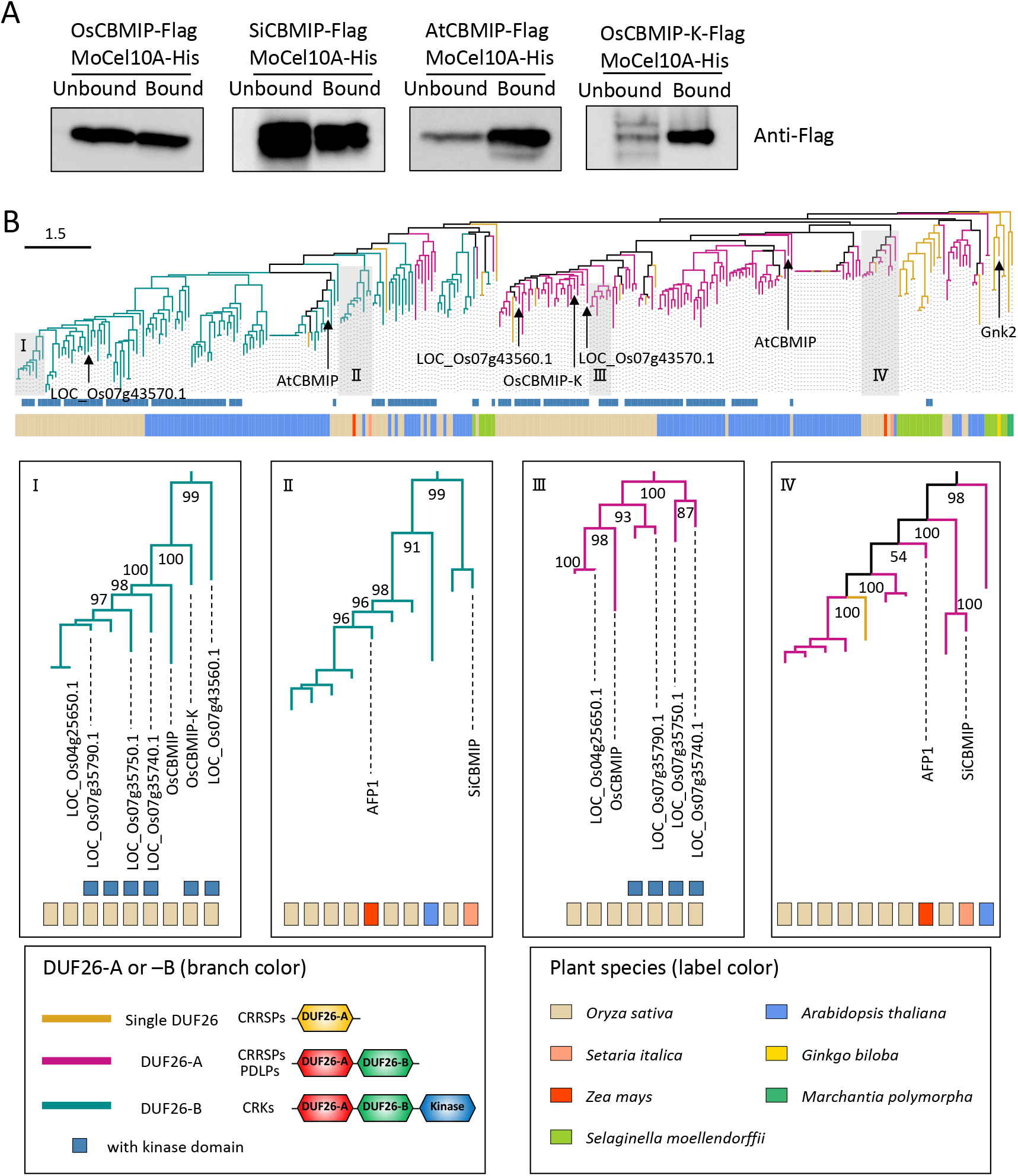
CBM1-binding proteins are widespread in plants. (A) CRRSPs (OsCBMIP, SiCBMIP, AtCBMIP) and OsCBMIP-K (*LOC_Os07g35580*.*1*) bind MoCel10A. Flag-tagged CRRSPs and OsCBMIP-K were incubated with MoCel10A-His in the presence of His-resin. Fractions unbound and bound to His-resin were subjected to immunoblot analysis using an anti-Flag antibody. (B) Phylogenetic tree reconstructed using single DUF26, DUF26-A and DUF26-B domains of CRRSPs and CRKs from *O. sativa, A. thaliana, Selaginella moellendorffii* and *M. polymorpha*, and Gnk2, SiCBMIP and AFP1. Bar, 1.5 amino acid substitutions per site.

To infer the evolution of CRRSP function, we reconstructed a phylogenetic tree of the DUF26 domain using amino acid sequences of CRRSPs and CRKs from *O. sativa* and *A. thaliana* together with CRRSPs of *Selaginella moellendorffii* (Lycopodiophyta) and *Marchantia polymorpha* (Bryophyte), Gnk2 of *G. biloba*, AFP1 of maize and SiCBMIP of *S. italica* (Fig. 3*B*). DUF26 domains in CRRSPs with two such domains were separately treated as DUF26-A (N-terminal) and DUF26-B (C-terminal). In the tree, DUF26-A and DUF26-B form two major clades that diverged from an ancestral singleton DUF26 domain of bryophytes (*M. polymorpha*), as suggested by Vaattovaara et al. (2019). Gnk2 of *G. biloba*, a mannose-binding, single DUF26 domain CRRSP, is located close to the root of the tree. The DUF26-A and DUF26-B domains of maize AFP1, another mannose-binding protein, are positioned close to the roots of their respective clades, which also include the DUF26 domains of SiCBMIP from *S. italica*. The DUF26-A and -B domains of each of the three CBM1-binding CRRSPs, OsCBMIP, AtCBMIP and SiCBMIP, which share relatively low amino acid identities (27–30%) (*SI Appendix*, Fig. S14, Table S1), are on separate branches in the tree, indicating that CBM1-binding CRRSPs may have evolved independently multiple times in these plant lineages. The DUF26 domains from *O. sativa* form large monophyletic groups in the DUF26-A and -B clades, respectively, with CRKs representing the majority of proteins. OsCBMIP and OsCBMIP-K are phylogenetically close, and both bind CBM1 (Fig. 3*A*). These findings indicate that OsCBMIP most likely originated from OsCBMIP-K by deletion of the transmembrane domain and the kinase domain.

### OsCBMIP binds mannose as well as CBM1

Previous studies showed that the two CRRSPs, Gnk2 of *G. biloba* (Sawano et al. 2007; Miyakawa et al. 2014) and AFP1 of maize (Lay-Sun et al. 2018), bind mannose and exhibit antifungal activities. Therefore, we investigated binding of OsCBMIP to mannose using a mannose-agarose binding assay (Fig. 4*A*). Incubation of OsCBMIP with mannose-agarose resulted in recovery of OsCBMIP in the bound fraction. This result indicates that OsCBMIP binds mannose as well as CBM1. The DUF26 domains of OsCBMIP-K, the likely progenitor of OsCBMIP, also bound mannose (*SI Appendix*, Fig. S15). Those of Os07g43560.1, a sister-group CRK of OsCBMIP and OsCBMIP-K, bound mannose but not CBM1 (*SI Appendix*, Fig. S15), whereas that of Os07g43570.1, distantly related to DUF26 domains in clade I, did not bind mannose or CBM1 (*SI Appendix*, Fig. S15).

**Fig. 4.**
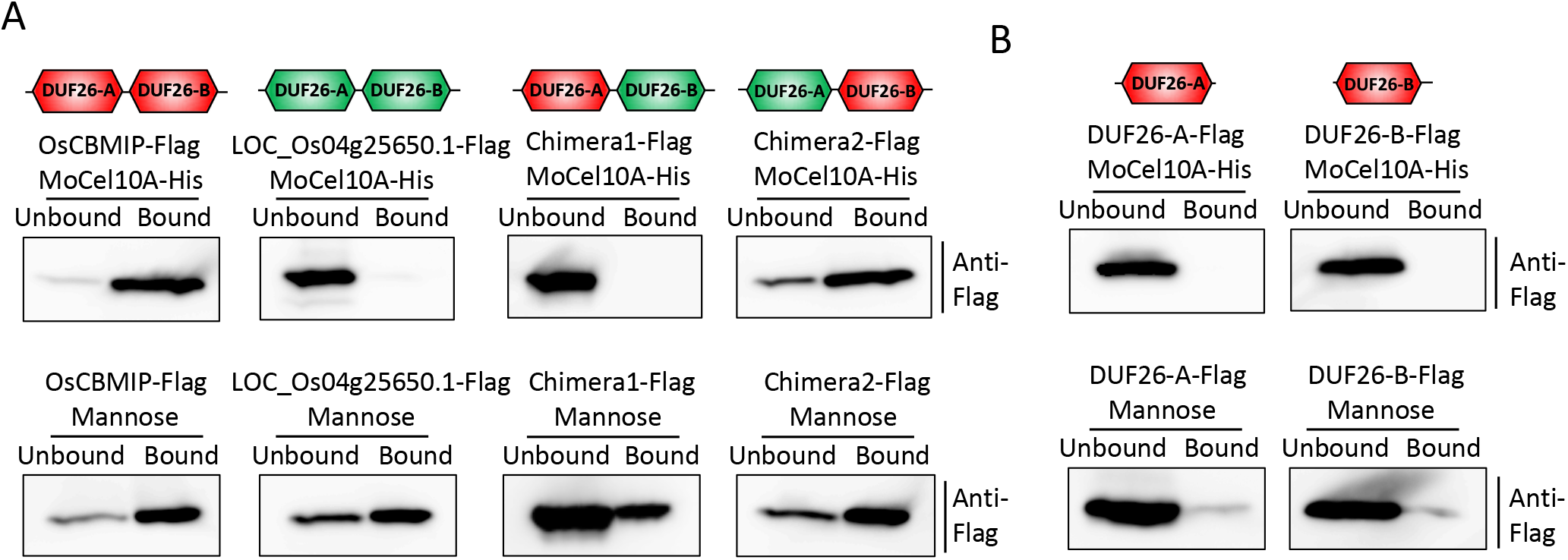
OsCBMIP DUF26-B determines its binding to CBM1. (A) OsCBMIP, LOC_Os04g25650.1 and their chimeric proteins were produced in *N. benthamiana* and assayed for MoCel10A (CBM1: top) and mannose (bottom) binding. Simplified schemes of OsCBMIP (235 amino acids; red) and LOC_Os04g25650.1 (251 amino acids; green) as well as two newly generated chimeric proteins, Chimera1 (244 amino acids, comprising 97 aa [24–120] of OsCBMIP and 147 aa [135–281] of LOC_Os04g25650.1) and Chimera2 (242 amino acids, comprising 104 aa [31–134] of LOC_Os04g25650.1 and 138 aa [121–258] of OsCBMIP) are shown. Flag-tagged proteins were assayed for binding to MoCel10A-His and to mannose. Unbound and bound fractions were subjected to immunoblot analysis with an anti-Flag antibody. (B) Binding assay of single DUF26 domains (DUF26-A and DUF26-B) of OsCBMIP to MoCel10A (CBM1: top) and mannose (bottom).

We also tested another rice CRRSP encoded by *LOC_Os04g25650*.*1*, which showed the highest amino acid sequence identity (59%) to OsCBMIP among rice proteins (Fig. 3*B*, *SI Appendix*, Fig. S14). Our phylogenetic analysis indicated that LOC_Os04g25650.1 is likely derived from OsCBMIP (Fig. 3*B*). This protein bound to mannose but not to MoCel10A (Fig. 4*A*, *SI Appendix*, Fig. S15).

To infer the roles of the DUF26-A and DUF26-B domains in CBM1 and mannose binding, we generated chimeric proteins by exchanging the DUF26-A and DUF26-B domains of OsCBMIP and LOC_Os04g25650.1 (Fig. 4*A*) and tested their binding to the two molecules. Both Chimera1 and Chimera2 bound mannose. Chimera2, with DUF26-A from LOC_Os04g25650.1 and DUF26-B from OsCBMIP, retained CBM1 binding, whereas Chimera1, with DUF26-A from OsCBMIP and DUF26-B from LOC_Os04g25650.1, lost CBM1 binding, suggesting that the DUF26-B domain determines CBM1-binding capability. We hypothesize that the DUF26 domains of the distant progenitor bound an unknown compound, as in the case of Os07g43570.1. After the divergence of clade I DUF26 domains from the rest, some gained the capability to bind mannose, as seen in Os07g43560.1, and subsequently obtained CBM1 binding, as is the case for OsCBMIP-K and OsCBMIP; however, their derivative protein LOC_Os04g25650.1 lost CBM1-binding capability through changes in its DUF26-B domain. A single DUF26 domain of OsCBMIP was unable to bind mannose or CBM1 (Fig. 4*B*), indicating that both DUF26-A and -B are necessary for binding to these two compounds.

Additionally, we tested binding of other cloned CRRSPs to mannose. AtCBMIP1 and SiCBMIP did not bind mannose (*SI Appendix*, Fig. S15). AFP1 showed binding to mannose at a level lower than that for OsCBMIP, but did not bind to CBM1 (*SI Appendix*, Fig. S16). Cloned CRRSPs and DUF26 domains of CRKs that did not bind to CBM1 (AT4G20670.1, AT4G23170.1, Os07g35290.1, Os07g35740.1, Os07g35750.1, Os07g35790.1) did not bind mannose either. These results suggest that the CRRSPs and DUF26 domains of CRKs tested may have other interactors than mannose and CBM1 or that their function may not involve binding to other compounds.

## Discussion

### CBMIP and CBM1: another front of plant–pathogen protein interactions

This study identified an apoplastic rice protein with two DUF26 domains that binds CBM1 of blast fungal xylanase MoCel10A. OsCBMIP binds at least two more blast fungal hydrolases and multiple proteins of *T. reesei*, suggesting that CBMIP targets multiple fungal hydrolases. In turn, screening of plant proteins using the CBM1-containing proteins MoCel10 and MoCel6A identified multiple CBMIPs from *S. italica* and Arabidopsis. *OsCBMIP* knockdown plants displayed reduced resistance against *M. oryzae*, indicating that OsCBMIP plays a key role in rice defense against this pathogen. These results reveal a hitherto undescribed mechanism of plant–pathogen protein interactions mediated by CBMIP and CBM1. We propose that plant apoplastic CBMIP proteins target conserved CBM1 motifs in various hydrolases of fungal pathogens to counter pathogen invasion. Roles for DUF26-containing proteins in plant defense have been reported in ginkgo (Gnk2) and maize (AFP1) (Miyakawa et al. 2014; Lay-Sun et al. 2018), where the proteins exhibit antifungal activity, presumably mediated by binding to pathogen mannose. OsCBMIP binds both CBM1 and mannose, but does not show antifungal activity against *M. oryzae* or *F. oxysporum*, suggesting that its role in defense is related to inhibition of fungal hydrolases mediated by CBM1-binding and is different from the function of Gnk2 and AFP1.

OsRMC, which is the same protein as OsCBMIP, is involved in seed germination and leaf growth under conditions of high NaCl concentration, but its biological function remains unclear (Zhang et al. 2008). We hypothesize that OsRMC/OsCBMIP may be a multi-functional protein possibly involved in responses to biotic and abiotic stresses. Future study is needed to elucidate the link between the two processes.

CBMs are widely distributed in fungi, bacteria and plants (*SI Appendix*, Table S2). CBM domain 18 (CBM18) is the most frequent among the *M. oryzae* CBM proteins. However, since plants also have CBM18 domains in their own proteins, this domain does not allow plants to discriminate “non-self” from “self.” By contrast, CBM1 is found only in fungal enzymes. We therefore hypothesize that targeting of CBM1 by OsCBMIP evolved to counteract fungal CWDEs.

### Function of DUF26-containing proteins and the origin of CBMIPs

There are several types of DUF26-containing proteins. The archetypical protein Gnk2 has only a single DUF26 domain, whereas numerous proteins contain two distantly related DUF26 domains, DUF26-A and DUF26-B. CRKs contain an extracellular domain with two DUF26 motifs, a transmembrane domain and an intracellular kinase domain. PDLPs contain an extracellular domain with two DUF26 motifs and a transmembrane domain. In the majority of cases, binding targets of DUF26 domains are not known. However, Gnk2 and AFP1 are reported to bind mannose. We identified CRRSPs that do not bind either mannose or CBM1 (*SI Appendix*, Fig. S12, Fig. S13, Fig. S15, Table S3). We hypothesize that the DUF26 domain was originally used for binding to mannose and other unknown compounds and, during the course of evolution, CBM1-binding capability was acquired independently in multiple DUF26 lineages through amino acid changes in the DUF26-B domain. Disulfide bridges formed by conserved cysteine residues of the CRR motif of DUF26 are predicted to contribute to structural stabilization (Vaattovaara et al. 2019), and this common fold may have provided CRRSPs with a versatile platform for binding various molecules.

Our phylogenetic analysis indicates that OsCBMIP is derived from OsCBMIP-K through truncation of the transmembrane domain and kinase domain. Since OsCBMIP-K binds CBM1, there is a possibility of signaling mediated by recognition of CBM1. Future studies should clarify whether CBM1 recognition by OsCBMIP-K plays a role in defense signaling.

In summary, our study and others suggest that DUF26 is a versatile multi-functional domain deployed by plants to bind various compounds (e.g. mannose, CBM1) and/or sense environmental conditions (e.g. ROS). Accordingly, the genes for DUF26-containing proteins were highly amplified in plant genomes. As shown in this study, a subset of DUF26-domain containing proteins evolved as apoplastic proteins to counteract pathogens by binding and inactivating CBM1-containing proteins. Future structure analysis would allow engineering of CBMIP for a higher CBM1-binding capability to generate pathogen resistant crops.

## Materials and Methods

### Materials

Water-insoluble wheat cell wall preparation was prepared from wheat coleoptiles by treating sequentially with methanol and amylase.

### Growth conditions of microorganisms and plants

*M. oryzae* was grown on oatmeal plates or YG (0.5% yeast extract, 2% glucose, w/v) medium at 25 °C. Conidium formation of *M. oryzae* was accomplished at 28 °C under dark-blue light for 4 days. *Agrobacterium tumefaciens* (GV3101) carrying plasmid vector was grown in YEB (0.5% yeast extract, 1% peptone, 0.5% beef extract, 0.5% sucrose, w/v) medium supplemented with rifampicin and kanamycin. Rice and *N. benthamiana* were grown in soil at 30 and 25 °C, respectively. Rice suspension cells were cultured at 25 °C with rotation at 140 rpm and transferred to fresh medium every 10 days as described previously (Okuyama et al. 2011).

### DNA sequencing

DNA was amplified from cDNA by PCR using PrimeStar GXL DNA polymerase (Takara Bio, Shiga, Japan) and verified by DNA sequencing using a 3130 Genetic Analyzer (Applied Biosystems, CA, USA). DNA primers used in this study are shown in *SI Appendix*, Table S4.

### Recombinant protein preparation

His-tagged or Flag-tagged *M. oryzae* proteins—xylanase (MoCel10A), CBM1-truncated xylanase (MoCel10AΔCBM), cellobiohydrolase (MoCel6A) and cellobiose dehydrogenase (MoCDH)—were overexpressed in *M. oryzae* as described previously (Takeda et al. 2010). His-tagged and HA-tagged OsCBMIPs were overexpressed in rice suspension cells as described previously (Okuyama et al. 2011). Flag-tagged CRRSPs and CRKs were expressed in *N. benthamiana* leaves as described previously (Maqbool et al. 2015). His-tagged Gnk2 was expressed in *Escherichia coli* (Origami) as described previously (Miyakawa et al. 2014). Purification of His-tagged proteins was carried out using His-tag affinity resin (His-resin) (Clontech, CA, USA).

### Pull-down assay with His-tagged proteins

Purified His-tagged MoCel10A or OsCBMIP protein was incubated with a crude protein preparation of rice leaves or *T. reesei* culture filtrate in binding buffer (sodium phosphate buffer [50 mM, pH 7.5] containing 150 mM NaCl) for 1 h at 4 °C. The mixture was further incubated with His-resin for 1 h at 4 °C. Proteins bound to His-resin were eluted using binding buffer containing 200 mM imidazole. Bound proteins were subjected to SDS-PAGE followed by silver staining. For CoIP assay, His-tagged protein and Flag-tagged protein were incubated for 1 h at 4 °C. The mixture was further incubated with His-resin for 1 h at 4 °C. The supernatant of this mixture, obtained by centrifugation at XX *g*, was used as an unbound fraction. Proteins bound to His-resin were eluted using binding buffer containing 200 mM imidazole. Both fractions were subjected to SDS-PAGE followed by immunoblotting using anti-His and anti-Flag antibodies.

### Peptide sequence identification

Protein bands separated by SDS-PAGE were excised and digested with trypsin as described previously (Kawamura and Uemura 2003). Peptide identification was carried out as described previously (Takeda et al. 2015). Digested peptides were applied onto a Magic C18 AQ nano column (0.1×150 mm, MICHROM Bioresources, Inc., CA, USA) in an ADVANCE UHPLC system (MICHROM Bioresources, Inc.) equilibrated with 0.1% formic acid (v/v) in acetonitrile and eluted using a linear gradient of 5–45% (v/v) acetonitrile at a flow rate of 500 nL/min. Mass analysis was performed using an LTQ Orbitrap XL mass spectrometer (Thermo Fisher Scientific, MA, USA) operating Xcalibur software ver. 2.0.7 (Thermo Fisher Scientific). Peptides were identified using a MASCOT MS/MS ion search (http://www.matrixscience.com/home.html) in error tolerance mode (one amino acid substitution allowed) using the NCBI database. Search parameters were as follows: taxonomy, plants; max missed cleavages, 0; fixed modifications, carbamidomethyl; peptide tolerance, ± 5 ppm; fragment mass tolerance, ± 0.6 Da.

### Gel-permeation chromatography

MoCel10A-His, OsCBMIP-His, and the mixture of MoCel10A-His and OsCBMIP-His preincubated at 4 °C for 30 min were applied on Superdex G-75 column (GE Healthcare, Buckinghamshire, UK) equilibrated with sodium phosphate buffer (50 mM, pH 7.5) and 150 mM NaCl. Proteins were detected by immunoblot analysis using anti-His antibody. Fractions contain 3 mL of the eluate. Proteins, albumin (75 kDa), carbonic anhydrase (29 kDa) and aprotinin (6.5 kDa), were used as a standard marker.

### Determination of binding kinetics

Integrated intensity representing the binding kinetics of OsCBMIP to MoCel10A, MoCel6A and MoCel10AΔCBM was determined on a BLItz instrument (ForteBio, CA, USA) using BLItz Pro software (ForteBio, CA, USA) as described previously (Song et al. 2016). OsCBMIP-His (10 µL, 2.0 µM) was loaded onto a His-tag biosensor for 10 min. A baseline of binding response was determined by incubating the sensor with binding buffer (50 mM sodium phosphate buffer, pH7.5, 150 mM NaCl) for 10 min. Association of MoCel10A, MoCel6A and MoCel10AΔCBM proteins (10 µL, 2.0 µM), respectively, was measured for a duration of 5 min. Binding buffer was used to measure protein dissociation for a duration of 4 min. Time zero was defined as the starting point of protein association. Fold increase in integrated intensity was calculated by dividing each trajectory by the value at time zero.

### Binding of MoCel10A to cellulose

MoCel10A-His (2.0 µg) was preincubated in sodium phosphate buffer (100 mM, pH 6.0) with OsCBMIP-His (0–2.0 µg) for 30 min at 4 °C. Cellulose (1 mg) was added, and the mixture was further incubated for 1 h at 4 °C with vigorous agitation. The supernatant obtained by centrifugation at 13,000 *g* and proteins eluted from cellulose by boiling in SDS-PAGE sample buffer were used as fractions unbound and bound to cellulose, respectively.

### Effects of OsCBMIP on xylan hydrolytic activity of MoCel10A

A mixture (20 µL) containing MoCel10A-His (2.0 µg), sodium phosphate buffer (100 mM, pH 6.0) and OsCBMIP-His (0-2.0 µg) was incubated for 15 min at 4 °C, and further incubated with wheat coleoptile cell wall preparation for 30 min at 30 °C. BSA was added to the mixture to adjust to a final protein content of 2.0 µg instead of OsCBMIP. Hydrolytic activity was determined by measuring absorbance at 640 nm after staining solubilized sugars from the cell wall preparation with 0.5% (w/v) anthrone in H_2_SO_4_. For hydrolytic activity towards water-soluble xylan, a reaction mixture (100 µL) containing MoCel10A-His (0.2 µg) preincubated with OsCBMIP (0-1.0 µg), water-soluble oat spelts xylan (1%, w/v) and sodium phosphate buffer (100 mM, pH 6.0) was incubated at 30 °C. BSA was added to the mixture to adjust to a final protein content of 1.0 µg instead of OsCBMIP. The activity was determined by measuring the absorbance at 410 nm after treatment with *p*-hydroxybenzoic hydrazide-HCl. Data are the means ± SD of independent three determinations.

### Extension assay of heat-inactivated wheat coleoptiles

Heat-inactivated wheat coleoptile segments were treated with MoCel10A-His (2.0 µg) or a mixture of MoCel10A-His and OsCBMIP-His (each 2.0 µg) in 200 µL of sodium phosphate buffer (100 mM, pH 6.0). Specimens treated with buffer were used as a control. Specimens were fixed between two clamps approximately 5 mm apart and loaded at a constant 200 mN for 3 min in an extensometer (TMA/SS6000, Seiko Instruments, Japan). Extension was automatically recorded by computer from time 0.5 s every 0.1 s for 1 min and every 1 s for the next 2 min. Strain was calculated as (*L*_*t*_ – *L*_0.1_)/*L*_0.1_ (*L*_*t*_, length of coleoptiles at each time point; *L*_0.1_, length of segments at time 0.1). Data are means ± SD of five independent determinations.

### Generation of transgenic rice plants

*OsCBMIP* DNA fragments, 311 bp from 456–766 of the ORF region and 433 bp from 784–1216 of the 3′-non-coding region, were cloned into pANDA vector for RNA interference of the *OsCBMIP* gene as described previously (Miki and Shimamoto 2004). Gene introduction into rice (‘Moukoto’) was carried out as described above. T_1_ *OsCBMIP*-knockdown and wild-type control progenies segregated from T_0_ heterozygous lines were used for *M. oryzae* infection assay.

### Quantitative PCR (qPCR) and reverse-transcription PCR (qRT-PCR)

Rice leaves (‘Hitomebore’ and ‘Moukoto’) spray-inoculated with *M. oryzae* spores (3.0 × 10^5^; compatible strain Ken53-33 and incompatible strain TH69-8) in 0.01% Tween-20 (v/v) were used to determine levels of *OsCBMIP* gene expression as well as for quantification of the rice *ACTIN* gene (*Os03g61970*.*1*) and *M. oryzae actin* gene (*XM_003719823*.*1*) from genomic DNA. qPCR was carried out using a Quantitect SYBR Green PCR Kit (Qiagen, Hilden, Germany) and specific DNA primers (*SI Appendix*, Table S4) in a StepOnePlus Real-Time PCR system (Applied Biosystems, CA, USA) using SYBR GreenER qPCR Super Mix (Invitrogen, CA, USA). Data are means ± SD of three independent determinations.

### Antifungal activity assay

Antifungal activity against *M. oryzae* was evaluated by measuring luciferase activity in transformed *M. oryzae* constitutively overexpressing luciferase under control of the MPG1 promoter (Takeda et al. 2010). *M. oryzae* grown on oatmeal plates was transferred to YG medium and cultured with protein additive for 30 h at 25 °C. Cells were then sonicated, and the supernatant obtained after centrifugation at 22,000 *g* was used for analyzing luciferase activity in a Varioskan LUX (Themo Scientific). Antifungal activity against *F. oxysporum* was carried out as described previously (Miyakawa et al. 2014).

### Protein detection

Protein detection was performed by silver staining and immunoblot analysis using antibodies against peptide epitope-tags and anti-OsCBMIP after SDS-PAGE. Protein concentration was determined using a Bradford protein assay kit (Thermo-Fisher, MA, USA) with bovine serum albumin (Sigma-Aldrich, MO, USA) as the standard. Proteins on a polyvinylidene fluoride membrane were stained using CBB Stain One (Nakalai, Japan).

### Phylogenetic tree reconstruction

Amino acid sequences of CRRSPs and CRKs from *O. sativa, A. thaliana, M. polymorpha* and *Selaginella moellendorffii* were retrieved from the Rice Genome Annotation Project (http://rice.uga.edu/index.shtml), The Arabidopsis Information Resource (https://www.arabidopsis.org/), EnsemblPlants Marchantia polymorpha (https://plants.ensembl.org/Marchantia_polymorpha/Info/Index) and EnsemblPlants Selaginella moellendorffii (https://plants.ensembl.org/Selaginella_moellendorffii/Info/Index), respectively. Amino acid sequences of SiCBMIP (*S. italica*), AFP1 (*Z. mays*) and Gnk2 (*G. biloba*) were added to these sequences. DUF26-containing proteins were annotated using InterproScan with default options (Zdobnov et al. 2001). The amino acid sequences of annotated DUF26 domains were aligned using MAFFT (Katoh et al. 2013) with the following method parameter set: --maxiterate 1000 –globalpair (*SI Appendix*, Table S5). A maximum-likelihood tree was reconstructed using IQ-TREE (Nguyen et al. 2015) with 1,000 bootstrap replicates calculated using UFBoot2 (Hoang et al. 2018). ModelFinder (Kalyaanamoorthy et al. 2017) was used for model selection; “WAG + R5” for Fig. S12 and “WAG + I + G4” for Fig. 3B were chosen as the best-fit models, according to BIC. Finally, the reconstructed tree was drawn using Iroki (Moore et al. 2020).

## Acknowledgements

This study was supported by JSPS KAKENHI 18K06121 to TT and by JSPS KAKENHI 15H05779 and 20H00421 to RT.

## Notes

### Competing Interest Statement

The authors have declared no competing interest.

## References

Acharya BR, Raina S, Maqbool SB, Jagadeeswaran G, Mosher SL, Appel HM, Schultz JC, Klessig DF, Raina R (2007) Overexpression of CRK13, an Arabidopsis cysteine-rich receptor-like kinase, results in enhanced resistance to Pseudomonas syringae. Plant J 50: 488–499.

Bourdais G, Burdiak P, Gauthier A, Nitsch L, Salojärvi J, Rayapuram C, Idänheimo N, Hunter K, Kimura S, Merilo E, Vaattovaara A, Oracz K, Kaufholdt D, Pallon A, Anggoro DT, Dawid G (2015) Large-scale phenomics identifies primary and fine-tuning roles for CRKs in responses related to oxidative stress. PLoS Genet 11: e1005373.

Brunkard JO, Zambryski PC (2017) Plasmodesmata enable multicellularity: new insights into their evolution, biogenesis, and functions in development and immunity. Curr Opin Plant Biol 35: 76–83.

Caillaud MC, Wirthmueller L, Sklenar J, Findlay K, Piquerez SJM, Jones AME, Robatzek S, Jones JDG,Faulkner C (2014) The plasmodesmal protein PDLP1 localises to haustoria-associated membranes during downy mildew infection and regulates callose deposition. PLoS Pathog 10: e1004496.

Chen Z (2001) A superfamily of proteins with novel cysteine-rich repeats. Plant Physiol 126: 473–476.

Chen K, Du L, Chen Z (2003) Sensitization of defense responses and activation of programmed cell death by a pathogen-induced receptor-like protein kinase in Arabidopsis. Plant Mol Biol 53: 61–74.

Chern M, Xu Q, Bart RS, Bai W, Ruan D, Sze-To WH, Canlas PE, Jain R, Chen X, Ronald PS (2016) A genetic screen identifies a requirement for cysteine-rich-receptor like kinases in rice NH1 (OsNPR1)-mediated immunity. PLoS Genet 12: eE1006049.

Cui W, Lee JY (2016) Arabidopsis callose synthases CalS1/8 regulate plasmodesmal permeability during stress. Nat Plants 2: 16034.

Czernic P, Visser B, Sun W, Savoure A, Deslandes L, Marco Y, Van Montagu M, Verbruggen N (1999) Characterization of an Arabidopsis thaliana receptor-like protein kinase gene activated by oxidative stress and pathogen attack. Plant J 18: 321–327.

Di C, Zhang M, Xu S, Cheng T, An L (2006) Role of poly-galacturonase inhibiting protein in plant defense. Crit Rev Microbiol 32: 91–100.

Du L, Chen Z (2000) Identification of genes encoding novel receptor-like protein kinases as possible target genes of pathogen-induced WRKY DNA-binding proteins. Plant J 24: 837–848.

Du D, Liu M, Xing Y, Chen X, Zhang Y, Zhu M, Lu X, Zhang Q, Ling Y, Sang X, Li Y, Zhang C, He G (2019) Semi-dominant mutation in the cysteine-rich receptor-like kinase gene, ALS1, conducts constitutive defense response in rice. Plant Biol 21: 25–34.

Gomez-Gomez L, Boller T (2002) Flagellin perception: a paradigm for innate immunity. Trends Plant Sci 7: 251–256.

Hägglund P, Eriksson T, CollEén A, Nerinckx W, Claeyssens M, Stålbrand H (2003) A cellulose-binding module of the Trichoderma reesei β-mannanase Man5A increases the mannan-hydrolysis of complex substrates. J Biotechnol 101: 37–48.

Hardie DG (1999) Plant protein serine/threonine kinases: classification and functions. Annu Rev Plant Physiol Mol Biol 50: 97–131.

Hefford MA, Laderoute K, Willick GE, Yaguchi M, Seligy VL (1992) Bipartite organization of the Bacillus subtilis endo-beta-1,4-glucanase revealed by C-terminal mutations. Protein Eng 5: 433–439.

Hervéa C, Rogowskib A, Blakea AW, Marcusa SE, Gilbertb HJ, Knoxa JP (2010) Carbohydrate-binding modules promote the enzymatic deconstruction of intact plant cell walls by targeting and proximity effects. Proc Natl Acad Sci USA 107: 15293–15298.

Hoang DT, Chernomor O, von Haeseler A, Minh BQ, Vinh LS (2018) UFBoot2: Improving the Ultrafast Bootstrap Approximation. Mol Biol Evol 35: 518–522.

Idänheimo N, Gauthier A, Salojärvi J, Siligato R, Brosché M, Kollist H, Mähönen AP, Kangasjärvi J, Wrzaczek M (2014) The Arabidopsis thaliana cysteine-rich receptor-like kinases CRK6 and CRK7 protect against apoplastic oxidative stress. Biochem Biophys Res Commun 445: 457–462.

Ito J, Fujita Y, Ueda M, Fukuda H, Kondo A (2004) Improvement of cellulose-degrading ability of a yeast strain displaying Trichoderma reesei endoglucanase II by recombination of cellulose-binding domains. Biotechnol Prog 20: 688–691.

Jashni MK, Mehrabi R, Collemare J, Mesarich CH, deWit PJGM (2015) The battle in the apoplast: further insights into the roles of proteases and their inhibitors in plant-pathogen interactions. Front Plant Sci 6: 584.

Jiang J, Li J, Xu Y, Han Y, Bai Y, Zhou G, Lou Y, Xu Y, Chong K (2007) RNAi knockdown of Oryza sativa root meander curling gene led to altered root development and coiling which were mediated by jasmonic acid signaling in rice. Plant Cell Environ 30: 690–699.

Jones JD, Dangl JL (2006) The plant immune system. Nature 444: 323–329.

Juge N (2006) Plant protein inhibitors of cell wall degrading enzymes. Trends Plant Sci 11: 359–367.

Kalyaanamoorthy S, Minh BQ, Wong TKF, von Haeseler A, Jermiin LS (2017) ModelFinder: fast model selection for accurate phylogenetic estimates. Nat Methods 14: 587–589.

Katoh K, Standley DM (2013) MAFFT Multiple Sequence Alignment Software Version 7: Improvements in Performance and Usability. Mol Biol Evol 30: 772–780.

Kawamura Y, Uemura M (2003) Mass spectrometric approach for identifying putative plasma membrane proteins of Arabidopsis leaves associated with cold acclimation. Plant J 36: 141–154.

Kim JY, Park SC, Hwang I, Cheong H, Nah JW, Hahm KS, Park Y (2009) Protease inhibitors from plants with antimicrobial activity. Int J Mol Sci 10: 2860–2872.

Kim SG, Wang Y, Lee KH, Park ZY, Park J, Wu J, Kwon SJ, Lee YH, Agrawal GK, Rakwal R, Kim ST, Kang KY (2013) In-depth insight into in vivo apoplastic secretome of rice-Magnaporthe oryzae interaction. J Proteomics 78: 58–71.

Kraulis J, Clore GM, Milges M, Jones TA, Pettersson G, Knowles J, Gronenborn AM (1989) Determination of the three-dimensional solution structure of the C-terminal domain of cellobiohydrolase I from Trichoderma reesei. A study using nuclear magnetic resonance and hybrid distance geometry-dynamical simulated annealing. Biochemistry 28: 7241–7257.

Kunze G, Zipfel C, Robatzek S, Niehaus K, Boller T, Felix G (2004) The N terminus of bacterial elongation factor Tu elicits innate immunity in Arabidopsis plants. Plant Cell 16: 3496–3507.

Lee DS, Kim YC, Kwon SJ, Ryu CM, Park OK (2017) The Arabidopsis cysteine-rich receptor-like kinase CRK36 regulates immunity through interaction with the cytoplasmic kinase BIK1. Front Plant Sci 8: 1856.

Lehtiö J, Sugiyama J, Gustavsson M, Fransson L, Linder K, Teeri TT (2003) The binding specificity and affinity determinants of Family 1 and Family 3 cellulose binding modules. Proc Natl Acad Sci USA 100: 484–489.

Lim GH, Shine MB, de Lorenzo L, Yu K, Cui W, Navarre D, Hunt AG, Lee JY, Kachroo A, Kachroo P (2016) Plasmodesmata localizing proteins regulate transport and signaling during systemic acquired immunity in plants. Cell Host Microbe 19: 541–549.

Ma LS, Wang L, Trippel C, Mendoza-Mendoza A, Ullmann S, Moretti M, Carsten A, Kahnt J, Reissmann S, Zechmann B, Bange G, Kahmann R (2018) The Ustilago maydis repetitive effector Rsp3 blocks the antifungal activity of mannose-binding maize proteins. Nat Commun 9: 1711–1725.

Maqbool A, Saitoh H, Franceschetti M, Stevenson CEM, Uemua A, Kanzaki H, Kamoun S, Terauchi R, Banfield MJ (2015) Structural basis of pathogen recognition by an integrated HMA domain in a plant NLR immune receptor. eLife 4:e08709.

Mattinen ML, Linder M, Teleman A, Annila A (1997) Interaction between cellohexaose and cellulose binding domains from Trichoderma reesei cellulases. FEBS Lett 407: 291–296.

Miki D and Shimamoto K (2004) Simple RNAi vector for stable and transient suppression of gene function in rice. Plant Cell Physiol 45: 490–495.

Miller MA (1972) New reaction for colorimetric determination of carbohydrates. Anal Biochem 47: 273–279.

Miyakawa T, Miyazono K, Sawano Y, Hatano K, Tanokura M (2009) Crystal structure of ginkbilobin-2 with homology to the extracellular domain of plant cysteine-rich receptor-like kinases. Proteins 77: 247–251.

Miyakawa T, Hatano K, Miyauchi Y, Suwa Y, Sawano Y, Tanokura M (2014) A secreted protein with plant-specific cysteine-rich motif functions as a mannose-binding lectin that exhibits antifungal activity. Plant Physiol 166: 766–778.

Moore RM, Harrison AO, McAllister SM, Polson SW, Wommack KE (2020) Iroki: automatic customization and visualization of phylogenetic trees. PeerJ 8: e8584.

Nguyen LT, Schmidt HA, von Haeseler A, Minh BQ (2015) IQ-TREE: A Fast and Effective Stochastic Algorithm for Estimating Maximum-Likelihood Phylogenies. Mol Biol Evol 32: 268–274.

Ohtake Y, Takahashi T, Komeda Y (2000) Salycilic acid induces the expression of a member of receptor-like kinase genes in Arabidopsis thaliana. Plant Cell Physiol 41: 1038–1044.

Okuyama Y, Kanzaki H, Abe A, Yoshida K, Tamiru M, Saitoh H, Fujibe T, Matsumura H, Shenton M, Galam CD, Undan J, Ito A, Sone T, Terauchi R (2011) A multifaceted genomics approach allows the isolation of the rice Pia-blast resistance gene consisting of two adjacent NBS-LRR protein genes. Plant J 66: 467–479.

Penttilä M, Nevalainen H, Rättö M, Salminen E, Knowles J (1987) A versatile transformation system for the cellulolytic filamentous fungus Trichoderma reesei. Gene 61: 155–164.

Sawano Y, Miyakawa T, Yamazaki H, Tanokura M, Hatano K (2007) Purification, characterization, and molecular gene cloning of an antifungal protein from Ginkgo biloba seeds. Biol Chem 388: 273–280.

Shibata N, Suetsugu M, Kakeshita H, Igarashi K, Hagihara H, Takimura Y (2017) A novel GH10 xylanase from Penicillium sp. accelerates saccharification of alkaline-pretreated bagasse by an enzyme from recombinant Trichoderma reesei expressing Aspergillus β-glucosidase. Biotechnol Biofuels 10: 278–294.

Song D, Graham TGW, Loparo JJ (2016) A general approach to visualize protein binding and DNA conformation without protein labelling. Nat Commun 7: 10976–10982.

Spoel SH, Dong X (2012) How do plants acieve immunity? Defence without specialized immune cells. Nat Rev Immunol 12: 89–100.

Stergiopoulos, I. and Pierre, J.G.M. de Wit. (2009) Fungal effector proteins. Annu Rev Phytopathol 47: 233–263.

Sweigard JA, Chumley F, Carroll A, Farrall L, Calent B (1997) A series of vectors for fungal transformation. Fungal Genet Newsl 44: 52–55.

Takahashi M, Takahashi H, Nakano Y, Konishi T, Terauchi R, Takeda T (2010) Characterization of a cellobiohydrolase (MoCel6A) produced by Magnaporth oryzae. Appl Environ Microbiol 76: 6583–6590.

Takahashi M, Yamamoto R, Sakurai N, Nakano Y, Takeda T (2014) Fungal hemicelluloses-degrading enzymes cause physical property changes concomitant with solubilization of cell wall polysaccharides. Planta 241: 359–370.

Takeda T, Nakano Y, Takahashi M, Konno N, Sakamoto Y, Arashida R, Marukawa Y, Yoshida E, Ishikawa T, Suzuki K (2015) Identification and enzymatic characterization of an endo-1,3-β-glucanase from Euglena gracilis. Phytochemistry 116, 21–27.

Takeda T, Furuta Y, Awano T, Mizuno K, Mitsuishi Y, Hayashi T (2002) Suppression and acceleration of cell elongation by integration of xyloglucans in pea stem segments. Proc Natl Acad Sci USA 99: 9055–9060.

Takeda T, Takahashi M, Nakanishi-Masuno T, Nakano Y, Saitoh H, Hirabuchi A, Fujisawa S, Terauchi R (2010) Characterization of endo-1,3-1,4-β-glucanase in GH family 12 from Magnaporthe oryzae. Appl Microbiol Biotechnol 88: 1113–1123.

Tanaka H, Osakabe Y, Katsura S, Mizuno S, Maruyama K, Kusabake K, Mizoi J, Shinozaki K, Yamaguchi Shinozaki K (2012) Abiotic stress-inducible receptor-like kinases negatively control ABA signaling in Arabidopsis. Plant J 70 599–613.

Vaattovaara A, Brandt B, Rajaraman S, Safronov O, Veidenberg A, Luklová M, Kangasjärvi J, Löytynoja A, Hothorn M, Salojärvi J, Wrzaczek M (2017) Mechanistic insights into the evolution of DUF26-containing proteins in land plants. Commun Biol 10.1038/s42003-019-0306-9.

Valueva TA, Mosolov VV (2014) Role of inhibitors of proteolytic enzymes in plant defense against phytopathogenic microorganisms. Biochemistry 69: 1305–1309.

van der Hoorn RAJ (2008) Plant proteases: From phenotypes to molecular mechanisms. Annu Rev Plant Biol 59: 191–223.

Vorwerk S, Somerville S, Somerville C (2004) The role of plant cell wall polysaccharide composition in disease resistance. Trends Plant Sci 9: 203–209.

Walton, J.D. (1994) Deconstructing the cell wall. Plant Physiol 104: 1113–1118.

Yadeta K, Elmore JM, Creer AY, Feng B, Franco JY, Rufian JS, He P, Phinney B, Coaker G (2017) A cysteine-rich protein kinase associates with a membrane immune complex and the cysteine residues are required for cell death. Plant Physiol 173: 771–787.

Yeh YH, Chang YH, Huang PY, Huang JB, Zimmerli L (2015) Enhanced Arabidopsis pattern-triggered immunity by overexpression of cysteine-rich receptor-like kinases. Front Plant Sci 6: 322.

York WS, Qin Q, Rose JK (2004) Proteinaceous inhibitors of endo-β-glucanases. Biochim Biophys Acta 1696: 223–233.

Zhang L, Tian LH, Zhao JF, Song Y, Zhang CJ, Guo Y (2009) Identification of an apoplastic protein involved in the initial phase of salt stress response in rice root two-dimensional electrophoresis. Plant Physiol 149: 916–928.

Zdobnov EM, Apweiler R (2001) InterProScan – an integration platform for the signature-recognition methods in InterPro. Bioinfomatics 17: 847–848.

