## Supplemental data for "Apoplastic CBM1-interacting proteins bind conserved carbohydrate binding module 1 motifs in fungal hydrolases to counter pathogen invasion"

### SUPPLEMENTARY FIGURES

#### Fig. S1.

```
MARCTLLVLL VAAAVAVVPL AAGQPWATCG DGTYEQGSAY ENNLLNLALT

LRDGASSQEI LFSTGSNGAA PNTVYGLLLC RGDISRAACY DCGTSVWRDA
    1)                                    2)

GSACRRAKDV ALVYNECYAR LSDKDDFLAD KVGPGQLTTL MSSTNISSGA
        3)               5)
          4)

DVAAYDRAVT RLLAATAEYA AGDIARKLFA TGQRVGADPG FPNLYATAQC
             6)                      8)
             7)

AFDITLEACR GCLEGLVARW WDTFPANVDG ARIAGPRCLL RSEVYPFYTG
           9)       10)                      11, 12)
                                             13, 14)

APMVVLRE
```

**Fig. S1. Peptide sequences of OsCBMIP detected by LC-MS/MS.**
A rice protein band detected by pull-down assay using MoCel10A-His was treated with trypsin and the resulting peptide fragments were analyzed by LC-MS/MS. The 14 peptides identified are underlined. Methionine residues of peptides 12 and 14 were oxidized. CRR motifs are indicated in red.

### Fig. S2.

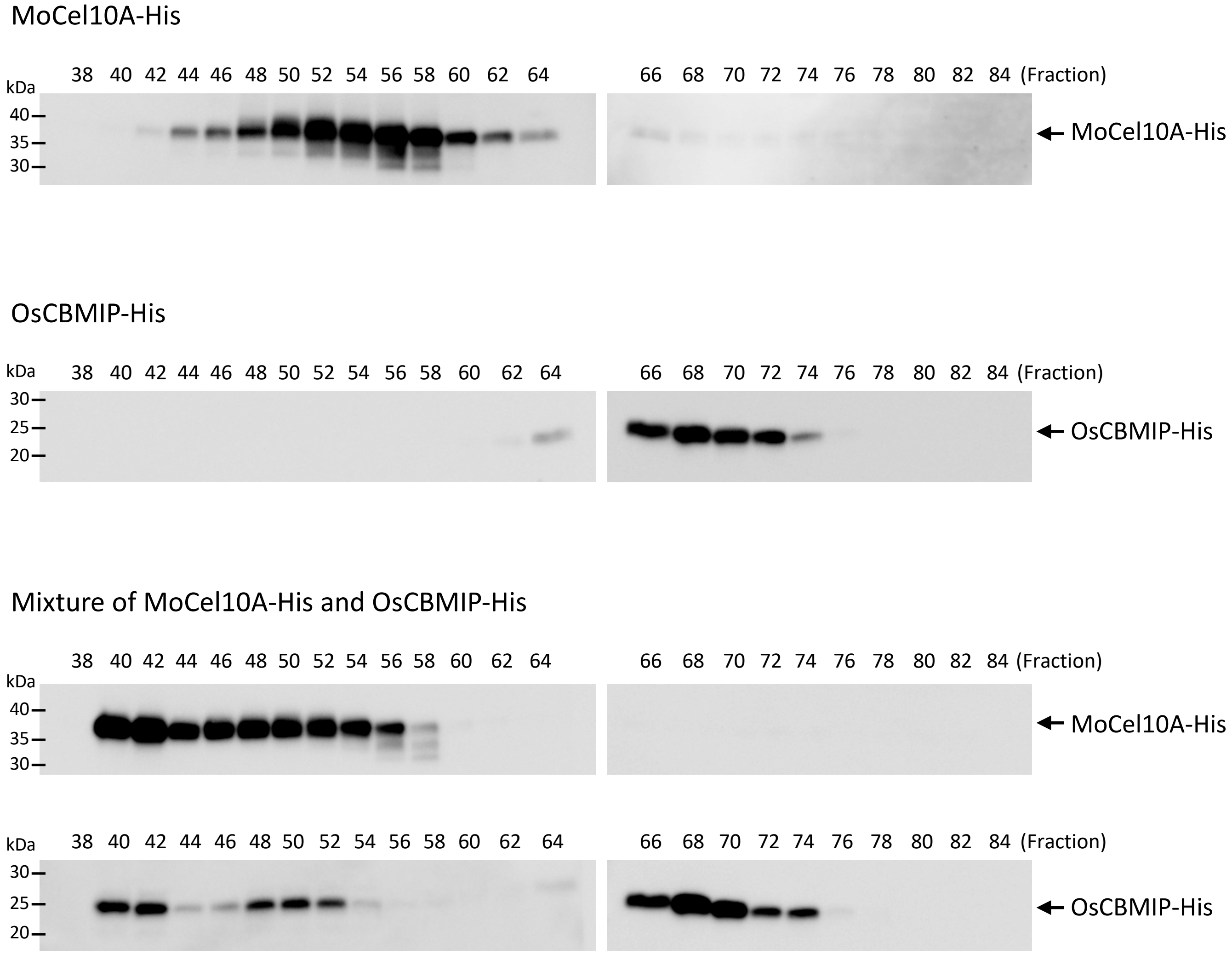


**Fig. S2. Gel-permeation chromatography of monomers and the complex of OsCBMIP and MoCel10A.**
MoCel10A-His, OsCBMIP-His, and a mixture of MoCel10A-His and OsCBMIP-His preincubated at 4 ˚C for 30 min were separated on a Superdex G-75 column equilibrated with sodium phosphate buffer (50 mM, pH 7.5) and 150 mM NaCl. Proteins were detected by immunoblot analysis using anti-His antibody. Protein standard markers albumin (75 kDa), carbonic anhydrase (29 kDa) and aprotinin (6.5 kDa) were eluted in fractions 48, 60 and 72, respectively.

### Fig. S3.

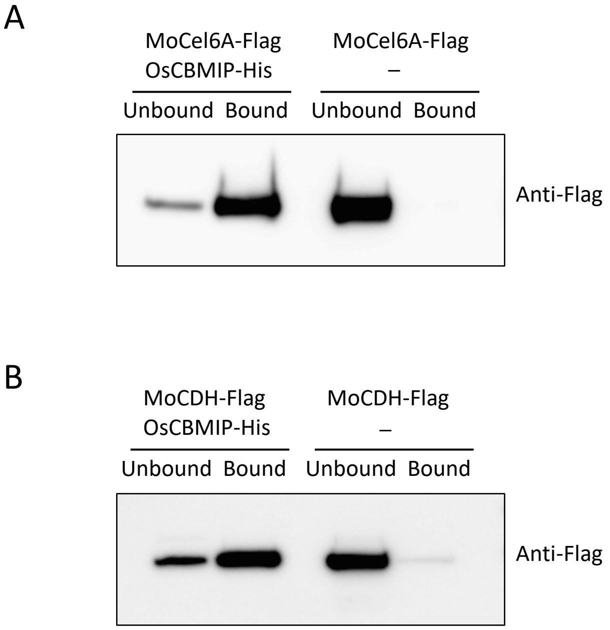


**Fig. S3. Binding of OsCBMIP to CBM1-containing proteins MoCel6A and MoCDH.** Crude protein preparations (20 μg) containing (A) Flag-tagged MoCel6A (MoCel6A-Flag) or (B) MoCDH (MoCDH-Flag) prepared from *M. oryzae* were incubated in sodium phosphate buffer (50 mM, pH 7.5) containing 150 mM NaCl with or without OsCBMIP-His (2.0 μg) for 1 h at 4 ˚C before being further incubated with His-resin. Fractions unbound and bound to His-resin were subjected to SDS-PAGE followed by immunoblot analysis using anti-Flag antibody.

Fig. S4.

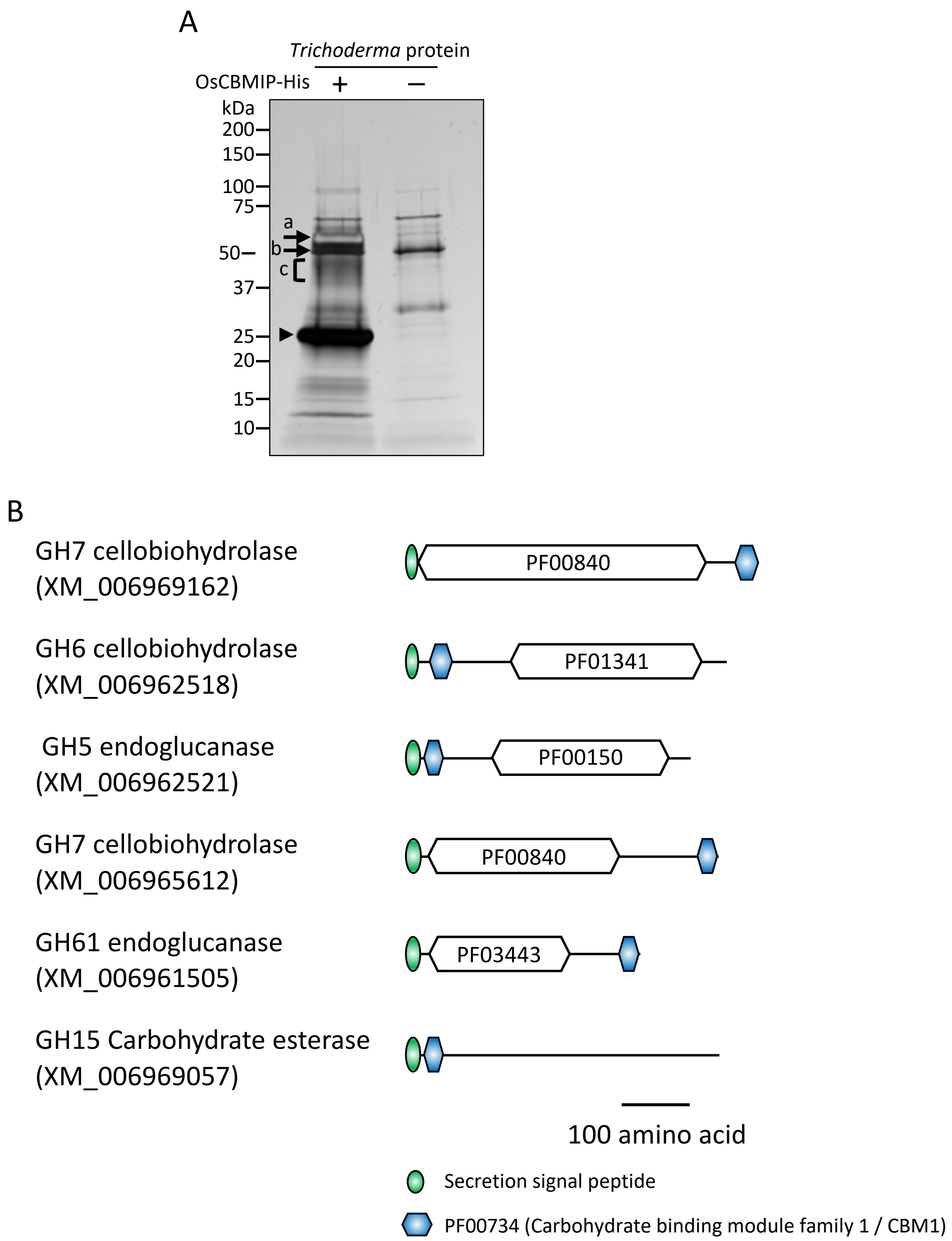


**Fig. S4. Identification of *T. reesei* proteins bound to OsCBMIP.**
(A) Culture filtrate from *T. reesei* culture grown in minimal medium (Penttilä et al. 1987) buffered to pH 6.0 and supplemented with 2% (w/v) cellobiose was incubated with (+) or without (-) OsCBMIP-His, and further incubated with His-resin. Fractions bound to His-resin were subjected to SDS-PAGE followed by silver staining. Arrowhead indicates OsCBMIP-His. Identification of proteins bound to OsCBMIP, indicated by arrows (a–c), was carried out by LC-MS/MS. Proteins labeled (a) and (b) correspond to GH7 cellobiohydrolase (XM_006969162) and GH6 cellobiohydrolase (XM_006962518), respectively. Proteins labeled (c) with negligible levels at the same position in the control (-) were GH5 endoglucanase (XM_006962521), GH7 cellobiohydrolase (XM_006965612), GH61 endoglucanase (XM_006961505) and GH15 carbohydrate esterase (XM_006969057). (B) Schematic structures of *T. reesei* proteins bound to OsCBMIP. All proteins possess CBM1 at the N- or C-terminal region connected to the catalytic core domain.

Fig. S5.

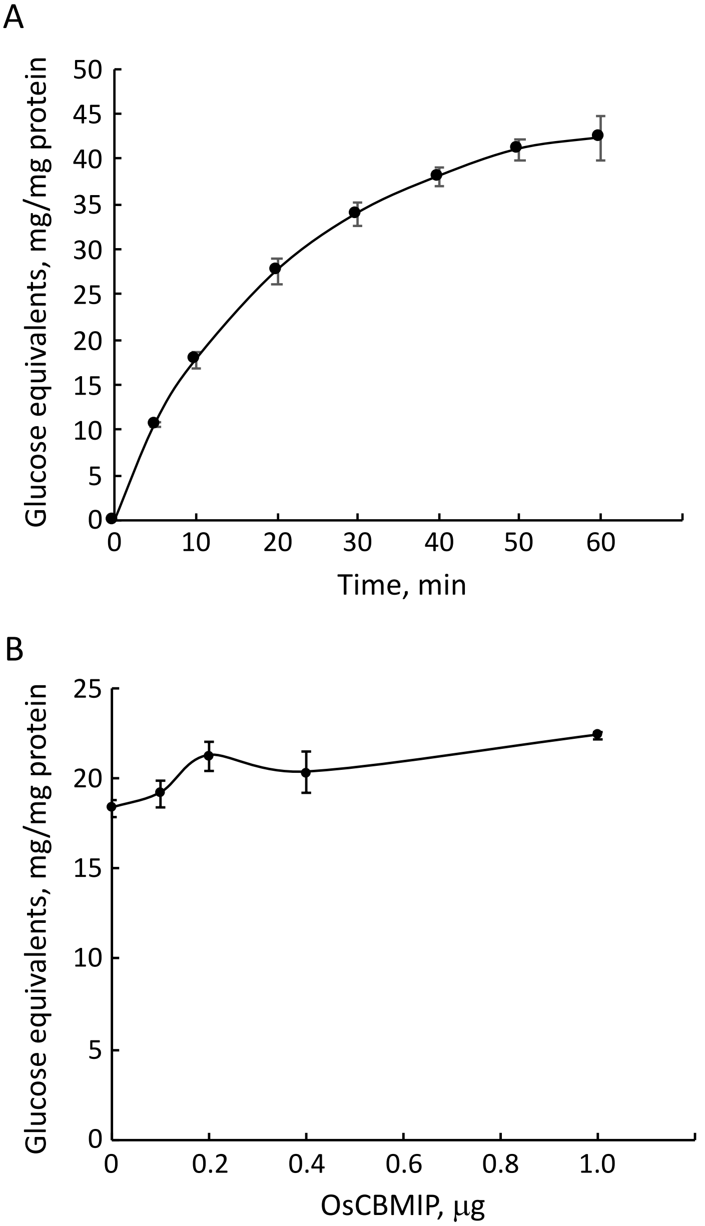


**Fig. S5. Hydrolytic activity of MoCel10A towards water-soluble xylan.**
(A) Dependence of the hydrolytic activity of MoCel10A-His towards water-soluble oat spelt xylan on incubation time. (B) Effects of OsCBMIP on the hydrolytic activity of MoCel10A-His towards water-soluble xylan. MoCel10A-His (0.2 μg) was preincubated in sodium phosphate buffer (100 mM, pH 6.0) with OsCBMIP-His (0–1.0 μg) at 4 ˚C for 30 min before further incubating with water-soluble oat spelt xylan at 30 ˚C for 10 min. Data are means ± SD of three independent determinations.

Fig. S6.

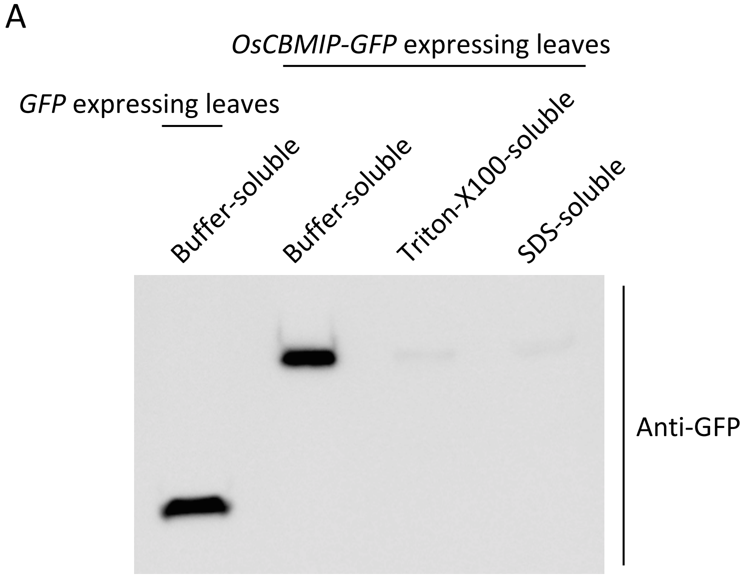


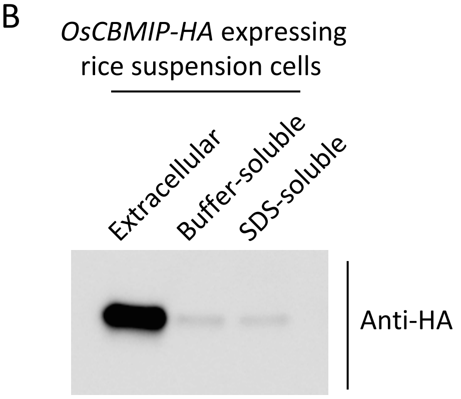


**Fig. S6. Fractionation of OsCBMIP protein expressed in *N. benthamiana* leaves and rice suspension cells.**

(A) OsCBMIP-GFP expressed in *N. benthamiana* leaves was sequentially extracted using sodium phosphate buffer (100 mM, pH 6.0) containing 100 mM NaCl (buffer-soluble), or the same buffer containing 1% (v/v) Triton-X100 (Triton-X100-soluble) or 1% (w/v) SDS (SDS-soluble). Buffer-soluble protein from *N. benthamiana* leaves overexpressing GFP was used as a GFP marker. (B) Rice suspension cells overexpressing OsCBMIP-HA were separated into culture filtrate (extracellular) and cells. Proteins were extracted from cells using sodium phosphate buffer (100 mM, pH 6.0) containing 100 mM NaCl (buffer-soluble) or 1% (w/v) SDS (SDS-soluble). The prepared proteins (1.0 mg) were subjected to SDS-PAGE followed by immunoblot analysis using anti-GFP and anti-HA antibodies.

Fig. S7.

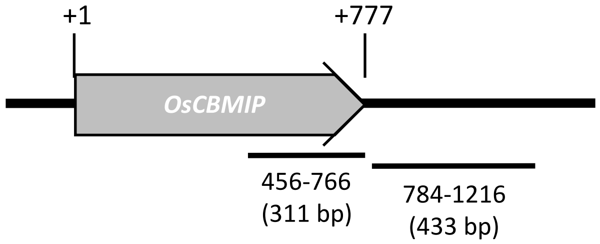


**Fig. S7. Gene structure of *OsCBMIP* and DNA fragments used for RNAi experiment.**
The *OsCBMIP* gene consists of one exon of 777 bp. DNA fragments of 311 bp in the ORF and 433 bp in 3′-UTR region were used for RNAi experiments.

### Fig. S8.

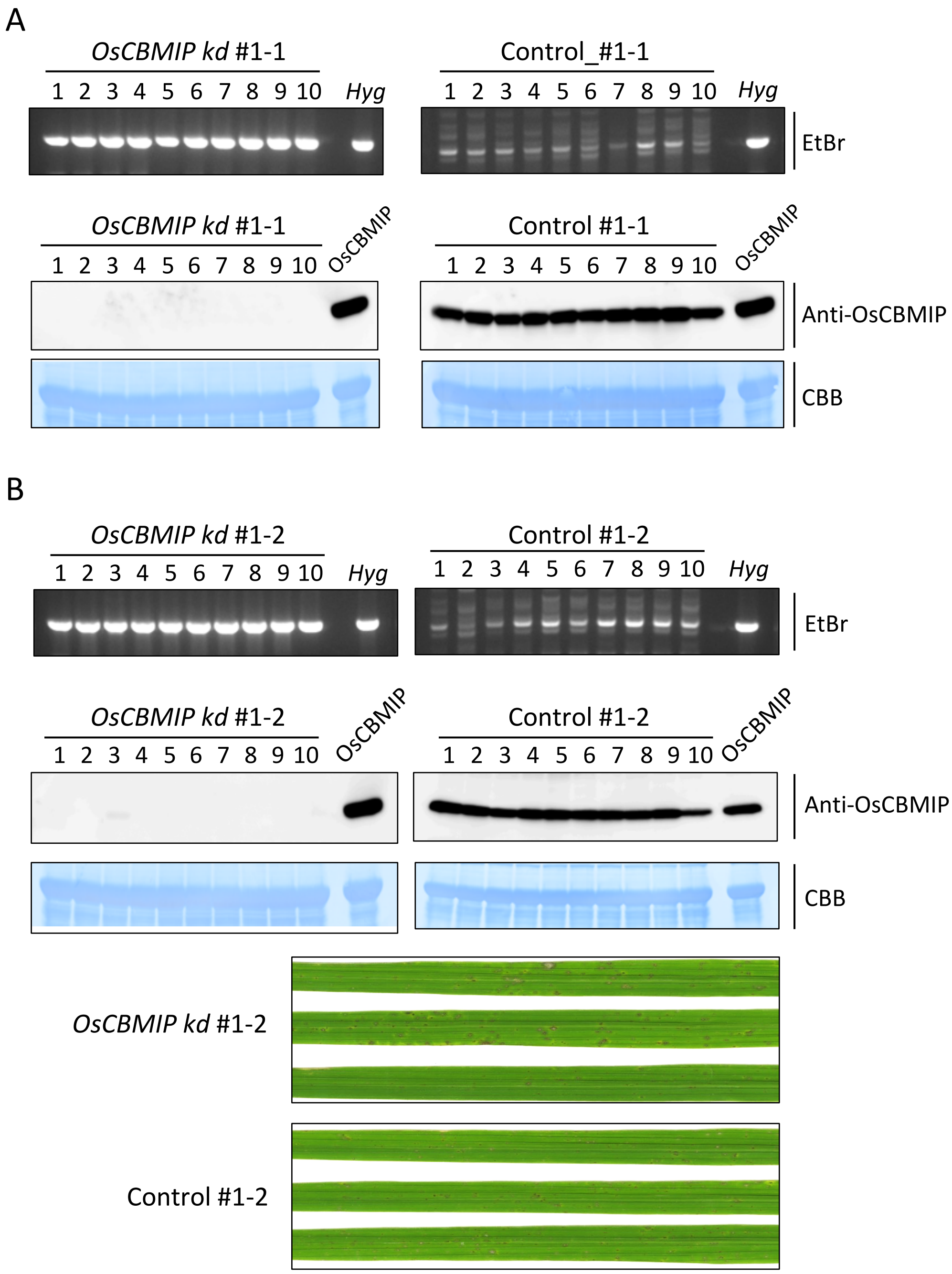
A
OsCBMIP kd #1-1
1 2 3 4 5 6 7 8 9 10
Hyg
Control_#1-1
1 2 3 4 5 6 7 8 9 10
Hyg
EtBr
OsCBMIP kd #1-1
1 2 3 4 5 6 7 8 9 10
OsCBMIP
Control #1-1
1 2 3 4 5 6 7 8 9 10
OsCBMIP
Anti-OsCBMIP
CBB
B
OsCBMIP kd #1-2
1 2 3 4 5 6 7 8 9 10
Hyg
Control #1-2
1 2 3 4 5 6 7 8 9 10
Hyg
EtBr
OsCBMIP kd #1-2
1 2 3 4 5 6 7 8 9 10
OsCBMIP
Control #1-2
1 2 3 4 5 6 7 8 9 10
OsCBMIP
Anti-OsCBMIP
CBB
OsCBMIP kd #1-2
Control #1-2

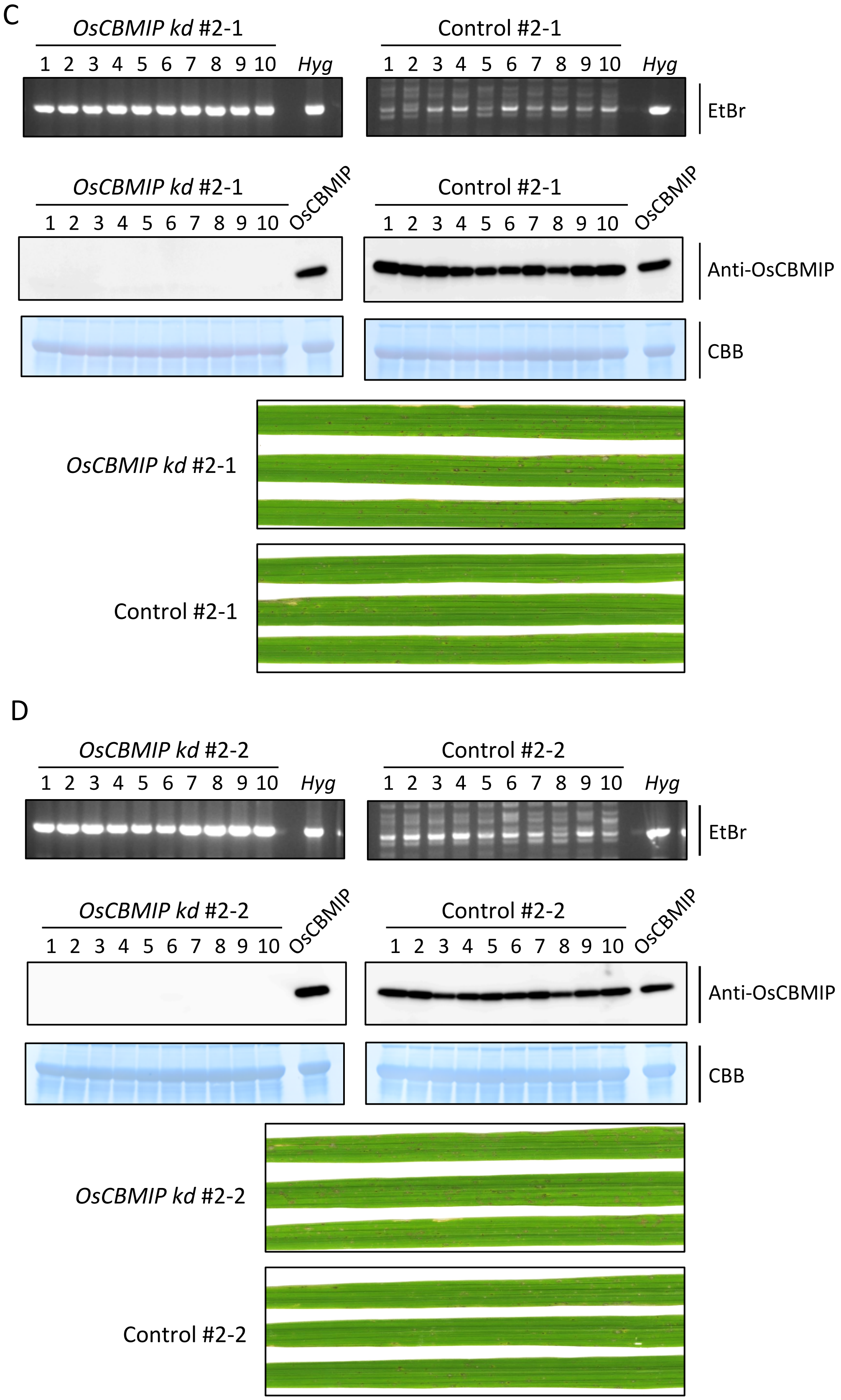


**Fig. S8. RNAi-mediated knockdown of the *OsCBMIP* gene in rice.**
PCR amplification of the hygromycin transgene, OsCBMIP protein accumulation and disease symptoms after *M. oryzae* inoculation of $T_1$ progeny segregating for *OsCBMIP*-knockdown and *OsCBMIP* expression. (A) *OsCBMIP*-knockdown (*kd*) #1-1 and *OsCBMIP*-expressing control #1-1 lines; (B) *OsCBMIP*-knockdown (*kd*) #1-2 and *OsCBMIP*-expressing control #1-2 lines; (C) *OsCBMIP*-knockdown (*kd*) #2-1 and *OsCBMIP*-expressing control #2-1 lines; (D) *OsCBMIP*-knockdown (*kd*) #2-2 and *OsCBMIP*-expressing control #2-2 lines.

Fig. S9.

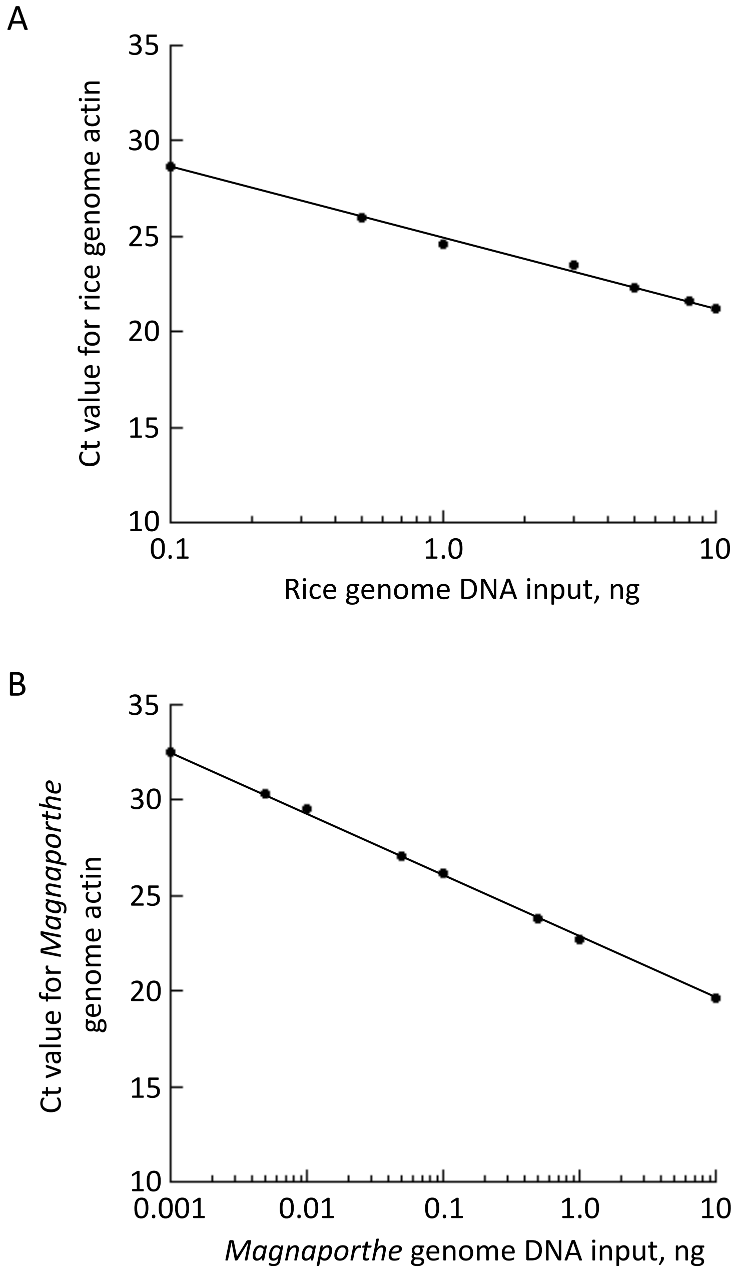


**Fig. S9. Determination of qPCR conditions for evaluating the level of *M. oryzae* infection in rice plants.**

(A) Variable amounts (0.1–10 ng per 15 μL reaction mixture) of rice genomic DNA were used to amplify the rice *ACTIN* gene by qPCR. Ct values decreased linearly as the amount of rice genomic DNA increased. (B) Variable amounts (0.001–10 ng) of *M. oryzae* genomic DNA were mixed with rice genomic DNA to a final content of 10 ng. The mixture was used for qPCR to amplify the *M. oryzae ACTIN* gene. Ct values decreased linearly as the amount of *M. oryzae* genomic DNA increased.

Fig. S10.

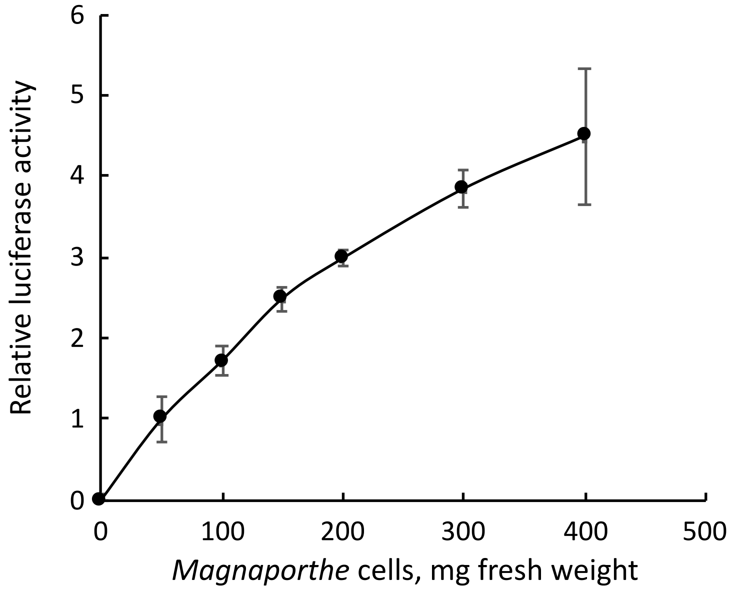


**Fig. S10. Determination of luciferase assay conditions for evaluating growth of *M. oryzae*.** Variable amounts (50–400 mg fresh weight) of luciferase-expressing *M. oryzae* were examined for luciferase activity. The average value for 50 mg fresh weight of *M. oryzae* was defined as the unit for the ratio. Activity increased linearly as the fresh weight increased up to 300 mg.

Fig. S11.

A

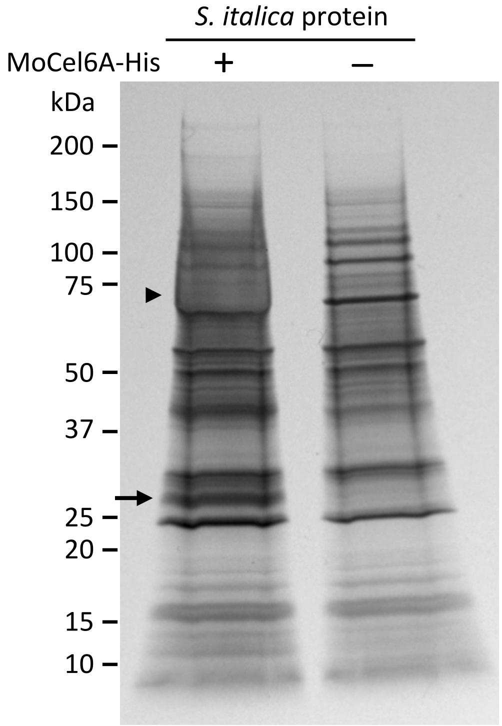


B

```
MILHASKTSR LTAGRRRHPQ TSQSMASSPV LVLLVLIAAT AAAPQLSSAV

DPVGTYCAKN FTGAQTQASI SQVLAALVPR ASAAYYATAT AGSGSSAIWG
                                 1)
LAQCRGDIPA SDCALCISAA AKQAASSCRG QADVRLWYDY CFLRYTDADF
            2)                                3)
LGLPDTGYTL ILINTMNASD PAAFDRAERK LMARVAAEAG DAASGGLVRE
                                        4)
TARFGSATTI YGLGWCTRDI TAADCGLCVA QAVAELPNYC QFRRGCRVLY
   5)                                                6, 7)
SSCMARYETY PFFFPVSGAA AASSHAGEYE KVILNHHS
```

**Fig. S11. Identification of CBM1-binding protein from *Setaria italica*.**
(A) Protein was extracted from *S. italica* leaves 4 days after *M. oryzae* inoculation using sodium phosphate buffer (50 mM, pH 7.5) containing 150 mM NaCl. Protein preparation was incubated with (+) or without (-) MoCel6A-His for 1 h at 4 ° C before being further incubated with His-resin. Fractions bound to His-resin were subjected to SDS-PAGE followed by silver staining. Arrowhead indicates added MoCel6A-His. Arrow indicates a candidate protein that interacts with MoCel6A-His. (B) The candidate protein was identified as a member of the CRRSPs by LC-MS/MS. Peptide sequences obtained are underlined. The methionine residue of peptide number 7 was oxidized. CRR motifs are indicated in red.

Fig. S12.

A

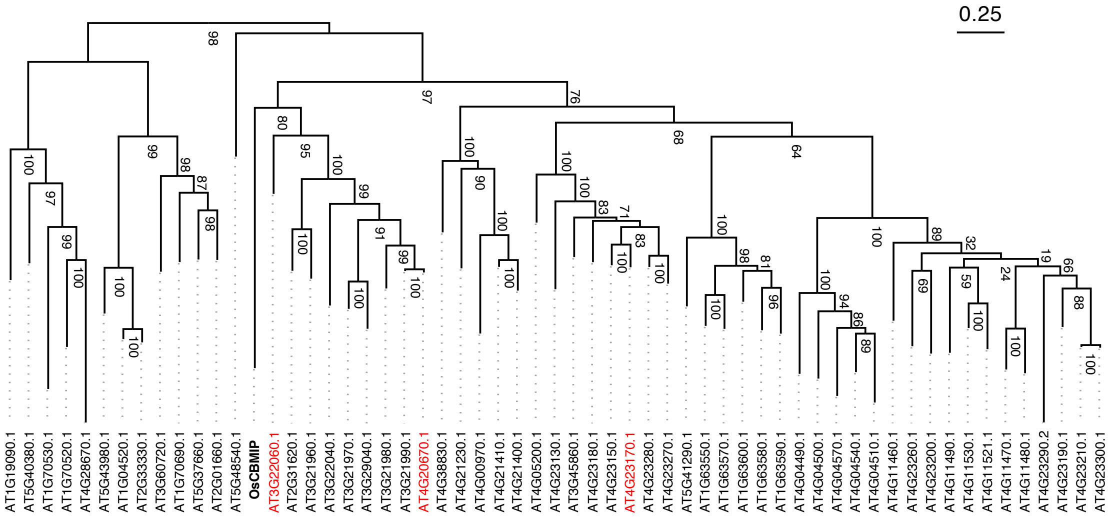


B

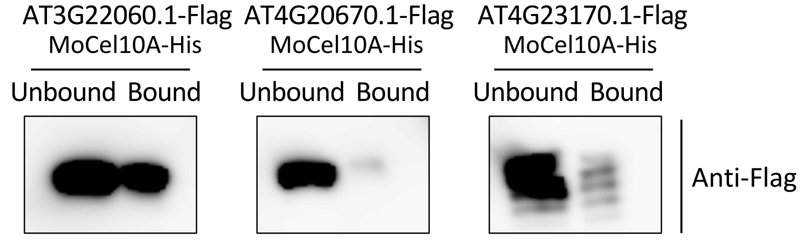


**Fig. S12. Identification of *A. thaliana* CRRSP that interacts with CBM1.**
(A) Phylogenetic tree of DUF26 domains (DUF26-A and DUF26-B) from *A. thaliana* CRRSPs and CRKs along with OsCBMIP, reconstructed using amino acid sequences. Bar, 0.25 amino acid substitutions per site . AtCRRSPs assayed for binding to MoCel10A-His are indicated in red. (B) Flag-tagged Arabidopsis CRRSPs were expressed in *N. benthamiana* leaves and extracted using sodium phosphate buffer (50 mM, pH 7.5) containing 150 mM NaCl. Prepared proteins were incubated with MoCel10A-His for 1 h at 4 ˚C before further incubating with His-resin. Fractions unbound and bound to His-resin were subjected to SDS-PAGE followed by immunoblot analysis using anti-Flag. A protein encoded by *AT3G22060.1* bound to CBM1.

Fig. S13.

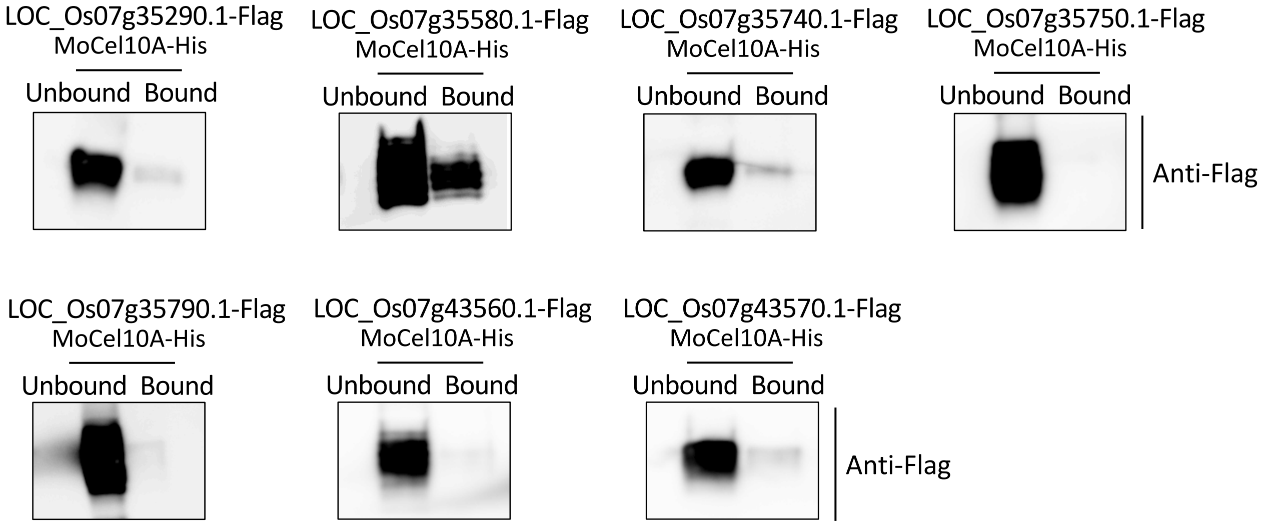


**Fig. S13. Identification of *O. sativa* CRK that interacts with CBM1.**
(A) DUF26 domains from *O. sativa* CRKs were expressed in *N. benthamiana* leaves and extracted using sodium phosphate buffer (50 mM, pH 7.5) containing 150 mM NaCl. Prepared proteins were incubated with MoCel10A-His for 1 h at 4 ˚C before being further incubated with His-resin. Fractions unbound and bound to His-resin were subjected to SDS-PAGE followed by immunoblot analysis using anti-Flag. DUF26 domains encoded by *LOC_Os07g35580.1* bound CBM1.

Fig. S14

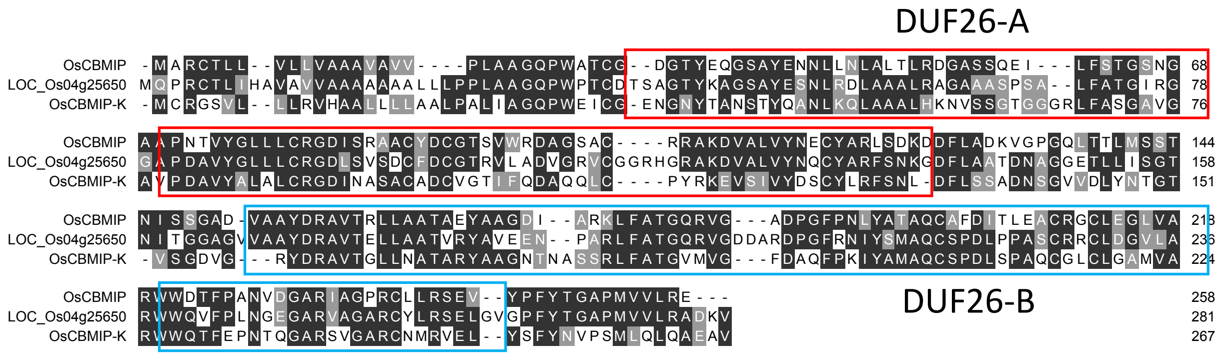


**Fig. S14. Amino acid sequence alignment among OsCBMIP, LOC_Os04g25650.1 and OsCBMIP-K.**

Fig. S15.

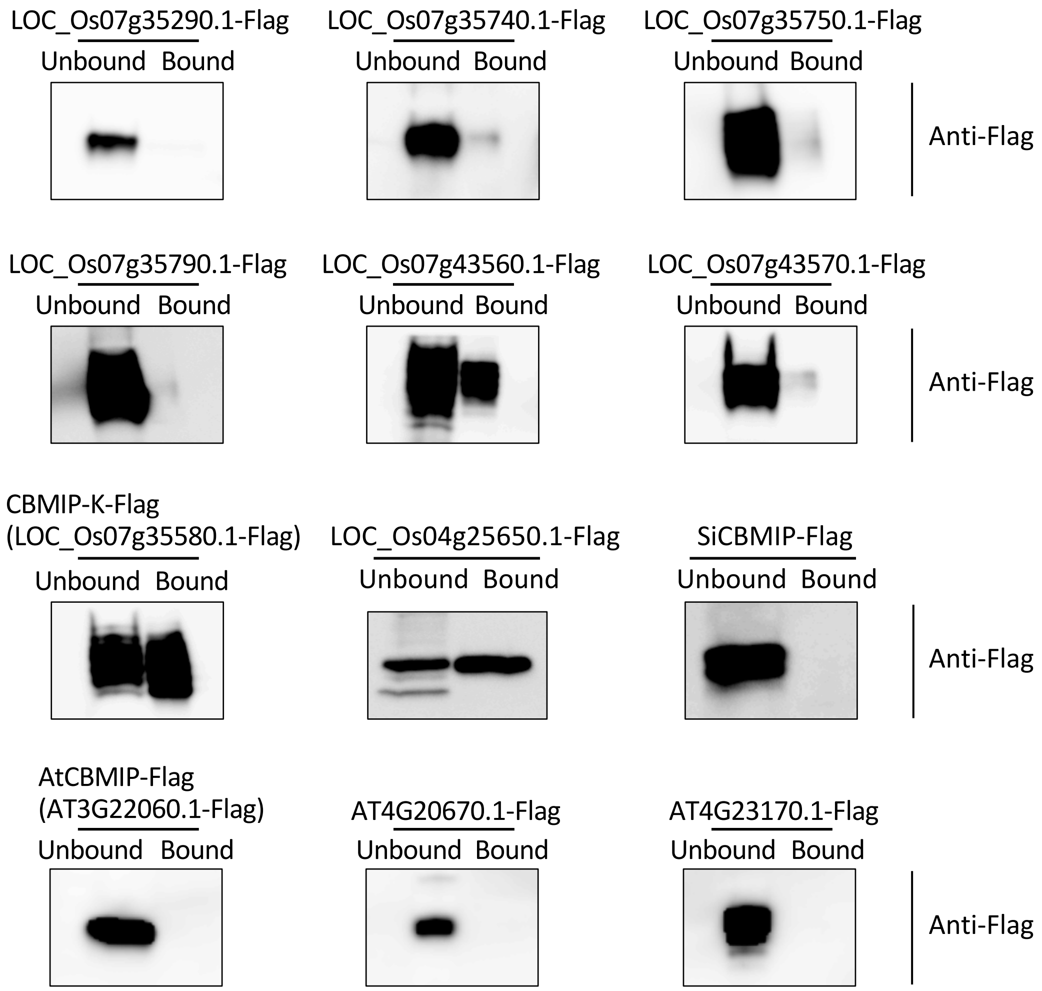


**Fig. S15. Binding assay of various DUF26-containing proteins to mannose.**
Flag-tagged DUF26 domains from *O. sativa* CRKs, CRRSP encoded by *LOC_Os04g25650.1*, SiCBMIP and Arabidopsis CRRSPs were expressed in *N. benthamiana* leaves and assayed for binding to mannose-agarose. Fractions unbound and bound to mannose-agarose were subjected to immunoblot analysis using an anti-Flag antibody.

Fig. S16.

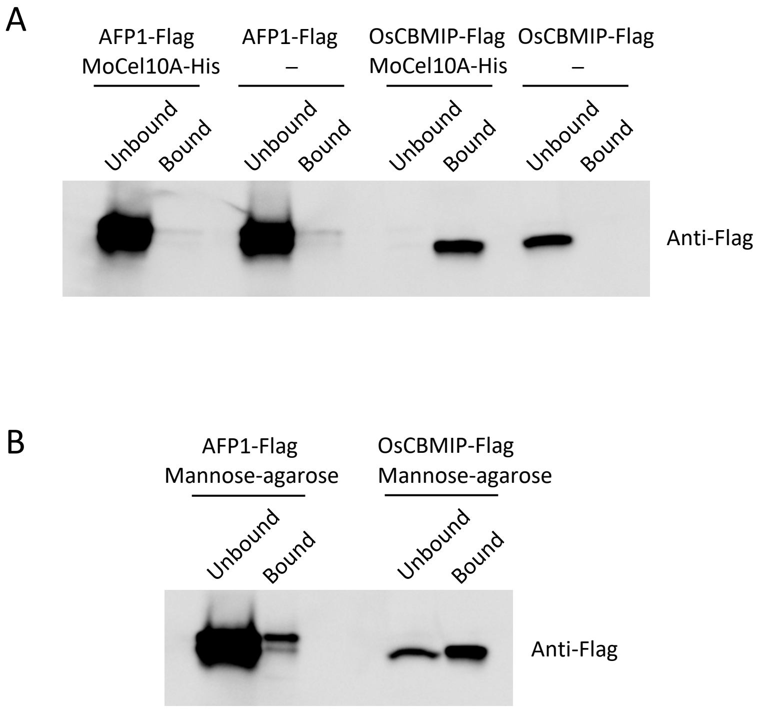


**Fig. S16. Binding assay of AFP1 to CBM1 and mannose.**
Flag-tagged AFP1 (AFP1-Flag) and OsCBMIP (OsCBMIP-Flag) expressed in *N. benthamiana* were assayed for binding to MoCel10A-His (A) and mannose-agarose (B). Fractions unbound and bound to His-resin or mannose-agarose were subjected to immunoblot analysis using an anti-Flag antibody.

Table S1. Amino acid sequence identities among four CBM1-binding proteins

| | OsCBMIP | OsCBMIP-K* | SiCBMIP | AtCBMIP |
|---|---|---|---|---|
| OsCBMIP | - | | | |
| OsCBMIP-K | 43% | - | | |
| SiCBMIP | 27% | 28% | - | |
| AtCBMIP | 29% | 31% | 30% | - |

*DUF26 domain was used for comparing amino acid identity.

Table S2. Summary of the number of CBM families

| * CBM Family | 1 | 2 | 5 | 6 | 9 | 13 | 18 | 20 | 21 | 22 | 32 | 35 | 42 | 43 | 45 | 46 | 48 | 49 | 50 | 52 | 53 | 57 | 63 | 66 |
|---|---|---|---|---|---|---|---|---|---|---|---|---|---|---|---|---|---|---|---|---|---|---|---|---|
| ***Magnaporthe oryzae*** | **28** | - | - | 2 | - | - | 53 | 3 | 1 | - | - | 3 | 2 | 2 | - | - | 3 | - | 14 | 1 | - | - | 1 | 1 |
| ***Fusarium graminearum PH-1*** | **13** | - | - | 1 | - | 2 | 34 | 2 | 2 | - | 3 | 3 | 1 | 1 | - | - | 3 | - | 19 | - | - | - | 3 | - |
| ***Xylanimonas cellulosilytica DSM 15894*** | **-** | 8 | 8 | - | 2 | - | - | - | - | 6 | - | - | - | - | - | 1 | 4 | - | - | - | - | - | - | - |
| ***Xanthomonas oryzae AXO1947*** | **-** | 1 | - | - | - | - | - | - | - | - | - | 1 | - | - | - | - | 5 | - | 2 | - | - | - | 1 | - |
| ***Oryza sativa*** | **-** | - | - | - | - | - | 15 | 6 | - | 9 | - | - | - | 55 | 4 | - | 21 | 5 | 14 | - | 6 | 25 | - | - |
| ***Arabidopsis thaliana*** | **-** | - | - | - | - | - | 10 | 4 | - | 17 | - | - | - | 62 | 6 | - | 16 | 3 | 1 | - | 3 | 4 | - | - |

*** CBM Family; Classification based on amino acid similarity**

Table S3. Summary of binding ability of various DUF26-containing proteins to CBM1 and mannose

| | CBM1 | Mannose | Note |
|---|---|---|---|
| OsCBMIP (LOC_Os04g56430.1) | +++ | +++ | See Fig.1, Fig. 4 |
| LOC_Os04g25650.1 | - | +++ | See Fig.4 |
| LOC_Os07g35290.1 | - | - | See Fig. S13, Fig. S15 |
| OsCBMIP-K (LOC_Os07g35580.1) | ++ | +++ | See Fig. 3, Fig. S15 |
| LOC_Os07g35740.1 | - | - | See Fig. S13, Fig. S15 |
| LOC_Os07g35750.1 | - | - | See Fig. S13, Fig. S15 |
| LOC_Os07g35790.1 | - | - | See Fig. S13, Fig. S15 |
| LOC_Os07g43560.1 | - | ++ | See Fig. S13, Fig. S15 |
| LOC_Os07g43570.1 | - | - | See Fig. S13, Fig. S15 |
| AtCBMIP (AT3G22060.1) | +++ | - | See Fig. S12, Fig. S15 |
| AT4G20670.1 | - | - | See Fig. S12, Fig. S15 |
| AT4G23170.1 | - | - | See Fig. S12, Fig. S15 |
| SiCBMIP (XM_004960453.3) | +++ | - | See Fig. 3, Fig. S15 |
| AFP1 (NM_001320826.1) | - | + | See Fig. 16 |

Table S4. DNA sequences of PCR primers used in this study

| | | |
|---|---|---|
| *OsCBMIP* | Forward; | 5′-ATGGCGCGGTGCACTTTGCTCGTTC-3′ |
| | Reverse; | 5′-CTACTCACGCAGCACCACCATCGGGGCG-3′ |
| *SiCBMIP* | Forward; | 5′-ATGATTCTGCATGCAAGCAAAACAAGCC-3′ |
| | Reverse; | 5′-CTAGCTGTGGTGGTTCAAGATGACTTTC-3′ |
| *AtCBMIP* | Forward; | 5′-ATGTCTTCATTAAAACGCATCGTTTG-3′ |
| | Reverse; | 5′-TCAAGCAGTCTTAACAAAAGGGTAAATCTC-3′ |
| *OsCBMIP-K* | Forward; | 5′-ATGTGCCGCGGATCAGTCCTCCTCCTCC-3′ |
| | Reverse; | 5′-TTATCTCGGCTCGAGCTCAGTGATGGACACC-3′ |
| *OsCBMIP* for qPCR | Forward; | 5′-CATCGCGAGGAAGCTGTTCGCG-3′ |
| | Reverse; | 5′-CTCGAGGCAGCCGCGGCACGCC-3′ |
| *AT4G20670.1* | Forward; | 5′-ATGAAAAAACACACACACCAAAAATATC-3′ |
| | Reverse; | 5′-TTAATATGGAGCTCTCATTGTACTGTTGCTGTAG-3′ |
| *AT4G23170.1* | Forward; | 5′-ATGTCTTCTCTGATCTCTTTTATCTTC-3′ |
| | Reverse; | 5′-TCAATTTGCTCTGCTTGGCGACGGTGGTGG-3′ |
| DUF26-domain of *LOC_Os07g35580.1* | Forward; | 5′-ATGTGCCGCGGATCAGTCCTCCTCCTCC-3′ |
| | Reverse; | 5′-CTTCCCTGCCGGCGCCGGCGCCGGCG-3′ |
| DUF26 domain of *LOC_Os07g35290.1* | Forward; | 5′-ATGGCGCGCCACCGCGTACGCTGCCTC-3′ |
| | Reverse; | 5′-CCTGGGAGACAATTGTGTCACGGTTTG-3′ |
| DUF26 domain of *LOC_Os07g35740.1* | Forward; | 5′-ATGCGGCGGCGCTCCACGTTCGCCGTTC-3′ |
| | Reverse; | 5′-TCTAGGAGACAATTGTGTAACGGTTTG-3′ |
| DUF26 domain of *LOC_Os07g35750.1* | Forward; | 5′-ATGCGGCGGCGCTCGTCGCTCGTCCAC-3′ |
| | Reverse; | 5′-TCTAGGAGACAATTGTGTAACGGTTTG-3′ |
| DUF26 domain of *LOC_Os07g35790.1* | Forward; | 5′-ATGCGGCGGCGCTCGTCGCTCGTCCAC-3′ |
| | Reverse; | 5′-TCTAGGAGACAATTGTGTAACGGTTTG-3′ |
| DUF26 domain of *LOC_Os07g43560.1* | Forward; | 5′-ATGTTGAGCTGTAGTGGCAACATGCTG-3′ |
| | Reverse; | 5′-GGGAGTTCCCGGCACCGCCGGCGGC-3′ |
| DUF26 domain of *LOC_Os07g43570.1* | Forward; | 5′-ATGCACGGCGGCGGCGCCCGTGCGCTC-3′ |
| | Reverse; | 5′-GCCTTCCATGGCAACCTGCAGGTGC-3′ |
| 311 bp for RNAi of *OsCBMIP #1* | Forward; | 5′-CACCGCCGCCTACGACCGCGCGGTGACGCG-3′ |
| | Reverse; | 5′-GCACCACCATCGGGGCGCCGGTGTAG-3′ |
| 433 bp for RNAi of *OsCBMIP #2* | Forward; | 5′-CACCGCCATGCGTGAAGCTAGAATAAAATGGC-3′ |
| | Reverse; | 5′-TGGATCCAATTCTATTTAGTCTGACTC-3′ |
| *Magnaporthe actin* gene (XM_003719823.1) | Forward; | 5′-AACGCCCCCGCTTTCTAC-3′ |
| | Reverse; | 5′-GGTACGACCCGAAGCGTAAA-3′ |
| Rice *actin* gene (*LOC_Os03g61970*.1) | Forward; | 5′-AACTGGGACGACATGGAGAAA-3′ |
| | Reverse; | 5′-AGCAACACGCAGCTCATTGT-3′ |
| *Hygromycin* gene | Forward; | 5′-CGACAGCGTCTCCGACCTGA-3′ |
| | Reverse; | 5′-CGACCTCGTATTGGGAATCC-3′ |

Table S5. Amino acid sequences of DUF26-A and DUF26-B used for phylogenetic tree construction

Amino acid sequences of DUF26-A

>AT1G04520.1_DUF26-A
QQFSDPSGLYSQALSAMFGSLVSQSTKTRFYKTTTGTSTTTITGLFQCRGDLSNHDCYNCVSRLPVLSDKLCGKTIASRVQLSGCYLLYE
>AT1G19090.1_DUF26-A
SSITDVSPPIYVFLQCREDLSVSDCRHCFNESRLELERKCSGSGGRIHSDRCFLRFDD
>AT1G61750.1_DUF26-A
CYTSGNYTPNSSYKSNLDTLISVLDSQSSNKGFYSYASGSSPTTTVYGSYLCRGDISSSTCETCISRASKNVFIWCPVQKEAIIWYEECFLRYSS
>AT1G63550.1_DUF26-A
CNPTNNFTQTSSYETNRDTLLASLRESSSLGHYSNATEGLSPDTVHGMFLCRGDITTASCVDCVQTATTEIASNCTLNKRAVIYYDECMVRYSN
>AT1G63570.1_DUF26-A
CSVDSFTQTSSYETNRNILLTTLSLTSSLVHYLNATIGLSPDTVYGMFLCRGDINTTSCSDCVQTAAIEIATNCTLNKRAFIYYDECMVRYSN
>AT1G63580.1_DUF26-A
PDTYQSNRNTVLSTLRNHSSLGSYYFNATAGLSPNTVYGMFLCIGNISKTSCSNCVHSATLEMDKSCESHDTSFMFSDECMVRYSD
>AT1G63590.1_DUF26-A
CTQFDNVTRTSSYLSNRDTVLSTLRNRSSIGSYSNATAGLSPNTIYGMFLCRGDLNRTSCSDCVNATTLEIYKSCFYRKSALVISNECIVRYSN
>AT1G63600.1_DUF26-A
CNQFSDNFTQTSTYETNRETVLSSLRLRSSLGSYSNATAGISPDTVRGMFLCRGDISETSCSDCVQTATLEISRNCTYQKEAFIFYEECMVRYSD
>AT1G70520.1_DUF26-A
ETAYVPNFVATMEKISTQVQTSGFGVALTGTGPDANYGLAQCYGDLPLNDCVLCYAEARTMLPQCYPQNGGRIFLDGCFMRAENY
>AT1G70530.1_DUF26-A
CNNRTTTPQQRSLFVTNFLAAMDAVSPLVEAKGYGQVVNGTGNLTVYAYGECIKDLDKKDCDLCFAQIKAKVPRCLPFQKGTRGGQVFSDGCYIRYDDY
>AT1G70690.1_DUF26-A
CSPAKFSPSSGYETNLNSLLSSFVTSTAQTRYANFTVPTGKPEPTVTVYGIYQCRGDLDPTACSTCVSSAVAQVGALCSNSYSGFLQMENCLIRYDN
>AT2G01660.1_DUF26-A
CSQEKYFPGSPYESNVNSLLTSFVSSASLYTYNNFTTNGISGDSSSVYGLYQCRGDLSSGSGDCARCVARAVSRLGSLCAMASGGALQLEGCFVKYDN
>AT2G31620.1_DUF26-A
CINSEGKYKAKNSYESRLKDHLDSMSNILDYGFIHGVGGADSSTYYIKAQCRGDASESKCRSCLFTAFSGILRRCPNNRGRIIWYDNCFLYIS
>AT2G33330.1_DUF26-A
PSGLYSQALSAMYGLLVTQSTKTRFYKTTTGTTSQTSVTGLFQCRGDLSNNDCYNCVSRLPVLSGKLCGKTIAARVQLSGCYLLYE
>AT3G21960.1_DUF26-A
CINGEGTFKSGSPYEKEIKQLIDFLSSFIKDYSFVHGVSGIGPDDINVKFQCRGDTLQAKCRSCLATAFSEIRSKCPNNKGRIIWYDNCHLDL
>AT3G21970.1_DUF26-A
CNNTQGRYTHGSTFEKNLNQVLHNISNLDLRYGYAYVSNVVAYKVSKDPNIVFVLLQCRGDSFGSKCHSCLSTAVSGLRERCPGNRGATIWYDQCLLEIS
>AT3G21980.1_DUF26-A
KCNNTEGKYSHGSAFEKYINLALRAIDSDNYLNGFAYIERGEDPNKVFVMYQCRGDSYGSKCKSCISAAIS
>AT3G21990.1_DUF26-A
KCSNTQGKYKQGSAFEKNLNLVLSTITSIGNFRDGFRYTEEGEDPNNVFVMFQCRGDSYWSKCPPCISTAVSGLRRRCPRNKGAIIWYDQCLLKIS
>AT3G22040.1_DUF26-A
CRVGQGKYHPGSRYEKDFDSLTSFVAANKFIDGFVHSSNSDGPNSTTIIFQCRGDSYKSNCRTCYDTALAGFRKRCPRNKGGIIWYDQCFLDVS
>AT3G22060.1_DUF26-A
CSDIEGSFTSKSLYESNLNNLFSQLSYKVPSTGFAASSTGNTPNNVNGLALCRGDASSSDCRSCLETAIPELRQRCPNNKAGIVWYDNCLVKYSS
>AT3G29040.1_DUF26-A
CNNTQGTYTRGSTFEKNLNQVIRNISHLHLRYGYTYNSNVEAYEVSKDPNIVFVLLQCRGDSYGSKCHSCLHTAFSGLRERCRGNKGAIIWYDQCVLEIS
>AT3G45860.1_DUF26-A
SSILTYSRNSTYFTNLKTLLSSLSSRNASYSTGFQTATAGQAPDRVTGLFLCRGDVSQEVCRNCVAFSVKETLYWCPYNKEVVLYYDECMLRYSH
>AT3G60720.1_DUF26-A
CSPEKYTPNTPFESNRDTFLSSVVTSSSDASFNSFAVGNDSSSSSSSAVFGLYQCRDDLRSSDCSKCIQTSVDQITLICPYSYGASLQLEGCFLRYET
>AT4G00970.1_DUF26-A
CLSQQSNFAKSSQFSKNLDSLVSSIPSLKSNTYNFYSLSVGSISDQERVEAIGICNRVVNRVDCLNCIAQAAVNLTTMYCPQHRGAYVRATKCMFRYSD
>AT4G04490.1_DUF26-A
EDFSPNTSYVENLESLLPSLASNVIRERGFYNVSLDGVYALALCRKHYEVQACRRCVDRASRTLLTQCRGKTEAYHWDSENDANVSCLVRYSN
>AT4G04500.1_DUF26-A
NYGVSRTYLFSSLPSNVVSNGGFYNASFGRDSKNNRVHVVALCRRGYEKQACKTCLEHVIEDTKSKCPRQKESFSWVTDEFDDVSCSLRYTN
>AT4G04510.1_DUF26-A
SFPTNSSYQKNRDSLFSTLSDKVTTNGGFYNASLDGVHVVGLCRRDYDRQGCINCVEESIRQIKTSCSNRVQSFHCNSDDRERVSCLVRT
>AT4G04540.1_DUF26-A
SFFNGNSSYAQNRRDLFSTLPNKVVTNGGFYNSSLGKSPNIVHAVALCGRGYEQQACIRCVDSAIQGILTTTSCLNRVDSFTWDKDEEDNVSCLVSTS
>AT4G04570.1_DUF26-A
FNGNSSTFAQNRQKLFPTLADKVIINDGFYNASLGQDPDKVYALVSCARGYDQDACYNCVQSLTQNTLTDCRSRRDSFIWGGNDDVTCLVRSSN
>AT4G05200.1_DUF26-A
CPNTTTYSRNSSYLTNLRTVLSSLSSPNAAYASLFDNAAAGEENDSNRVYGVFLCRGDVSAEICRDCVAFAANETLQRCPREKVAVIWYDECMVRYSNQ
>AT4G11460.1_DUF26-A
CSEKFGTFTPGGTFDKNRRIILSSLPSEVTAQDGFYNASIGTDPDQLYAMGMCIPGAKQKLCRDCIMDVTRQLIQTCPNQTAAIHWSGGGKTVCMARYYN
>AT4G11470.1_DUF26-A
VFFRPNGNYDTNRRLVLSTLASNVSSQNNRFYNVSVGEGAGRIYALGLCIPGSDPRVCSDCIQLASQGLLQTCPNQTDSFYWTGDNADKTLCFVRYSN
>AT4G11480.1_DUF26-A
FFRPNGTYDTNRHLILSNLASNVSSRDGYYNGSVGEGPDRIYALGLCIPGTDPKVCDDCMQIASTGILQNCPNQTDSYDWRSQKTLCFVRYSN
KTSRAGSGGAPTAYGRATCKQSISQSDCTACLSNLVNRIFSICNNAIGARVQLVDCFIQYEQ

>AT4G11490.1_DUF26-A
CNETGYFEPWKTYDTNRRQILTSLASKVVDHYGFYNSSIGKVPDEVHVMGMCIDGTEPTVCSDCLKVAADQLQENCPNQTEAYTWTPHKTLCFARYSN
>AT4G11521.1_DUF26-A
YFKPNGTYDLNRRRILSSLASKVTAHNGFYSSTIGQNPNQMFIISMCIPGTKPERCSDCIKGSTDGLLRSCPNQTVGYVWPDCCMVRYSN
>AT4G11530.1_DUF26-A
FFKPNSTYDLNRRQILSTLSSNVTSHNGFFNSKFGQAPNRVFINGMCIPGTKPETCSDCIKGASDKISESCPNKTDAYTWPDCCMVRYSN
>AT4G20550.1_DUF26-A
KCSNTQGKYKQGSAFEKNLNLVLSTITSIGNFRDGFRYTEEGEDPNNVFVMFQCRGDSYWSKCPPCISTAVSGLRRRCPRNKGAIIWYDQCLLKIS
>AT4G20560.1_DUF26-A
KCSNTQGKYKQGSAFEKNLNLVLSTITSIGNFRDGFRYTEEGEDPNNVFVMFQCRGDSYWSKCPPCISTAVSGLRRRCPRNKGAIIWYDQCLLKIS
>AT4G20570.1_DUF26-A
KCSNTQGKYKQGSAFEKNLNLVLSTITSIGNFRDGFRYTEEGEDPNNVFVMFQCRGDSYWSKCPPCISTAVSGLRRRCPRNKGAIIWYDQCLLKIS
>AT4G20580.1_DUF26-A
KCSNTQGKYKQGSAFEKNLNLVLSTITSIGNFRDGFRYTEEGEDPNNVFVMFQCRGDSYWSKCPPCISTAVSGLRRRCPRNKGAIIWYDQCLLKIS
>AT4G20590.1_DUF26-A
KCSNTQGKYKQGSAFEKNLNLVLSTITSIGNFRDGFRYTEEGEDPNNVFVMFQCRGDSYWSKCPPCISTAVSGLRRRCPRNKGAIIWYDQCLLKIS
>AT4G20600.1_DUF26-A
KCSNTQGKYKQGSAFEKNLNLVLSTITSIGNFRDGFRYTEEGEDPNNVFVMFQCRGDSYWSKCPPCISTAVSGLRRRCPRNKGAIIWYDQCLLKIS
>AT4G20610.1_DUF26-A
KCSNTQGKYKQGSAFEKNLNLVLSTITSIGNFRDGFRYTEEGEDPNNVFVMFQCRGDSYWSKCPPCISTAVSGLRRRCPRNKGAIIWYDQCLLKIS
>AT4G20620.1_DUF26-A
KCSNTQGKYKQGSAFEKNLNLVLSTITSIGNFRDGFRYTEEGEDPNNVFVMFQCRGDSYWSKCPPCISTAVSGLRRRCPRNKGAIIWYDQCLLKIS
>AT4G20640.1_DUF26-A
KCSNTQGKYKQGSAFEKNLNLVLSTITSIGNFRDGFRYTEEGEDPNNVFVMFQCRGDSYWSKCPPCISTAVSGLRRRCPRNKGAIIWYDQCLLKIS
>AT4G20670.1_DUF26-A
KCSNTQGKYKQGSAFEKNLNLVLSTITSIGNFRDGFRYTEEGEDPNNVFVMFQCRGDSYWSKCPPCISTAVSGLRRRCPRNKGAIIWYDQCLLKIS
>AT4G21230.1_DUF26-A
GNFTSNTSYSLNLNRLISSLPDLTPTINGFYNISINGEVNAIALCRGDVKPNQDCISCITTAAKQLVESCPNIIEANIWLEKCMFRYT
>AT4G21400.1_DUF26-A
CVASGGNFTANSSFAGNLNGLVSSLSSLTSKPYGFYNLSSGDSSGERAYAIGLCRREVKRDDCLSCIQIAARNLIEQCPLTNQAVVWYTHCMFRYSN
>AT4G21410.1_DUF26-A
CVDRGNFTANSTFAGNLNRLVSSLSSLKSQAYGFYNLSSGDSSGERAYAIGLCRREVKRDDCVSCIQTAARNLTKQCPLTKQAVVWYTHCMFRYSN
>AT4G23130.1_DUF26-A
CTNRISRNSIYFSNLQTLLTSLSSNNAYFSLGSHSLTKGQNSDMVFGLYLCKGDLSPESCRECVIFAAKDTRSRCPGGKEFLIQYDECMLGYSD
>AT4G23140.1_DUF26-A
CPNTTTYSSNSTYSTNLRTLLSSLSSRNASYSTGFQNATAGKAPDRVTGLFLCRGDVSPEVCRNCVAFSVNQTLNLCPKVREAVFYYEQCILRYSH
>AT4G23150.1_DUF26-A
CPNATTYSSNSTYLTNLKTLLSSLSSRNASYSTGFQNATVGQALDRVTGLFLCRGDVSPEVCRNCVTFAVNNTFSRCPNQREAVFYYEECILRYSH
>AT4G23170.1_DUF26-A
CPNTTTYSRFSTYSTNLRTLLSSFASRNASYSTGFQNVTVGQTPDLVTGLFLCRGDLSPEVCSNCVAFSVDEALTRCPSQREAVFYYEECILRYSD
>AT4G23180.1_DUF26-A
CQNTANYTSNSTYNNNLKTLLASLSSRNASYSTGFQNATVGQAPDRVTGLFNCRGDVSTEVCRRCVSFAVNDTLTRCPNQKEATLYYDECVLRYSNQ
>AT4G23190.1_DUF26-A
CTTDKGTFRPNGTYDVNRRLILSSLPSNVTDQDGLYYNGSIGQQPNRVYAIGMCIPGSTSEDCSDCIKKESEFFLKNCPNQTEAYSWPGEPTLCYVRYSN
>AT4G23200.1_DUF26-A
TYFIPNSTYDTNRRVILSLLPSNVTSHFGFFNGSIGQAPNRVYAVGMCLPGTEEESCIGCLLSASNTLLETCLTEENALIWIANRTICMIRYSD
>AT4G23210.1_DUF26-A
CIENRKYFTPNGTYDSNRRLILSSLPNNTASRDGFYYGSIGEEQDRVYALGMCIPKSTPSDCSNCIKGAAGWLIQDCVNQTDAYYWALDPTLCLVRYSN
>AT4G23260.1_DUF26-A
CDNTTGTFIPNSPYDKNRRLILSTLASNVTAQEGYFIGSIGIAPDQVFATGMCAPGSERDVCSLCIRSTSESLLQSCLDQADAFFWSGEETLCLVRYAN
>AT4G23270.1_DUF26-A
CSVTTTFSSNSTYSTNLKTLLSSLSSLNASSYSTGFQTATAGQAPDRVTGLFLCRVDVSSEVCRSCVTFAVNETLTRCPKDKEGVFYYEQCLLRYSN
>AT4G23280.1_DUF26-A
CSITTTYSSNSTYSTNLKTLLSSLSSRNASYSTGFQNATAGQAPDMVTGLFLCRGNVSPEVCRSCIALSVNESLSRCPNEREAVFYYEQCMLRYSN
>AT4G23300.1_DUF26-A
CIENRKYFTPNGTYDSNRRLILSSLPNNTASQDGFYYGSIGEEQDRVYALGMCIPRSTPSDCFNCIKGAAGWLIQDCVNQTDAYYWALDPTLCLVRYSN
>AT4G28670.1_DUF26-A
CNNGTVSNEEAYRRSYQINLDAIRGDMRHVKFGTHEHGDPPERMYVLSQCVSDLSSDECSLCWSRATDLLSQCFPATGGWFHLDGCFVRADNY
>AT4G38830.1_DUF26-A
CSNVTGNFTVNTPYAVNLDRLISSLSSLRRNVNGFYNISVGDSDEKVNSISQCRGDVKLEVCINCIAMAGKRLVTLCPVQKEAIIWYDKCTFRYSN
>AT5G37660.1_DUF26-A
CSQQKFSPASAYESNLNSLLTSLVNSATYSSYNNFTIMGSSSSDTARGLFQCRGDLSMPDCATCVARAVSQVGPLCPFTCGGALQLAGCYIKYDN
>AT5G40380.1_DUF26-A
TFVEDMHSLSLKLTTRRFATESLNSTTSIYALIQCHDDLSPSDCQLCYAIARTRIPRCLPSSSARIFLDGCFLRYETY
>AT5G41290.1_DUF26-A
CMKSSRNTTSNTTYNKNLNTLLSTLSNQSSFANYYNLTTGLASDTVHGMFLCTGDVNRTTCNACVKNATIEIAKNCTNHREAIIYNVDCMVRYSD
>AT5G41300.1_DUF26-A
CIDSSRNTTGNTTYNKNLNTMLSTFRNQSSIVNNYNLTTGLASDTVYGMFLCTGDVNITTCNNCVKNATIEIVKNCTNHREAIIYYIDCMVRYSD

>AT5G41300.1_DUF26-A
CIDSSRNTTGNTTYNKNLNTMLSTFRNQSSIVNNYNLTTGLASDTVYGMFLCTGDVNITTCNNCVKNATIEIVKNCTNHREAIIYYIDCMVRYSD
>AT5G43980.1_DUF26-A
CANQKSPDPTGVFSQNLKNLFTSLVSQSSQSSFASVTSGTDNTTAVIGVFQCRGDLQNAQCYDCVSKIPKLVSKLCGGGRDDGNVVAARVHLAGCYIRYES
>AT5G48540.1_DUF26-A
CNTNSNISASSQVSKNIDSLLATLVSKTPSKGFKTTTSSSYNNKEKVYGLAQCRGDISNTDCSTCIQDAAKKIREVCQNQSDSRILYDFCFLRYSQ
>EFJ11427_DUF26-A
AQMGRNLLTVLATLQDQGPAAGGYKSTSFGSMSDTVYGFIQCRGDISAQECSSCARSAVQILNSTCVRNSTKGGRIHPGNCFLRYED
>EFJ16458_DUF26-A
CGEEKFTNGSSYSTALDNLLATLITNAPVTGFFQSSVQSAELTLYGLLQCRQDMSRDSCDQCARDIGQAVKSRCSSSLGAKIQLYGCFLRYEN
>EFJ17937_DUF26-A
CGEEKFTNGSSYSTALDNLLATLITNAPVTGFFQSSVQSAELTLYGLLQCRQDMSRDSCDQCARDIGQAVKSRCSSSLGAKIQLYGCFLRYEN
>LOC_Os01g23970.1_DUF26-A
CNATAGNHTAVGSAYLSNLRALGGALSRRALATGFASGSYGAAPDEVHGLVLCRGDFTGGNCTDGLASAFRDAAAQFCPGAADATVYYDQYMIRYTN
>LOC_Os01g26390.1_DUF26-A
CQDSRGKYTSNSTYQANIQSLSSTLPAKAAAPSTGLFATRVAGNAPDTVYALAFCRGDITNASACAGCVASGFQDAQQLCPFNKAASLYYDLCLLRFAD
>LOC_Os01g36790.1_DUF26-A
MDDLNSNVSANGFGTSAVGTTAGLNPNAVFGLGQCYRDLSPVDCKLCFAEVRSLLPKCYPSAGGRLYLDGCFGRYANY
>LOC_Os01g38850.1_DUF26-A
CSLSGGKYEPNSTYEANLRALASLLLAEARATAFASDSFGAAPDAVYGIALCRGDYAGDACAGGLRKAFRDAIDHGVFCAGFRDVTVYYDEHMFRFS
>LOC_Os02g43000.1_DUF26-A
CSPSKYEPNTAFQSNLNSLLSSIASTASSGAAYNSFTAGGGAGPDPAAGTAAYGLYQCRGDLSPGDCVACVRQTVARLGAVCANAYAASLQVDGCYVRYD
>LOC_Os02g50200.1_DUF26-A
CSQGRYASGTQYASDVDSVLTSVANSAPYSPYANFTSPTSNSVVGVYQCRSDLPASVCTGCVRSAISRLSSLCAWATGGAVQLRACFVRYGN
>LOC_Os03g16950.1_DUF26-A
AGNSKAVASINSVLTDLVAKGSTGGGFATSSAGKANNVIYGLAQCRGDVSTSDCQACLASAANQILTSCNYQSDSRIWYDYCFMRFEN
>LOC_Os03g16960.1_DUF26-A
SKAVASINSVLTDLVTKGSTGVGFATSTAGKGNNVIYGLVQCRGDVSTSDCQACLASAANQILTSCNYQSDSRIWYDYCFMRFEN
>LOC_Os03g36650.1_DUF26-A
CSATDSFAADSSFAGNLGRLVSLLEAKAPAIGFDIATVGVGGDGEDQRVHGLALCRGDVARATCAECIRAAGALARRVCPSKKDAVVWLDACMLRYS
>LOC_Os04g09780.1_DUF26-A
CSPQNYTAGSAYGTSLRGVLKDVVDAAVSGGGYAVANDAGGAAHGLAICYADAPPEVCRLCLAMAAGNLSLACPRAVGGAMLYNNCLLRYA
>LOC_Os04g25060.1_DUF26-A
CDTSAGTYKAGSAYESNLRDLAAALRADAAASPSALFATGNRGGAPDAVYGLLLCRGDLSVSDCFDCGTRVLADVGRVCGGRHGRAKDVALVYNQCYARFSN
>LOC_Os04g25650.1_DUF26-A
CDTSAGTYKAGSAYESNLRDLAAALRAGAAASPSALFATGIRGGAPDAVYGLLLCRGDLSVSDCFDCGTRVLADVGRVCGGRHGRAKDVALVYNQCYARFSN
>LOC_Os04g45460.1_DUF26-A
CSPSKYQPGTPFEGNLNSLLASIANAAPNGGYNSFTAGSNGTGDGAAAYGLYQCRGDLGNADCAACVRDAVGQLNEVCAAAYAASLQLEGCYVRYDS
>LOC_Os04g56430.1_DUF26-A
CGDGTYEQGSAYENNLLNLALTLRDGASSQEILFSTGSNGAAPNTVYGLLLCRGDISRAACYDCGTSVWRDAGSACRRAKDVALVYNECYARLSD
>LOC_Os05g02200.1_DUF26-A
QASVAQVLSELVPRASAGYYATATAGRGGDGSAIWGLAQCRGDIPAPDCALCASAAARQLAGACRGRADARVWYDYCFARYDD
>LOC_Os05g03920.1_DUF26-A
ADQEGFDVSFVNTLELIYQNVTRSGFGAAASGEGADTVYGLGQCMGYLSPTDCQLCYAQSRVKLPHCLPATGGRIYLDGCFLRYG
>LOC_Os05g41370.1_DUF26-A
CPTTSSTNSSHVDDGAFGANLRALLSSLSAAAAASSSGFAENATGAAPDTAYGLAQCRGGIVGGGNGTSCRSCLDDSVRDAAKACPGEKSAVIISDYCLVRYSN
>LOC_Os06g14280.1_DUF26-A
CSQARYDAGTQYAADVDTALSALTNSAGYTAYANYTSPSAASGTGLVGVYQCRSDLPAAICGGCVRSAATKLASLCNSAAGAGVQLRACFVRYGN
>LOC_Os07g30410.1_DUF26-A
CGSSGNYTAGSKYQANLQALAATLPSTASSSSPALFAKDAAGGGDAEPDRVFALTLCRGDTASANASSSSCADCASRAFRDAQSVCPYSKEVAVYYDPCLLYFS
>LOC_Os07g30510.1_DUF26-A
CGSSGNYTAGSKYQANLQALAATLPSTASSSSPALFAKDAAGGGDAEPDRVFALTLCRGDTASANASSSSCADCASRAFRDAQSVCPYSKEVAVYYDPCLLYFS
>LOC_Os07g34980.1_DUF26-A
CSNANTITHMPEGTYKTNLLQLAKNLITNVNQTQLHSANGTAGAAGPDTVYGAVLCRGDSSAESCATRLQRVLDTAS
>LOC_Os07g35004.1_DUF26-A
CNTTAARRTYLPNSTFEANLNGLFAVLSRNASASGYAAGAFGAAPDTAYGLLLCRGDFTGNDCSAARLASSFQQAASSCLYSKDVAVYYDQYQLRYSDQ
>LOC_Os07g35140.1_DUF26-A
CNGTRGNFTEGSAFGLNLELLAAELPANASSSRSLFASAAVGAAAAPEDRVFGLALCRGDMRDAAACAGCVSGAFQRLRALCGRDRDATYYHDLCVVRYS
>LOC_Os07g35280.1_DUF26-A
SGDVFAPRSTYQSNLALLSAGLAKNASASPALFAAGGVGDPPDTVYGLALCRGDTTNATACGACVAAAFQDGQQLCAYAREATVFYDPCYLRFS
>LOC_Os07g35290.1_DUF26-A
CGQSGNFSANSAYQSNLRQLSATLPKNASAALFAAGSLGTVPDIVYALALCRGDANASACESCVDNAFQGGQQLCPYNKDVFIVYDLCYLRFTN
>LOC_Os07g35300.1_DUF26-A
CGQGGNYSANGTYQSNLAGLSATLPKNASASRTLFAKDSLGAVPDIVYALALCRGDVANATACESCVATAFQDAQQLCPYDKDAFIVYDLCYLAFSN
>LOC_Os07g35310.1_DUF26-A
CGTTGNFTANSTYQANLDAVAAALPRNISSSPDLFATAMVGAVPEQVSALALCRGDANATECSGCLATAFQDVQNMCAYDKDAAIYYDPCILYYSN
>LOC_Os07g35330.1_DUF26-A
CASGAYAANSTYEANLAVLAAALPGNASTAAAAGYATATVGAVPDQVSALALCRGDANATACRACVAASFRVARRDCPSSKDATTYQDGCIVRFSD

>LOC_Os07g35340.1_DUF26-A
NTYVGNSTFEANLNHLAAELPGNVSTAHTGGFAVATVGADPDQVFALALCRGDVNATACRACVAAAFVDGKNACPGINGVTVYEDACVVRFS
>LOC_Os07g35380.1_DUF26-A
CDEGVGNTYVANSTFEANLNVLAAALSPNVSVAPAGFAVATVGADPDKVFAMALCRGDVNASACSACVAAAFVDGKKDCPGNSGVAMYEDACVARFSRY
>LOC_Os07g35390.1_DUF26-A
CNDTAGEFPARRSSYLASINLIAATLPGNASASPDLFATAEGVGAPPDQVSALALCRGDANASTCLACLTQAFLDLPNACAYHKVAAIFYDSCLLAYSN
>LOC_Os07g35410.1_DUF26-A
CDMGVATYAANSTFEANLDRLGAELPANVSAARATGGYAVATVGAAPDLVYALALCRGDVNASACGACVAAAFADGKRSCPGIKGATVSGPGDGCVLRYS
>LOC_Os07g35540.1_DUF26-A
TNVTGNSAFDRNLGLLAAALAANASAAGAPGFAVRTAGAAPDQVYALALCRGDVNASACRACVAAAFVDAKGVCPGGISLYEDACLIRFT
>LOC_Os07g35580.1_DUF26-A
CGENGNYTANSTYQANLKQLAAALHKNVSSGTGGGRLFASGAVGAVPDAVYALALCRGDINASACADCVGTIFQDAQQLCPYRKEVSIVYDSCYLRFSN
>LOC_Os07g35640.1_DUF26-A
CNATAGNYTEGSAYQANVRALASALPANASSSRALFAEGAAGTAPDKVYAIALCRGDTNASSCAACLAAAFDTAQQLCAFNRRATLFNDPCILRYSDQ
>LOC_Os07g35650.1_DUF26-A
CDPYSASGRYSENSTFQANVNRLSATLPRNTSSSPAMYATGAAGDVPDKVYGYALCRGDVADAHACERCVAAALRDAPRVCPLVKDALVFHDLCQLRYSN
>LOC_Os07g35660.1_DUF26-A
CDTAGGNYTEGSTYQANVRALASALPVNASSSRALFAKGAAGAAPDVVYAIALCRGDTNASSCAACVATAFQDAQQLCAFNRRATMFDDPCILRYSDQ
>LOC_Os07g35690.1_DUF26-A
GNYSKNGTYQVNLDLLSTTLPKNTSSSPAMYATGTVGDVPDKVYGLALCRGDANASACERCVAAALRDAPRRCPLVKDVLVFYDLCQLRYSN
>LOC_Os07g35700.1_DUF26-A
CGDSGNYTEHGTYHANIQYLATSLPSYASSSPSLFASGSSGTVPDAIYALALCRGDTNSSSCATCVAAAIQSAQELCPLVKTVIVYDDTCILRFAN
>LOC_Os07g35740.1_DUF26-A
GTYTANSTYDTNLQSLIAALQQNASTSPTLFAAGALGAAPDAVYGLILCRGDVSSSDCYDCGTRAGQDVAPACNRTRDAILVYNQCYTRFS
>LOC_Os07g35750.1_DUF26-A
CGTGGTYAANSTYETNLLDLISALQGNASSSPTLYASGAVGSGGRDAVYGVMLCRGDLSTSDCNDCGTRAGQDVGRVCNRTRDAALVYNQCYVRVSD
>LOC_Os07g35790.1_DUF26-A
CGTSGGNYTAGSTYESNLLRLASTLRANASASPTLFASGVRGAGPDAVYGLLLCRGDMNPSDCFDCGTRVGDDVAQACNRTKDAILVYNQCYAQFSD
>LOC_Os07g35810.1_DUF26-A
CGTSGGNYTAGSTYESNLLRLASTLRANASASPTLFASGVRGVGPDAVYGLLLCRGDMNPSDCFDCGTNVWRDAGPTCNRTKDAILVYNQCYAQFSD
>LOC_Os07g43560.1_DUF26-A
CNGSSNYTANSAFQRNLGVLAAALPGNASTSPDLLANATVGGAPDTVYALAFCPPIDNQNASGCRACVASAFADARSLCPNNRGAHIIYDGCVLTFS
>LOC_Os07g43570.1_DUF26-A
CSGRRYAANSSFDASLQQVARTLPGNASSSPLLFATLAVAGEAYALALCQGGTSAGSCNYCVAQTMRDGEHACAGDADVAMYDDICTVRFSD
>LOC_Os07g47230.1_DUF26-A
CPSTADGTYAPNSTYQSNLAALAAELIENSTEYGSAAGSFGAAPDAVYGVALCRGDSKGPLCAGYLREAFDAAMNRTTSSRPLCELRRNVTLFYDRFQLRFA
>LOC_Os08g04210.1_DUF26-A
INFVVSDLVAKARTGGGFATSKAGRGYDAFYGLAQCRGDVSGGDCDACLAQAAKQMVSYCNYTSDSRLWYEYCFMRYDNY
>LOC_Os08g04230.1_DUF26-A
RSINSVVSDLAAKARAGGGFATSSAGRGIDAFYGLAQCRGDVSGGDCDACLAQAAKQMVTNCNYTLDSRIWYEYCFMRY
>LOC_Os08g04240.1_DUF26-A
RSVNFVVSDLVAKARTGGGFATSRAGRGYDAFYGLAQCRGDVSGGDCDACLAQAAKQIVSNCNYTSDSRIWYEYCFMRYYNY
>LOC_Os08g04250.1_DUF26-A
RAVNFVVSDLVAKARTGGGFATSKAGRGSEVFYGLAQCRGDVSGGDCDACLAQAAKQMVSNCNYTSDSRIWYEYCFMRYY
>LOC_Os10g04720.1_DUF26-A
CGNGGNYTANGTYQSNLARLAAALPSNASSSPDHFATATAGQAPDAAYALALCRGDVANATACGDCVAASFQDARRTCPSDKSATIYYDDCLLRFA
>LOC_Os10g04730.1_DUF26-A
GVFCDNLKFVSATLPNKTSSSPHHYATAAAGQAPDVVYVLALCRGDLNDTACGESVAYTFARLINESCVANYTAGAYYGDCTGVYS
>LOC_Os10g17950.1_DUF26-A
CNNGSAYAANTTYDTNVHSILATLTARTPNTTTGFATATTGRGTDTEAWGLALCRGDTDRVGCASCLAAVPAVAFNECRGDMDVTVFYDRCLARFS
>LOC_Os10g17960.1_DUF26-A
CNNGSSYAANTTYHSNVRAVLTALSAITPNSTARFATASAGRGGADAVWGLALCRGDTDRAGCASCLAAVPAVAFGECRGDRDVAVFYDRCLARFS
>LOC_Os11g28104.1_DUF26-A
CSQYNATPAAAFLAALNATFAVLRANLSSAGGGGGFATAAEPRAAAPAFAMAQCRPYVAGAGSCAACFDAAASRLRARCGAANGGRAILDGCVVRYES
>LOC_Os11g38850.1_DUF26-A
CGATNYTARSAYESNLERLIAGLAKNASTPSLFGKGAAGAAPDTVYGVALCRGDLPNASACGDCVAGASRVARRACPLAEDVVVADDAGCQLRFS
>LOC_Os11g45540.1_DUF26-A
CGSSKYTANSIYQSNLDSLLSSSFLVVSGDSSSGALFAKGSRGAAPDTVYAVALCRGDANASACSGCVDAAYAAATARLCPLSKDAAVFYDECALRFSD
>LOC_Os12g41270.1_DUF26-A
CSSWNNFSVGSGYQVNLFKLLGNLAAGGAAAGSGGFYSGSYGALSDMVFGVAMCYVDRHWTKCRRCLDAATSGAAAFCPYSRRVDVMYDECVLRYSD
>LOC_Os12g41410.1_DUF26-A
CSTTGNYSGDSQYKKNLDQLLSTLATAATDDGWFNTSSVGTGGDDQVFGLIMCYADRNPTQCKECLAGAPAGITQVCPGSRTVNANYDACLLRYSD
>LOC_Os12g41490.1_DUF26-A
CSTAGNYTGDSQYKKNLDQLFTTLSAGAIAGDWFNTSSVGTGADQVFGLIMCYADRNSTQCQECLAGAPAGIVQVCPGSRTADANYDACLLRYSD
>LOC_Os12g41530.1_DUF26-A
CSTTGNYTGDSQYKKNLDQLFTSLSGGAIAGDWFNTSSVGTGADQVFGLIMCYADSNATECQKCLAMAPAVVQHPCRGSRSVNANYDACLLRYSD
>AFP1_DUF26-A
CSGSSYAGSSKAVANINSVLADLVASASSTGGYATSTAGKGNNSIIYGLAQCRGDVSASDCASCLADAAKQLPSTCSYSSDARIWYDYCFMRYEN
>SiCBMIP_DUF26-A
GAQTQASISQVLAALVPRASAAYYATATAGSGSSAIWGLAQCRGDIPASDCALCISAAAKQAASSCRGQADVRLWYDYCFLRYTD

### Amino acid sequences of DUF26-B

>AT1G04520.1_DUF26-B
AGTGFEERRDTAFGVMQNGVVSGHGFYATTYESVYVLGQCEGDVGDTDCSGCVKNALEKAQVECGSSISGQIYLHKCFIAYSY
>AT1G19090.1_DUF26-B
GEFWRFLDEALVNVTLKAVKNGGFGAASVIKTEAVYALAQCWQTLDENTCRECLVNARSSLRACDGHEARAFFTGCYLKYST
>AT1G61750.1_DUF26-B
FINTVEYRMDRLIQEAYSSSSYFAEETYHVSYLGEVYDLNGLVQCTPDLNQYDCYRCLKSAYNETKDCCYGKRFALVYSSNCMLTYK
>AT1G63550.1_DUF26-B
NPTRFNQTLTEKFSELIFNVSSSSLVPYFVEDQERVTQSEGSYDLDTMVQCSPDLDIFNCTVCLRVAFFRISTCCGLPSYAKIFTPKCLLRFQT
>AT1G63570.1_DUF26-B
NSNRFNQTLSNKLDQLIPNVSPSTLIPYFVEDQERVTQLEGSYDLVSMIQCSPDLDPSNCTICLRFAYATVSTCCGVPSSALIFTPKCILRYR
>AT1G63580.1_DUF26-B
YNQTLPGKLDELILKAPSSFSSPVPYFVEDKEHVTQVEGSYDLEAMAQCSPDLDPSSCTVCLGLVVEKFSECCSQSRWARIHFPKCLLRYD
>AT1G63590.1_DUF26-B
FSQTLLEKLDALILRASLSSSLPVPYFVDDQQHVTQLEGSYDLHAMVQCSPDLDPRNCTVCLRLAVQRLSGCCSHAQFARIFYTKCLITYE
>AT1G63600.1_DUF26-B
PNLDRFPQTLSDKMDELIINATSSPSLSSTPYFVEDQERVKQFEGSFDIDSMAQCSPDLDPRNCTTCLKLAVQEMLECCNQSRWAQIFTPKCLLRYEA
>AT1G70520.1_DUF26-B
CGNTTRKNKTFGDAVRQGLRNAVTEASGTGGYARASAKAGESESESAFVLANCWRTLSPDSCKQCLENASASVVKGCLPWSEGRALHTGCFLRYSDQ
>AT1G70530.1_DUF26-B
NRTVFRDNAAELVKNMSVEAVRNGGFYAGFVDRHNVTVHGLAQCWETLNRSGCVECLSKASVRIGSCLVNEEGRVLSAGCYMRFST
>AT1G70690.1_DUF26-B
RTGGNGNVQGVAQCSGDLSTSQCQDCLSDAIGRLKSDCGMAQGGYVYLSKCYARFS
>AT2G01660.1_DUF26-B
YRVGVSGELQGVAQCTGDLSATECQDCLMEAIGRLRTDCGGAAWGDVYLAKCYARYSA
>AT2G31620.1_DUF26-B
NKKLFNKNTKALLDKLKEKAIRKEQEPYTRDYMYAAGEESLGTTKLYGMMQCTQDLSVKNCSVCLDSIIAKLPRCCNGKQGGRVLNPSCTFRYELY
>AT2G33330.1_DUF26-B
NVAGTGFEQRRDTAFGVMQNGVVQGHGFYATTYESVYVLGQCEGDIGDSDCSGCIKNALQRAQVECGSSISGQIYLHKCFVGYSF
>AT3G21960.1_DUF26-B
DKKSFYKNMKAFLHKLKAKASSKENKPYVKDYMYAAGRESLGTVKLYAMVQCTQDLSLKNCTVCLDWIMTKLPECCNGKQGGRVLSPSCNF
>AT3G21970.1_DUF26-B
NVTNDPKRFEDKRRDLLHKLMLEATKDSKENGAKGLLYAVGEMRIGRNKMYAMVQCTQDLWQTGCHVCLEWITQMKYGEFFYRKPGGRVCGRSCSFRYELY
>AT3G21980.1_DUF26-B
NDKELFNKETSALLEELTNKATDKNNMIGNKFVLYAAGDKRIGTKNVYAMVQCTKDLVTTTSAACFEWIFKMFSKCCEGKQGGRVLGTSCNFRYELY
>AT3G21990.1_DUF26-B
KNMSDRELFNKETSALLEKLANKASDRNNLDGKQLVLYAAGEKRIGTKKVSAMVQCTKDLIFTKCFECLEGILRKFPECCDGKQGGRVLGTSCNFRYELY
>AT3G22040.1_DUF26-B
GDTKMFSKKTNDFLQQLIVKADKPDMDGVELLYYAAGEMRIGREKLHAMVQCAKDLADCKSCLEWSFKELSKCCDGKRGARFLGTSCNLRYELY
>AT3G22060.1_DUF26-B
NVSDPSTFNSQTKALLTELTKKATTRDNQKLFATGEKNIGKNKLYGLVQCTRDLKSITCKACLNGIIGELPNCCDGKEGGRVVGGSCNFRYEI
>AT3G29040.1_DUF26-B
KNVSSNAEQFKNKRKDLFHKLLLGATKDVSDSNDAYAVGETRIGRNKMYAMMQCALDLTTNGCYVCLEWIIGRYDSFYFDRRQGTRVLSRSCSLRYELY
>AT3G45860.1_DUF26-B
NQNQVDEFRDLVSSTLNLAAVEAANSSKKFYTRKVITPQPLYLLVQCTPDLTRQDCLRCLQKSIKGMSLYRIGGRFFYPSCNSRYENY
>AT3G60720.1_DUF26-B
CSSKSVENDYDFFKRRDDVLSDLESTQLGYKVSRSGLVEGYAQCVGDLSPSDCTACLAESVGKLKNLCGSAVAAEVYLAQCYARY
>AT4G00970.1_DUF26-B
DRNEFIRLQSELLNRLRSMAASGGSKRKYAQGTDPGSPPYTTFFGAVQCTPDLSEKDCNDCLSYGFSNATKGRVGIRWFCPSCNFQIE
>AT4G04490.1_DUF26-B
QEFAARANRTVEVASTADESSVLKYYGVSSAEFTDTPEVNMLMQCTPDLSSSDCNHCLRENVRYNQEHNWDRVGGTVARPSCYFRWDDY
>AT4G04500.1_DUF26-B
NMAMFSQEWIAMVNRTLEAASTAENSSVLKYYSATRTEFTQISDVYALMQCVPDLSPGNCKRCLRECVNDFQKQFWGRQGGGVSRPSCYFRWDL
>AT4G04510.1_DUF26-B
TLFRQEWEAMVDRTLEAVTIDNSTTVLKYYGALKSEFSEFPNVYMMMQCTPDINSGACKRCLQASVTYFRDQNWGRQGGGICRPSCVFRWEF
>AT4G04540.1_DUF26-B
KNMTLFEQEWNAMANRTVESATEAETSSVLKYYSAEKAEFTEFPNVYMLMQCTPDITSQDCKTCLGECVTLFKEQVWGRQGGEVYRPSCFFRWDL
>AT4G04570.1_DUF26-B
TLFKQQWEEMVNRTLEAATKAEGSSVLKYYKAEKAGFTEFPDVYMLMQCTPDLSSRDCKQCLGDCVMYFRKDYMGRKGGMASLPSCYFRWDL
>AT4G05200.1_DUF26-B
NITENQVSRFNESLPALLIDVAVKAALSSRKFATEKANFTVFQTIYSLVQCTPDLTNQDCESCLRQVINYLPRCCDRSVGGRVIAPSCSFRYELY
>AT4G11460.1_DUF26-B
NLTDFDRLWERLIAHMVTKASSASIKYLSFDNSRFYAADETNLTNSQMVYALMQCTPDVSPSNCNTCLKQSVDDYVGCCHGKQGGYVYRPSCIFRWDL
>AT4G11470.1_DUF26-B
AYTRTWDAFMNFMFTRVGQTRYLADISPRINQEPLSPDLIYALMQCIPGISSEDCETCLGKCVDDYQSCCNGFIGGVVNKPVCYFRWD
>AT4G11480.1_DUF26-B
AYTRTWEEFMNSMITRVGRTRYLADISPRIGSARIYALMQCIRGISSMECETCIRDNVRMYQSCCNGFIGGTIRKPVCFFRWD

>AT4G11490.1_DUF26-B
IWEALTDRLMSDASSDYNASLSSRRYYAANVTNLTNFQNIYALMLCTPDLEKGACHNCLEKAVSEYGNLRMQRGIVAWPSCCFRWDL
>AT4G11521.1_DUF26-B
FDRIWDELMSRTITTASRTHGSLSFGHKYYAADVASLTTFQTIYTMVQCTPDVSSGDCEFCLKRTVLDYKKCCRGHIGGAFVRPFCFIRWDL
>AT4G11530.1_DUF26-B
TVFDRIWEELMLRTITAASLSSSNGSSFGQKYFAAEVASLTTFQTMYAMMQCTPDVSSKDCEFCLKTSVGDYESCCRGKQGGAVIRPSCFVRWDL
>AT4G20550.1_DUF26-B
NNMSDRGLFNKETSALLEKLAYKASDRNNLDGKQLVLYAAGEKRIGTKKVYAMVQCTKDLIFTKCFECLEGILRKFPQCCDGKRGGRVFGTSCNFR
>AT4G20560.1_DUF26-B
NNMSDRGLFNKETSALLEKLAYKASDRNNLDGKQLVLYAAGEKRIGTKKVYAMVQCTKDLIFTKCFECLEGILRKFPQCCDGKRGGRVFGTSCNFR
>AT4G20570.1_DUF26-B
NNMSDRGLFNKETSALLEKLAYKASDRNNLDGKQLVLYAAGEKRIGTKKVYAMVQCTKDLIFTKCFECLEGILRKFPQCCDGKRGGRVFGTSCNFR
>AT4G20580.1_DUF26-B
NNMSDRGLFNKETSALLEKLAYKASDRNNLDGKQLVLYAAGEKRIGTKKVYAMVQCTKDLIFTKCFECLEGILRKFPQCCDGKRGGRVFGTSCNFR
>AT4G20590.1_DUF26-B
NNMSDRGLFNKETSALLEKLAYKASDRNNLDGKQLVLYAAGEKRIGTKKVYAMVQCTKDLIFTKCFECLEGILRKFPQCCDGKRGGRVFGTSCNFR
>AT4G20600.1_DUF26-B
NNMSDRGLFNKETSALLEKLAYKASDRNNLDGKQLVLYAAGEKRIGTKKVYAMVQCTKDLIFTKCFECLEGILRKFPQCCDGKRGGRVFGTSCNFR
>AT4G20610.1_DUF26-B
NNMSDRGLFNKETSALLEKLAYKASDRNNLDGKQLVLYAAGEKRIGTKKVYAMVQCTKDLIFTKCFECLEGILRKFPQCCDGKRGGRVFGTSCNFR
>AT4G20620.1_DUF26-B
NNMSDRGLFNKETSALLEKLAYKASDRNNLDGKQLVLYAAGEKRIGTKKVYAMVQCTKDLIFTKCFECLEGILRKFPQCCDGKRGGRVFGTSCNFR
>AT4G20640.1_DUF26-B
NNMSDRGLFNKETSALLEKLAYKASDRNNLDGKQLVLYAAGEKRIGTKKVYAMVQCTKDLIFTKCFECLEGILRKFPQCCDGKRGGRVFGTSCNFR
>AT4G20670.1_DUF26-B
NNMSDRGLFNKETSALLEKLAYKASDRNNLDGKQLVLYAAGEKRIGTKKVYAMVQCTKDLIFTKCFECLEGILRKFPQCCDGKRGGRVFGTSCNFR
>AT4G21230.1_DUF26-B
SVTDKEGFSKGLGDLLDSLGAKIDAANETKEVKFAAGVKGTIYALAQCTPDLSESDCRICLAQIFAGVPTCCDGKTGGWWTNPSCYFRFEV
>AT4G21400.1_DUF26-B
ISANRDEFDRLQIELLDRLKGIAAAGGPNRKYAQGSGSGVAGYPQFYGSAHCTPDLSEQDCNDCLVFGFEKIPGCCAGQVGLRWFFPSCSYRFET
>AT4G21410.1_DUF26-B
NRDDFERLQRGLLDRLKGIAAAGGPNRKYAQGNGSASAGYRRFYGTVQCTPDLSEQDCNDCLVFGFENIPSCCDAEIGLRWFSPSCNFRFET
>AT4G23130.1_DUF26-B
TADQSDRFNDAVLSLMKKSAEEAANSTSKKFAVKKSDFSSSQSLYASVQCIPDLTSEDCVMCLQQSIKELYFNKVGGRFLVPSCNSRYEV
>AT4G23140.1_DUF26-B
KQIDGFTSFVSSTMSEAAGKAANSSRKLYTVNTELTAYQNLYGLLQCTPDLTRADCLSCLQSSINGMALSRIGARLYWPSCTARYELY
>AT4G23150.1_DUF26-B
QFTNLVLSNMNQIAIEAADNPRKFSTIKTELTALQTFYGLVQCTPDLSRQNCMNCLTSSINRMPFSRIGARQFWPSCNSRYELY
>AT4G23170.1_DUF26-B
SQINQFIVLVQSNMNQAAMEAANSSRKFSTIKTELTELQTLYGLVQCTPDLTSQDCLRCLTRSINRMPLSRIGARQFWPSCNSRYELY
>AT4G23180.1_DUF26-B
DLLSDLVLPTLNQAATVALNSSKKFGTRKNNFTALQSFYGLVQCTPDLTRQDCSRCLQLVINQIPTDRIGARIINPSCTSRYEI
>AT4G23190.1_DUF26-B
TEFTKIWEGLMGRMISAASTAKSTPSSSDNHYSADSAVLTPLLNIYALMQCTPDLSSGDCENCLRQSAIDYQSCCSQKRGGVVMRPSCFLRWDL
>AT4G23200.1_DUF26-B
YKTNETEFNTVWSRLTQRMVQEASSSTDATWSGAKYYTADVAALPDSQTLYAMMQCTPDLSPAECNLCLTESVVNYQSCCLGRQGGSIVRLSCAFRAEL
>AT4G23210.1_DUF26-B
LTEFKTIWEDLTSRTITAASAARSTPSSSDNHYRVDFANLTKFQNIYALMQCTPDISSDECNNCLQRGVLEYQSCCGNNTGGYVMRPICFFRWQ
>AT4G23260.1_DUF26-B
NTNQTVFDIEWNNLTSSMIAGITSSSSGGNNSSKYYSDDIALVPDFKNISALMQCTPDVSSEDCNTCLRQNVVDYDNCCRGHQGGVMSRPNCFFRWEV
>AT4G23270.1_DUF26-B
IDLNASQSLYGMVRCTPDLREQDCLDCLKIGINQVTYDKIGGRILLPSCASRYDNY
>AT4G23280.1_DUF26-B
KRFAVTKFDLNALQSLYGMVQCTPDLTEQDCLDCLQQSINQVTYDKIGGRTFLPSCTSRYDNY
>AT4G23300.1_DUF26-B
TDFKNIWEDLTSRTITAASAARSTPSSSDNHYRVDFANLTKFQNIYALMQCTPDISSDECNNCLQRGVLEYQSCCGNNTGGYVMRPICFFRWQ
>AT4G28670.1_DUF26-B
KEKSAEFKGLVKEVTKSIVEAAPYSRGFSVAKMGIRDLTVYGLGVCWRTLNDELCKLCLADGALSVTSCLPSKEGFALNAGCYLRYSNY
>AT4G38830.1_DUF26-B
NFTGDRDSWEKSLRGLLEGLKNRASVIGRSKKNFVVGETSGPSFQTLFGLVQCTPDISEEDCSYCLSQGIAKIPSCCDMKMGSYVMSPSCMLAY
>AT5G37660.1_DUF26-B
YFRAGGSGDVQGMGQCVGDLTVSECQDCLGTAIGRLKNDCGTAVFGDMFLAKCYARYST
>AT5G40380.1_DUF26-B
FGVAGENGVHALAQCWESLGKEDCRVCLEKAVKEVKRCVSRREGRAMNTGCYLRYSD
>AT5G41290.1_DUF26-B
LESIVQCTPDLDPRNCTTCLKLALQELTECCGNQVWAFIYTPNCMVSFDTY
>AT5G41300.1_DUF26-B
SYDLDSLVQCSPHLNPENCTICLEYALQEIIDCCSDKFWAMIFTPNCFVNY

>AT5G43980.1_DUF26-B
QYESVYVLGQCEGSLGNSDCGECVKDGFEKAKSECGESNSGQVYLQKCFVSYSY
>AT5G48540.1_DUF26-B
PKKFDNELGALFDKIRSEAVLPKNKGLGKGKTKLTPFVTLNGLVQCTRDLSELDCAQCFATAVGSFMTTCHNKKGCRVLYSSCYVRYEF
>EFJ11427_DUF26-B
LAIFQSNSRTALEFLKAKVPRNARKFVAAVAGIGSAVPAYALGQCIPDLSAADCASCLAAAENAIASTCRGGGGGVCYYSSCTLFFQ
>EFJ16458_DUF26-B
IASSPGYDDNLKSALATLSRRGSSFTATVGSGKSQVFVLGECRGDLSSQQCDSCVRVAMRNMTDRCIGNTSGSILLNSCYTRF
>EFJ17937_DUF26-B
IASSPGYDDNLKSALATLSRRGSSFTATVGSGKSQVFVLGECRGDLSSQQCDSCVRVAMRNMTDRCIGNTSGSILLNSCYTRF
>LOC_Os01g23970.1_DUF26-B
AAARFMAKATELMNRTADLAAFGSSSSPSRYATGETWFDEQGVSVVYGLVQCTPDLTGEQCRSCLAGIIAQMPKLFGDASSRPVGGRILGVRCNLRYEK
>LOC_Os01g26390.1_DUF26-B
LLLFTLLNATAESAASSSRRFTTSRLDVSSLPTLYCLMQCTPDLTAGECAACFEDFPRLTLQYLDGARGGRILATRCTMRYEI
>LOC_Os01g36790.1_DUF26-B
NYTGANPRGFADAVRAALANVTGVAASAAVPGGGDGYAVGSASAGGATAFALAQCWGSLNATACGQCLRAAAAAAARCAPAAAEGRALYTGCYLRYST
>LOC_Os01g38850.1_DUF26-B
FGDRVMELINTTAEFAAWNSSKRGYATGEAGFGELDVGATRLGLVEQQCRSSPDLVIFALVQCTPDLSPAGCLSCLSGIASQMPRWFTGAADYRLGGRILGVRCNL
RYE
>LOC_Os02g43000.1_DUF26-B
CSSSTSRDGAFLSSRDGVLGELQAAAGYKLSTSGTVQGVAQCLGDVPANDCTACLAEAVGQLKGACGTALAADVYLAQCYVRY
>LOC_Os02g50200.1_DUF26-B
GSGGVQAMSQCVGDLGAKACSDCVSAAAGQLKAGCGYATAGEVYLGKCYARF
>LOC_Os03g16950.1_DUF26-B
DNGKAFQKAVGKVMGKATSQASQAGSGGLGRTKDQYTPFINIYGLAQCTQDLSPLACAQCLSTAVSRFGQYCGAQQGCQINYSSCRVRYEI
>LOC_Os03g16960.1_DUF26-B
DNAKAFQKAVGKVMSKATAQVSQAGSGGLGRVKDQYTPFINIYGFAQCTRDLSPLTCAQCLSTAVSRFDQYCGAQQGCRILYSSCMVRYEI
>LOC_Os03g36650.1_DUF26-B
REVSRLMKRLTRTAYLSPLLFAAGEAVAVGGAQRLHGMAQCTKDLSGGDCKMCLESAIDQLLARGCAKEGGKVLGGSCSLRYDF
>LOC_Os04g09780.1_DUF26-B
GDAAQFGAALSRLMDRLALAAASSSSSSRGRRFAFGQTNITGDGGDSLYAFVQCVDDLSPDDCRRCLQSIAASLPMTRGGRAYSLTCYTRFE
>LOC_Os04g25060.1_DUF26-B
AAYDRAVTELLAATVRYAVEENPARLFATGQRVGDDARDPGFRNIYSMAQCSPDLPPASCRRCLDGVLARWWQVFPLNGEGARVAGARCYLRSE
>LOC_Os04g25650.1_DUF26-B
AAYDRAVTELLAATVRYAVEENPARLFATGQRVGDDARDPGFRNIYSMAQCSPDLPPASCRRCLDGVLARWWQVFPLNGEGARVAGARCYLRSE
>LOC_Os04g45460.1_DUF26-B
CSTSTSGDGDFLKNRDAVLAALQGGLANGYKVSSSGNVQGVSQCLGDLAAGDCTTCLAQAVGQLKGTCGTSLAADVYLAQCYVRY
>LOC_Os04g56430.1_DUF26-B
ADVAAYDRAVTRLLAATAEYAAGDIARKLFATGQRVGADPGFPNLYATAQCAFDITLEACRGCLEGLVARWWDTFPANVDGARIAGPRCLLRSE
>LOC_Os05g02200.1_DUF26-B
QNATDPEAFEKAQRKVMARVAADAGDAGGGGLARETARFKDGVTIYGLGWCTRDITAADCGLCVAQAVAEMPNYCRFRRGCRVLYSSCMARYETY
>LOC_Os05g03920.1_DUF26-B
CSNATVSSPASFAATSAALLRNVTAAAPGARDYYYYSASSSSSASALPSVSPRVYAAAQCWRSLNATACAACVATARDRVVGRCLPRAAEGYGLNAGCVVRYSTQ
>LOC_Os05g41370.1_DUF26-B
DNATQPERFKSLLGTLMGNLTDAAARASSPLMFAAGETDLPPFTKIYGMAQCTRDLAAGDCYRCLVGAVNNIPKCCDGKQGGQVITRSCSIRFEV
>LOC_Os06g14280.1_DUF26-B
GAAGYVQAMSQCVGDLGAKACTDCVSAASSQLKAGCGYASAGEVYLGKCYARF
>LOC_Os07g30410.1_DUF26-B
YTTVRMDVVTPPLFSLMQCTPDMSGGDCRQCLQDLVGNTTFNGSVSGVRNIGARCGYRYDTY
>LOC_Os07g30510.1_DUF26-B
YTTVRMDVVTPPLFSLMQCTPDMSGGDCRQCLQDLVGNTTFNGSVSGVRNIGARCGYRYDTY
>LOC_Os07g34980.1_DUF26-B
RAQFSQLFSELMEKIAAAVVSRRPVNYLTGRGWFDLKSQTVYALAQCTDGMPPENCRSCLDGIIDEGKKMVGGGLTGGAVLGMRCSLWYQT
>LOC_Os07g35004.1_DUF26-B
VAAFDALVAELVNAVADRASNATRRYAAGKAGFAPEAMTVYAIAQCTPDLSPPQCRGCLAGIIDQMPKWFSGRVGGRILGVRCDFRYEK
>LOC_Os07g35140.1_DUF26-B
TTRSFFLSLVGTLFGEMAMYGSYNSSARRYASAVMYVNPQLPTVYGLAQCTPDLSPAQCWHCFQGLQEQNRQWYDGRQGGRILGVRCNFRYESY
>LOC_Os07g35280.1_DUF26-B
APAEVFDAAVVALLNATADHAAASSPRRFATGVEAFRGWGVRDIYALVQCTPDMSPAGCRSCLAGIISWVNDPDYFSGSPTGRVLGVRCNYWYD
>LOC_Os07g35290.1_DUF26-B
NASATAEVFDAAAATLLNATSGYAAANSSRRFATGEEAFDAADPTIYGLSQCTPDMSPDDCRSCLGGIIALIPQYFGRKRGARVIGTRCNYRYEV
>LOC_Os07g35300.1_DUF26-B
NASAPAEVFDAAVATLLNATSSYAAENSSRRFATGEEAFDAAAATPTIYGLSQCTPDMSPDDCRSCLGRIIALIPRYLSRRKGGRAIGMRCNFRYE
>LOC_Os07g35310.1_DUF26-B
SDPGRFNGMVAALVNATADYAAHNSTRRYASGEAVLDRESEFPKVYSWAQCTPDLTPAQCGDCLAAIIAKLPRLFTNRIGGRVLGVRCSYRYE
>LOC_Os07g35330.1_DUF26-B
FNAAVVALMNATVDTAVAAGSGSNNTKKYFATAVEDFDPKHYPKIYGMAQCAPVMTAAQCRSCLGGFVSSIPWFLNGKPGGRVLGIWCNLRYS

>LOC_Os07g35340.1_DUF26-B
VGWFNAAIAKILAALVDHAVATATGNNSTTKKYFATGEEDFDPNIYGFAQCVPDLTQEQCKECLNTFLFQAKQVYFGKSLSWVGMNSVWCRLMYS
>LOC_Os07g35380.1_DUF26-B
FNAAVTKILAAMVDHAVTSTTGNSTTKKYFVTGEEEFDPNIYGFAQCVPDLTPAQCNDCLKDLLFYAKQAYLGKSLSWVRVNSVWCRLMYS
>LOC_Os07g35390.1_DUF26-B
ATTEQARFNRLVAALVNATADYAARNSTRRRYASGEADFNAEFPKVYSWAQCTPDLTPASCRSCLAQIIGTYIGFFENRVGGFVRAVWCSFQYST
>LOC_Os07g35410.1_DUF26-B
WFNAAVAKILAALVEHTWATTTSNATAKKYFSTGEEEFNPKIYGFVQCVPDLSPEQCKECVRTLHDQAKIHYMGNSLPWASTYSVWCSLMYS
>LOC_Os07g35540.1_DUF26-B
WFNAAVAKILAALVEHAWATTTTTTTGNNSTTTIKYFATGEESFNPKIYGFAQCVPVLTPEQCKECLRSLHDNAKTVYMGNSLRWVGIYSVWCRLMYS
>LOC_Os07g35580.1_DUF26-B
GRYDRAVTGLLNATARYAAGNTNASSRLFATGVMVGFDAQFPKIYAMAQCSPDLSPAQCGLCLGAMVARWWQTFEPNTQGARSVGARCNMRVEL
>LOC_Os07g35640.1_DUF26-B
DYASAVYDAFSGMLVNATADYAAKDSVRRFGTGEMGFNVFDSPYHNIFSLAQCTPDMSEADCRSCLGDIIRRMMPKYFVGKPGGRVFGVRCNFRFEAY
>LOC_Os07g35650.1_DUF26-B
AAFDAAVAMLANATAEYAAAANTSRRYGTAEEEGVDGDGDSGRPRMYALAQCTPDKAADVCRACLTTLTTVQLPKLYSGGRTGGGVFGVWCNLRYE
>LOC_Os07g35660.1_DUF26-B
AAAFDAASGRLVNATAGYAAADPVRRFGTGEVGFDDATYPRIFSLAQCTPDLSEADCRSCLGRIIRWVPQYFAGKPGGRVFGVRCNFRFESY
>LOC_Os07g35690.1_DUF26-B
AAAAFDAAVAVLVNATADYAAADSSRRYGTGEEEGVDGDSDRPKIYALAQCTPDKTPEVCRTCLSTVIGQLPKEFSGRTGGGMFGVWCNFRYE
>LOC_Os07g35700.1_DUF26-B
PAFEAAVVRLINTTADYAATDSVRRFGTGEEAFDETTFPKIYSLAQCTPDMAATACRSCLEDIVGRMVSGNLIGRMGGRVLGVRCNLWFEV
>LOC_Os07g35740.1_DUF26-B
DVAGYDRAVTELLSATLMYAVVNTTRLFATGQRVGADPGFPNIYSAAQCTPDLSPALCRSCLEDLVARWWKTFPRTTVGARIVGTRCSLRSE
>LOC_Os07g35750.1_DUF26-B
ADVRAYDAAVVSLLNATVRYAVENSTRMFATGQRVGSDPGFSDIYSMAQCSPALSRPLCRSCLDGLVGQWWDTFPVNVEGARIAGTRCNLRSE
>LOC_Os07g35790.1_DUF26-B
VAGYDRAVTELLNATVRYAVENSTRLFATGQRVGADPGFRNIYSMAQCSPDLSPAQCRSCLDGLVGQWWTGFLFPRNGEGARVAGPRCYLRSE
>LOC_Os07g35810.1_DUF26-B
TDVAGYDRAVTELLNATVRYAVENSTKLFATGQRVGNDTGFSNIYSMAQCSPDLSPAQCRSCLDGLVGQWWKTFPLNGKGARVAGPRCYLRSE
>LOC_Os07g43560.1_DUF26-B
SDVGEFNGAIYEVLNATADYTAAAARRFGTGEISFDPTYPVIYSMAWCTPDMAPGRCRACLADTIAQMHAYFNPNAQGARLVGVRCAARYEI
>LOC_Os07g43570.1_DUF26-B
AAGRFYRLVGELLDATADYAVANSTARFATGDVGVGGYFDGEPFSKIYALAQCTPDLTPAQCRACLASAMEEMTRQVFAASSPGGKVIGERCGLRFE
>LOC_Os07g47230.1_DUF26-B
AGRFREHVAALLNATARDAAAQPDRYGTGDSWFQEGGSMVYALVQCTRDMDPGRCGACLQRIISEMPRMLDASQIGGRVLGVRCLLRYE
>LOC_Os08g04210.1_DUF26-B
KAFQKAAGKAMGKATAQAVAVGSSGLGRAKEQYTPFVSVYALAQCTRDLSPPSCAQCLSAAVSKFDKACGSGPGCQIDYSSCWARYEI
>LOC_Os08g04230.1_DUF26-B
FVNVYALAQCTRDLAPPLCAQCLSTTVSKFAEACGSGQGCQIDYSSCWVRYEI
>LOC_Os08g04240.1_DUF26-B
DNPKAFQKAAGKAMGKATAQAVAVGRSGLGRAKEQYTPFVSVYALAQCTRDLAPPACARCLSEIVSKFDKTCNSAQGCQIDYSSCWARYEI
>LOC_Os08g04250.1_DUF26-B
KAFQKVVGKAMVKATTQAVSVGGNGLGRAKEQYTPFVSVYALAQCTRDLAPPACAQCLSSTVSKFDKACGAAQGCQIDYSSCWARYEI
>LOC_Os10g04720.1_DUF26-B
ANVRELLTVTARTAAAAARRFATGFMDGSSESKQTLYSLAQCTPDLAAGDCLACLQRLIAMVNSTTSVRLGGRVLLLRCNLRFE
>LOC_Os10g04730.1_DUF26-B
QQLLSETVERAAGAAGRFATGVVDTGRTFPLVYSLAQCTPDLSAGDCLACLRRLTGMINSTMAVRMGAQIHVTRCYFRYEAY
>LOC_Os10g17950.1_DUF26-B
AGHFDALVADLAGALADWAAYNSTLRYAAGVMTSGDGFMSTTEDMVHNIYGVVQCTPDQAAAACRACLEALRVDMPKVFAGKMGGRFNAVWCNLRYET
>LOC_Os10g17960.1_DUF26-B
AGRFDALVARLAGALADWAAYNSTRRYAAGLMASGDGFTSTTEDMVHNIHGVVQCTPDQAAAACRACLETLRVDMPKVFAGRIGGRFDAVWCNLRYET
>LOC_Os11g28104.1_DUF26-B
ADGSFAGAARGLVGDLAAAAPRAPGLAAAAARGGVYAAAQCVETVGEGGCAQCLAVPARNIDGCPPDSDGRAVDAGCFMRYSD
>LOC_Os11g38850.1_DUF26-B
LVQETARTAAYNSSPPPPATTTYYATGRMDVSATFPTLYSMAQCTPDLRPGGCWRCLQSINDMTTRYFAGRRGGRILGLWCNFRYETY
>LOC_Os11g45540.1_DUF26-B
FTQFFIKTMNYIVAQALSTTKHYAAIRVDMDDADASNTVTLPRRLFCLAQCAPDLVEDICYNCLQNFSDLATANFAGRQGGRILALRCNLRYDT
>LOC_Os12g41270.1_DUF26-B
YTDSRGESLTVYGMVQCGRGLLPEECSKCLRHQLGELTTGLPNNTAGIIRGYSCYSRYD
>LOC_Os12g41410.1_DUF26-B
ETRWQLMSQLAETAGQTKLRLDTGSTRLGSTSMMYGLAQCTRDLAVSECSTCLSDYIVQLSKIFPNNSWAAIKGYSCYLRYD
>LOC_Os12g41490.1_DUF26-B
TMNDTRRRLMSQLAERAGDTKLRLDNGSLPYADSKLGTSALYGLAQCTRDLAASECRRCLSGYVDDLSNTFPNNSGGAIKGYSCYLRYH
>LOC_Os12g41530.1_DUF26-B
RWQLMSQLAERAGDTKLRLDNGSLPYVDSKLGTSALYGLAQCTRDLAASECRRCLSGYVNDLSNTFPNNSGGAIKGYSCYLRYQ
>AFP1_DUF26-B
DNPKAFEKAVGKVMGKATAQASAAGSAGLGRDKEQYTPFVSIYGLAQCTRDLAPLTCAQCLSTALSRFGDYCGAQQGCQINYSSCRVRYEI

>SiCBMIP_DUF26-B
NASDPAAFDRAERKLMARVAAEAGDAASGGLVRETARFGSATTIYGLGWCTRDITAADCGLCVAQAVAELPNYCQFRRGCRVLYSSCMARYETY
>AT3G04370.1_DUF26-B
NYSNQNLFLRAQALSSFLRKLESESSRSKFLKTLVGNEKHAVSGWFQCREDYPSEICHKCVGDLREISSRSCGNATSARIHLRGCHLIYK
>AT3G22030.1_DUF26-B
GNVNSFNKKTTEFLYKLIGKADRLDVDGINFLYYAAGEMRLGKQTLFAMVQCAKDILSCKDCLEWSIKELSKCCDGKQGARVVGTICNLRYELY
>AT4G20530.1_DUF26-B
KCSNTQGKYKQGSAFEKNLNLVLSTITSIGNFRDGFRYTEEGEDPNNVFVMFQCRGDSYWSKCPPCISTAVSGLRRRCPRNKGAIIWYDQCLLKIS
>AT4G20540.1_DUF26-B
KCSNTQGKYKQGSAFEKNLNLVLSTITSIGNFRDGFRYTEEGEDPNNVFVMFQCRGDSYWSKCPPCISTAVSGLRRRCPRNKGAIIWYDQCLLKIS
>AT4G20630.1_DUF26-B
KCSNTQGKYKQGSAFEKNLNLVLSTITSIGNFRDGFRYTEEGEDPNNVFVMFQCRGDSYWSKCPPCISTAVSGLRRRCPRNKGAIIWYDQCLLKIS
>AT4G20650.1_DUF26-B
KCSNTQGKYKQGSAFEKNLNLVLSTITSIGNFRDGFRYTEEGEDPNNVFVMFQCRGDSYWSKCPPCISTAVSGLRRRCPRNKGAIIWYDQCLLKIS
>AT5G41280.1_DUF26-B
CNDSSGNFTRNTTYNTNLNTLLSTLSNQSSFANYYNLTTGLGSDTVHGMFLCIGDVNRTTCNACVKNATIEIAKNCTNHREAIIYYFSCMVRYSD
>EFJ04200_DUF26-B
NVNSAMVVLSSHGSSTTVTSGDGKSRVFGLRECRRDMSAEQCNTCMAVATKNLHRETCYARFDTFL
>EFJ10327_DUF26-B
SFSPGSAFERNLDAALQSVISSSRSSPSAALGASPDVAYARGECYNNLSPQDCVLCLGMANQTIRGQAPRTIGARLFSNSSDYSCYLRYENY
>EFJ12758_DUF26-B
NTSLGSPFQQNWLDVYQVLLDHAPQAVSSEFSHGQSPNTVSGFALCIKGSNCKGCLQTIKANLDRAAPLSKGARLCMKTSTDVCYMRYEAY
>EFJ12964_DUF26-B
ESSEDSPFEKNLASAYESLLSPSQESPRQAQAGDDPDTVYGYAACFEGNCEDCLATCKNYFTQVAPHAVGARLCLKSGSDACYLRYENY
>EFJ13641_DUF26-B
RTYARGSVYGSNLNLLFDRLITTTGYDHVGSGSNRVYGFSECYGSPNCRSCLLAAVNSIRSAAPRAIGARICKALCYLRYENY
>EFJ13924_DUF26-B
NTSLGSPFQQNWLDVYQVLLDHAPQAVSSEFSHGQSPNTVSGFALCIKGSNCKGCLQTIKANLDRAAPLSKGARLCMKTSTDVCYMRYEAY
>EFJ15639_DUF26-B
STPYATNQNRLLDILKDTGSDSFIYTSVGDAPYIAYGRAECYGNCGNCLVEARNTIKNAVPQAVGARLCTDDCYLRFENY
>EFJ15855_DUF26-B
NTTEGSPFQKNVETVLNFFYEHNPPNGFATCLGYGDFPNTVYGHTGCYNMSDQPSCSECIKKAYQGIQQNAPYTLGGTYWINETEIRCRLRYENY
>EFJ16456_DUF26-B
NVNSAMVVLSSHGSSTTVTSGDGKSRVFGLRECRRDMSAEQCNTCMAVATKNLHRETCYARFDTFL
>EFJ18708_DUF26-B
KTTEGSPFQKNVETVLKFFYEHNPPNGFATCLGYGDFPNTVYGHTGCYNMSDQPSCSECIKRAYQGIQQYALYALGGTYWINETESRCRLRYENY
>EFJ22561_DUF26-B
ESSEDSPFEKNLASAYESLLSPSQESPRQAQVGDDPDTVYGYATCFEGNCEDCLATCKNYFTQVAPHAVGARLCLKSGSDACYLRYENY
>EFJ22969_DUF26-B
CSDESIPTGSTYWTSLNQTLTDLIQNTAYANNMTYSVKNGGDDVSAYGQAQCFNMVGNPGMCKSCVEDLVRRAWRECGDAIAGMLMLKDKCTIRYQD
>EFJ28936_DUF26-B
RTYARGSVYGSNLNLLFDRLITTTGYDHVGSGSNRVYGFSECYGSSNCRSCLLAAVNSIRSAAPRAIGARICKALCYLRYENY
>EFJ30128_DUF26-B
RKTTQGSQFQKNVETVIQFLYEHNPPNGFASCLGYGDFPDTAYGHTGCYNMSDQTSCASCIQQTYSEFQQLAPLAVGGVLWTNDTSNRCRIRFENY
>EFJ30567_DUF26-B
CSDESIPTGSTYWTSLNQTLTDLIQNTAYANNMTYSVKNGGDDVSAYGQAQCFNMVGNPGMCKSCVEDLVRRAWRECGDAIAAMLMLKDKCTIRYQD
>EFJ32980_DUF26-B
NTTEGSPFQKNVETVLNFFYEHNPPNGFATCLGYGYFPNTVYGHTGCYNMSDQPSCSECIKKAYQGIQQNAPYTLGGTYWINETEIRCRLRYENY
>EFJ33692_DUF26-B
RNTQPGSAFQRNLSNVYRRIINGATFRQGSISYNAGVDPDRVYGFSACFGGFTAPGDCVQCLVDIRANFDSVAPLAVGARMCSRNGERTCYLRYENY
>EFJ33923_DUF26-B
LYSFGDPFGKNLGLVLDDLLGTTPGRDFKDYTAISPRPSRFVYGHAKCRHALSAGECGVCLKGVAAALADQCGNAVGGRAMFVDCFMRYEAY
>EFJ37042_DUF26-B
RNTQPGSAFQRNLSNVYRRIINGATFRQGSISYNAGVDPDRVYGFSACFGGFTAPGDCVQCLVDIRANFDSVAPLAVGARMCSKNGERTCYMRYENY
>EFJ37434_DUF26-B
LYSFGDPFGKNLGLVLDDLLGTTPRRDFKDYTAISPRPFRFVYGHAKCRHALSAGECGVCLKGVAAALADQCGNAVGGRAMFVDCFMRYEAY
>LOC_Os01g36500.1_DUF26-B
CGASNYTADSMYRLNLDGMSASLFPEGAGGSGGGIFVRGSSGADPDKVYAVALCRGDVDDAPACSSCFDAAFRRAMQLCPRSKDATIYYDECLLRFSD
>LOC_Os01g38910.1_DUF26-B
CSLSGGRYGQNTTYEDNLKALAARLVGVARVSNFASHTVGSAPDAAYGIALCRGDYTGDECANGLRKAFENAVENRLFCDRFRDATIYYDQYMLRFS
>LOC_Os01g56720.1_DUF26-B
GRVVAQCAAGVAPADCVQCLEGAAREMPRCFREARREEQGEGVGIVVSDDCVLRFD
>LOC_Os01g56840.1_DUF26-B
TGSNGTAFRANLLTLLASLPDQAAPTGFASMQAGAGGRAPGGDDDDRAFARGSCLGDSTPSQCRDCLAAAVIDVAEGCGADTRRAGAWLSGCYLAYAD
>LOC_Os03g19840.1_DUF26-B
NGKTRRRSINSVVSDLVAKAASNGGFATSSAGKGNNVFYGLAQCRGDVSASDCKACLVEAANYTLSFCHYASDSRMWYDYCFMRYKN
>LOC_Os07g35190.1_DUF26-B
YAANSTYEANLRYLAATLPAKVMNGSSSSSVDVLAGERPNLIAASASCNSSSSEYHDCGACVAEAFRCARRLCPYSRHAVVHLGGGACSVRYY

>LOC_Os10g10540.1_DUF26-B
CTSTAAGNYTQDGAYAANLGRLLAMLPNETVSKNGGFFNGTVGNGTATVYGLAMCAADFSRADCMDCLVAAGISAGGVVKRCPGSTTVSAMFDQCLLRYSD
>LOC_Os12g41510.1_DUF26-B
CSTTGNYTVGNQFEKNLDQLLSTLATAATDDGWFNTSSVGTGTAYQVFGLIMCHADYNATECKKCLAGAPAGIKQVCPGSRTVKANYDACLLRYSD
>PTQ30062_DUF26-B
RNGTAYQHNVRSVLSKLMAKVQSRPLFDVWDEASDAYDNSETAYGVAGCNTVTPEECRYCLQYIVTNLQQWCPKDPLSAKIVLKDCYLHYDRY
>PTQ32071_DUF26-B
AVSACIDSLTEEPMDKKTASAEVKTPDKVVYCSMDCKQNLTLQECQACLQTAKLLRENYCPRAVSASSVIGICSLMHET
>Gnk2_DUF26-B
CNTQKIPSGSPFNRNLRAMLADLKQNTAFSGYDYKTSRAGSGGAPTAYGRATCKQSISQSDCTACLSNLVNRIFSICNNAIGARVQLVDCFIQYEQ
